# CrJAMs are transcriptional repressors of the monoterpenoid indole alkaloid pathway in *Catharanthus roseus*

**DOI:** 10.64898/2026.11.02.763677

**Authors:** Samuel Breselge, Krystyna K. F. Traverse, Lauren F. Cole-Osborn, Suphinya Sathitloetsakun, Kristina Dürr, Jie Wang, Kevin L. Childs, Erin J. Cram, Carolyn W.T. Lee-Parsons

## Abstract

*Catharanthus roseus* produces monoterpenoid indole alkaloids (MIAs) in response to the defense hormone jasmonate (JA). In *Arabidopsis*, JA regulates defense processes through MYC2 and JA-Associated MYC2a-like transcription factors (JAMs). Here we characterize two previously identified *C. roseus* JAMs, CrJAM2 and CrJAM3, as regulators of MIA biosynthesis.

Functional screens using viral-induced gene silencing and transient promoter activity assays showed that CrJAMs repress the MIA pathway; silencing both *CrJAM2* and *CrJAM3* increased expression of MIA pathway genes and MIA levels while their individual overexpression repressed promoter activity. To assess their activity broadly across MIA regulatory and biosynthetic genes, we generated *C. roseus* hairy root cultures either constitutively overexpressing individual *CrJAMs* or *CRISPR-Cas9* creating mutations in multiple *CrJAMs*, and profiled their expression by RNA-Seq. *CrJAM2* overexpression decreased expression of most MIA biosynthetic genes relative to the *GUS* control, while double mutant lines showed increased basal expression of these genes relative to non-targeting controls. Notably, methyl JA treatment perturbs this equilibrium, reversing these trends at 24 hrs in both *CrJAM* overexpression and mutant lines relative to basal conditions.

In summary, CrJAMs broadly repress MIA regulatory and biosynthetic gene expression and together with CrJAZ and CrMYC2a regulates MIA biosynthesis.

**Summary sentence:** We characterize two previously identified *Catharanthus roseus* JAMs; CrJAM2 and CrJAM3 regulate overlapping and unique genes and broadly repress the expression of monoterpenoid indole alkaloid regulatory and biosynthetic genes.

## 1. Introduction

*Catharanthus roseus* produces over 130 different alkaloids (Van der Heijden et al., 2004), including the valuable chemotherapeutics, vinblastine and vincristine (Noble et al., 1958; Aslam et al., 2010; Škubník et al., 2021). Vinblastine and vincristine are produced from the condensation of precursors from the monoterpenoid indole alkaloid (MIA) pathway; this defense-associated pathway is induced by jasmonates (JAs; Aerts et al., 1994), hormones that act in various developmental and stress responses (Wasternack and Strnad, 2018).

Many molecular components of the JA signaling pathway are conserved between the model organism *Arabidopsis thaliana* and other plant species (Thaler et al., 2012). For instance, *A. thaliana* and *C. roseus* share the early steps of JA perception and regulation, e.g. the role of JAZ (Jasmonate ZIM-domain) and MYC2 (Zhang et al., 2011; Patra et al., 2018). In addition, a class of JA-induced basic helix-loop-helix (bHLH) repressors in *A thaliana,* known as JAMs (Jasmonate-associated MYC2-like transcription factors), were identified in *C. roseus* (Patra et al., 2018; Schweizer et al., 2018) and are characterized in this paper.

MYCs and JAMs are members of the bHLH transcription factor (TFs) family which play a key role in mediating JA responses. In *A. thaliana*, bHLH subgroup IIIe contains key activators of JA-responses, *AtMYC2*, *AtMYC3*, *AtMYC4*, and *AtMYC5*, while subgroup IIId contains repressors of JA-responses*, AtbHLH17* (*AtJAM1*), *AtbHLH13* (*AtJAM2*), *AtbHLH03* (*AtJAM3*), and *AtbHLH14* (Heim et al., 2003). In the absence of JAs, JAZ proteins bind and inhibit JA-responsive TFs like MYCs and JAMs (see top tier of Figure 1; reviewed in Chini et al., 2016; Li et al., 2024). In the presence of JAs, the SCF^COI1^-JAZ (Skp/Cullin/F-box complex with CORONATINE INSENSITIVE1 and JAZ) coreceptor complex forms, leading to the ubiquitination and degradation of JAZ proteins (Thines et al., 2007). The degradation of JAZ releases MYCs and JAMs. Released MYC2 binds to G-boxes (CACGTG and its variants) and activates JA-responsive gene expression (Lorenz et al., 2004; Dombrecht et al., 2007; Fonseca et al., 2014); MYC2 also activates the expression of *JAZ* and *JAM* (Chini et al., 2007; Sasaki-Sekimoto et al., 2013), regenerating JAZ. JAMs also bind to G-boxes and its variants but lack an activator domain; therefore, JAM proteins compete with MYC2 and attenuate JA-responses (Fonseca et al., 2014). For instance, AtMYCs activate while AtJAMs repress the expression of the senescence-associated protein *SAG29*, fine-tuning JA-induced leaf senescence (Qi et al., 2015). The competition of JAMs and MYCs for the same binding site appears to fine-tune defense and developmental responses.

**Figure 1.**
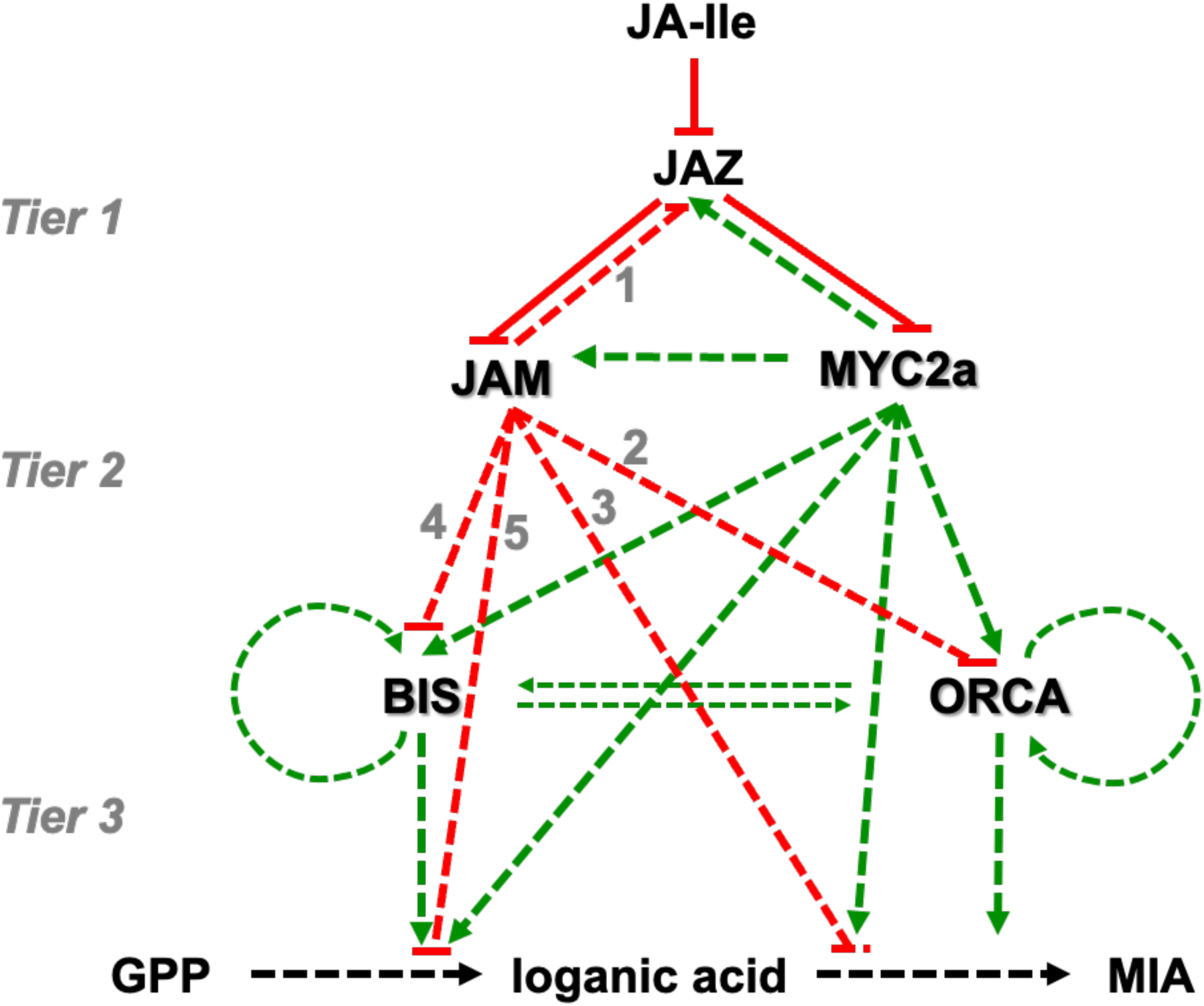
Model for the jasmonate-responsive regulation of MIA biosynthesis by JAZ, MYC2a, ORCA, and BIS previously proposed in the literature, overlaid with regulation by JAM from this paper The regulation of MIA biosynthesis by MYC2a, ORCA, and BIS in this figure was previously published in Schweizer et al., 2018. The regulation of MIA biosynthesis by JAMs is investigated and presented in this paper and overlaid on the figure adapted from Schweizer et al., 2018. Activation is indicated by green lines and green arrowheads whereas inhibition is indicated by red lines and red flat-head arrow. Solid lines indicate confirmed protein-protein interactions whereas dashed lines indicate either confirmed protein-DNA interactions or regulation based on overexpression of JAMs or MYC2a. JA-Ile: jasmonate-isoleucine conjugate; GPP: geranyl pyrophosphate. Please refer to the Introduction and the Discussion Sections for detailed descriptions of the regulatory relationships; the grey numbers indicate relationships inferred from the experimental results and summarized in the Discussion.

In *C. roseus*, additional components of the JA-responsive network regulating MIA biosynthesis have been identified (Figure 1). As in *A. thaliana*, CrJAZ proteins interact and repress CrMYC2a (Patra et al., 2018; Schweizer et al., 2018). Upon CrJAZ degradation, CrMYC2a (particularly the derepressed CrMYC2a^D126N^) activates MIA pathway gene expression and MIA accumulation (Zhang et al., 2011; Schweizer et al., 2018) by inducing the expression of key MIA pathway activators like CrBIS (bHLH Iridoid Synthesis) and CrORCA (Octadecanoid-Responsive Catharanthus AP2-domain) (see second tier of Figure 1; Van Der Fits et al., 2000; Zhang et al., 2011; Van Moerkercke et al., 2015, 2016; Paul et al., 2017; Schweizer et al., 2018; Colinas et al., 2021); CrMYC2a, CrBIS, and CrORCA then directly induce the expression of the MIA pathway genes (see third tier of Figure 1; Zhang et al., 2011; Schweizer et al., 2018). The proposed regulation of *CrBIS* and of the genes in the seco-iridoid pathway by CrMYC2a differ between Patra et al (2018) and Schweizer et al (2018), potentially due to the different tissue context studied. In addition to CrJAZs, other JA-induced repressors of MIA biosynthesis have been characterized (not shown in Figure 1), such as CrZCTs (zinc finger *Catharanthus* Transcription Factors) and CrGBFs (G-box Binding Factors) (Sibéril et al., 2001; Pauw et al., 2004; Schweizer et al., 2018). Of these repressors, CrZCTs have been studied most extensively (Pauw et al., 2004; Peebles et al., 2009; Goklany et al., 2013; Li et al., 2013; Rizvi et al., 2016; Traverse et al., 2024).

Here we characterize the JA-induced repressors, JAMs, in *C. roseus*. We hypothesize that CrJAMs attenuate JA-responses similarly as in *A. thaliana*. Using the *A. thaliana* JAM amino acid sequences, we identified only the previously reported three bHLH IIId subgroup members: CrJAM2, CrJAM3 (Schweizer et al., 2018), and Repressor of MYC2 Targets 1 (RMT1; Patra et al., 2018). Overexpression of *CrRMT1* represses CrMYC2 targets (*CrMYC2*, *CrORCA3*, and *CrSTR1*) and reduces MIAs (catharanthine, tabersonine, and ajmalicine) in hairy roots (Patra et al., 2018).

Here we focus on characterizing *CrJAM2* and *CrJAM3*. In an initial screen, virus-induced gene silencing (VIGS) of *CrJAM2* and *CrJAM3* together increased the expression of several MIA activators and pathway genes and increased specific MIAs. In addition, we demonstrated that CrJAM2 and CrJAM3 significantly repress the transcriptional promoter activity of several MIA activators and pathway genes in transiently transformed *C. roseus* seedlings. To evaluate the regulation of the entire MIA biosynthetic pathway, we developed hairy root cultures that overexpress either the individual *CrJAMs* or *CRISPR-Cas9* to create double or triple mutants and conducted a global RNA-Seq. These results show that JAMs broadly act as transcriptional repressors of the MIA regulatory and biosynthetic genes in *C. roseus*.

## 2. Methods

### Accession numbers

*A. thaliana bHLH IIId* genes: *AtbHLH03/AtJAM3* (*Arabidopsis* Genome Initiative number: AT4G16430), *AtbHLH13/AtJAM2* (AT1G01260), *AtbHLH14* (AT4G00870), *AtbHLH17/AtJAM1* (AT2G46510). *C. roseus bHLH IIId* genes: *CrJAM2* (GenBank: MG676662), *CrJAM3* (GenBank: MG676663), *CrRMT1* (GenBank: KY851107). *C. roseus* MIA pathway enzymes and TFs used to identify their promoters and 5’UTRs: *CrBIS1* (GenBank: KM409646), *CrBIS2* (GenBank: KM409645), *CrG8O* (also referred to as *CrG10H*, GenBank: AJ251269), *CrGO* (GenBank: MF770508), *CrLAMT* (GenBank: EU057974), *CrMYC2a* (GenBank: AF283507), *CrORCA3* (GenBank: AJ251249), *CrSTR1* (GenBank: Y10182), *CrT16H1* (GenBank: FJ647194), *CrT16H2* (GenBank: JF742645), *CrT19H* (GenBank: HQ901597), *CrZCT1* (GenBank: AJ632082).

### Identification of *C. roseus* bHLH proteins

To identify bHLH proteins in *C. roseus*, the bHLH domains of the following proteins were used in a BlastP search against the *C. roseus* genome version 3 (Li et al., 2023): CRO_07G000280 (CrMYC2a), CRO_08G020500 (CrBIS1), CRO_07G004820 (CrMYC1), CRO_04G030430 (a bHLH DNA-binding superfamily protein), CRO_05G017550 (a bHLH DNA-binding superfamily protein), AT1G18400.1 (AtBEE1), AT1G68240.1, AT2G46510.1 (AtJAM1), AT4G00480.1 (AtMYC1), AT1G32640.1 (AtMYC2), and AT1G09530.1 (AtPIF3). Hits with E-values ≤0.05 were included for further consideration. For each hit, the longest isoform sequence was used to generate a list of candidate proteins. Proteins were scanned using the Pfam-A v37.3 database and the HMMER3 package version 3.4 with domain specific E-value cutoffs in CLC Main Workbench 8.1.3 (QIAGEN). Hits were further analyzed for the presence of bHLH domains using several databases through InterPro (Mitchell et al., 2019) and the domain annotated using PROSITE (Sigrist et al., 2013) and aligned to identify the 5-9-13 motif (Heim et al., 2003; Toledo-Ortiz et al., 2003).

### Identification of *C. roseus* bHLH IIId proteins

Amino acid sequences of the four *A. thaliana* bHLH IIId proteins (AtbHLH17, AtbHLH13, AtbHLH3, and AtbHLH14) were used in a BlastP search with the *C. roseus* genome v3 (Li et al., 2023) in CLC Main Workbench. The best hits were used for a NCBI Blast search to identify if the sequences were reported previously.

### Construction of phylogenetic trees

Amino acid sequences were aligned in CLC Main Workbench (gap open cost = 10; gap extension cost = 1; end gap cost = as any other; alignment mode = very accurate). Identified domains were aligned as described above and manually curated. The amino acid alignment was used to create a maximum likelihood phylogenetic tree (Neighbor Joining; WAG; 100 replicates for full-length proteins and 1000 replicates for domains) in CLC Main Workbench.

### Hierarchical clustering of *C. roseus* RNA-Seq data

RNA-Seq reads were downloaded from NCBI-SRA databases (BioProject: PRJNA252611) and processed to obtain sequencing reads in FASTQ format. Reads were processed to trim the sequencing adapter sequences and read positions with low sequencing quality using Trimmomatics version 0.36 (Bolger et al., 2014). Clean reads were aligned to the *C. roseus* reference genome v2 using STAR version 2.5.2b (Dobin et al., 2013). Aligned sequences were counted using the featureCount version 1.22 package (Liao et al., 2014) in R to get number of reads aligned to a gene feature. Read counts data were normalized to get gene expression values in FPKM (fragments per kilo base million). Heatmap was generated using gplots package (Warnes et al., 2009) in R.

### Cloning of VIGS vectors

Fragments (400 bp) targeting *CrJAM2* or *CrJAM3* were designed using the Sol Genomics Network VIGS tool (Fernandez-Pozo et al., 2015) (n-mer = 23, mismatches = 0), which minimized off-targets in the *C. roseus* version 2 transcriptome (Franke et al., 2019). Fragments were amplified from previously cloned plasmids using Phusion High-Fidelity DNA Polymerase (New England Biolabs) with primers listed in Table S 3. Fragments were inserted, either individually or together, into pTRV2-GG (Addgene plasmid #105349) using modular cloning and confirmed with Sanger Sequencing. Final plasmids were confirmed with a restriction enzyme digest and then electroporated into *Agrobacterium tumefaciens* GV3101 (pMP90). Plasmids targeting *CHLH* and *GFP* for silencing were cloned previously (Cole-Osborn et al., 2024a).

### Virus-induced gene silencing (VIGS)

*C. roseus* Little Bright Eye (NESeeds) were sterilized, germinated, and finally cultivated in soil under the 16 hr light / 8 hr dark photoperiod until two true leaves appeared as described previously (Cole-Osborn et al., 2024a). Once seedlings had two true leaves, they were infected with *A. tumefaciens* according to the pinch-wounding method (Liscombe and O’Connor, 2011).

A single colony of *A. tumefaciens* GV3101 (pMP90) harboring pTRV1 or pTRV2 was used to inoculate LB with the appropriate antibiotics, cultivated, and prepared as described in Cole-Osborn et al (2024a). *A. tumefaciens* strains containing pTRV1 and pTRV2 plasmids were then combined in a 1:1 ratio (OD_600_ of each strain = 2-4). Modified tweezers were dipped into the *A. tumefaciens* solution and the plant was pinched three times in the highest internode beneath the shoot apical meristem.

After infection, plants were cultivated in the dark for two days before being placed back into a 16 hour light / 8 hour dark photoperiod as described previously (Cole-Osborn et al., 2024a). Plants were grown until two pairs of leaves emerged after silencing and the *CHLH*-silenced plant exhibited yellow leaves. At this point, the two youngest leaf pairs were harvested for RNA and alkaloid extraction.

### RNA extraction

Briefly, RNA was extracted from liquid nitrogen flash-frozen leaves or hairy root tissue; tissue was crushed by shaking in the Mini-BeadBeater-16, extracted with RNAzol-RT (Molecular Research Center) and the Direct-zol RNA Miniprep Plus Kit (Zymo Research) with on-column DNAse treatment to remove genomic DNA as described previously (Cole-Osborn et al., 2024a). For RNA-Seq experiments, an additional DNAse treatment was performed using the TURBO DNA-freeTM kit (Invitrogen), according to manufacturer’s instructions. RNA integrity was assessed using agarose gel electrophoresis, and concentration and purity were quantified with a NanoDrop (ND-1000 Spectrophotometer; ThermoScientific).

### Monitoring gene expression

cDNA was synthesized from extracted RNA using either the SuperScript II First-Strand Synthesis System (Invitrogen) or the LunaScript RT SuperMix Kit (New England Biolabs) with up to 2.5 µg of RNA, according to manufacturer’s instructions. cDNA was diluted 1:4, and 1 µL was used in a 10 µL reaction with SYBR Green ROX qPCR Master Mix (Qiagen or ABClonal) and 300 nM primers. Amplification efficiency for each primer set was confirmed using Ct values over a range of cDNA dilutions and was 100% ± 10% for each gene monitored. If primers did not show 100% efficiency, the concentrations of the forward and reverse primers were varied until 100% efficiency was achieved. Optimal concentrations of primers that differed from the standard 300 nM for both forward and reverse primers were as follows: JAM2q_set3F = 600 nM; JAM2q_set3R = 600 nM; JAM3q_set3F = 600 nM; JAM3q_set3R = 600 nM; Orca3-forward = 300 nM; Orca3-reverse = 600 nM. Specificity of each primer set was confirmed by gel electrophoresis and sequencing.

Reactions were run on the MX3000P (Agilent) or CFX96 (Bio-Rad) qPCR instrument using the thermocycler protocol as described previously (Cole-Osborn et al., 2024a). Transcript levels were normalized to the housekeeping gene, SAND (Pollier et al., 2014), and fold changes relative to the negative control condition were calculated according to the 2-ΔΔCt method (Livak and Schmittgen, 2001). SAND Ct values in no reverse-transcriptase controls were confirmed to be at least 5 Ct values above the respective experimental sample, indicating minimal genomic DNA contamination (Svec et al., 2015).

### Alkaloid extraction from leaf tissue

At the time of harvest, individual leaves were weighed to determine their fresh weight, flash-frozen in liquid nitrogen, crushed using a Mini-BeadBeater-16 (Biospec), and extracted in methanol, as described previously (Cole-Osborn et al., 2024a). Ajmaline (50 ng) per mg tissue served as an internal standard prior to extraction. After extraction, concentration, and centrifugation steps, the final extract was diluted 1:50 in 50% methanol/50% water.

### Quantification of alkaloid content in leaf tissue with HPLC-MS/MS

Alkaloid quantification was performed at the Mass Spectrometry Facility at Northeastern University, as described previously (Cole-Osborn et al., 2024a). Briefly, alkaloids were separated using the Thermo Scientific^TM^ Vanquish HPLC system with a Phenomenex Luna Omega LC column and detected on a Tandem HRMS orbitrap mass analyzer (Thermo Scientific Exploris 240) coupled to an electrospray ionization (H-ESI) source in the positive mode. Quantitative analysis was performed in the full scan MS with data-dependent tandem mass spectrometry. For normalization, peak areas of each compound were divided by the peak area of ajmaline in each sample.

### Vector construction

All vectors (except pSB310-312) were constructed using the Golden Gate system, MoClo (Weber et al., 2011). Vectors and parts were used from Weber et al., 2011, Engler et al., 2014, and Mortensen et al., 2019. All sequences from *C. roseus* (Little Bright Eye) gDNA or cDNA were amplified with Phusion High-Fidelity DNA Polymerase. All new promoters and coding sequences were confirmed by Sanger Sequencing. Correct assembly of transcriptional units (L1 vectors) and expression vectors (L2 vectors) were confirmed by partial sequencing, colony PCR, and/or restriction digest. All final vectors were assembled in the pSB90 vector backbone (Mortensen et al., 2019) containing a constitutively active *VirG* gene (*VirGN54D*) for increased *Agrobacterium* virulence (Pazour et al., 1992) and *KanR* as bacterial selection marker. Vector cartoons for the final vectors are given in Figure S 6.

### Cloning of *CrJAMs* and MIA pathway promoters

*CrJAM2* and *CrJAM3* coding sequences were amplified as fragments to remove and replace type IIS restriction enzyme recognition sites (*BpiI, Eco31I, Esp3I*) with synonymous codons by mutating an A to T or a T to A within the recognition site. Fragments were cloned into pAGM1311 (Weber et al., 2011) before combining in pICH41308 (Weber et al., 2011). Oligonucleotides used in cloning are given in Table S 3 and coding sequences of *CrJAM2* and *CrJAM3* are given in Table S 6.

The promoters and 5’UTRs of several MIA pathway transcription factors and pathway enzymes (including the MEP-secoiridoid pathway) were used to test if CrJAM2 and CrJAM3 can repress their activity. Promoter and 5’UTR sequences of the genes of interest were identified with Blast using the coding sequences (see section ‘Accession numbers’) against the *C. roseus* genome v2 (Franke et al., 2019; v3 was not available when constructs were cloned) in CLC Main Workbench. Up to 2 kb upstream of the translational start site was amplified from *C. roseus* Little Bright Eye gDNA and cloned into pICH41295 (Weber et al., 2011). *BpiI* or *Eco31I* sites were mutated as described above. In the process no E-boxes (CANNTG) were created or removed. Primers can be found in Table S 3 and promoter sequences are given in Table S 6.

There are two different gene locations with 100% exon identity for *CrGO* in the genome of the *C. roseus* Sunstorm Apricot variety (cro_v2_scaffold_43, Franke et al., 2019) but their 5’ sequences differ. The 5’ region from 324.327 to 325.969 in cro_v2_scaffold_43 was chosen to represent the *GO* promoter. The amplified *CrT16H2* 5’ sequence from *C. roseus* Little Bright Eye variety gDNA contained a 653 bp deletion compared to the expected sequence in the genome of the *C. roseus* Sunstorm Apricot variety (Franke et al., 2019).

### Promoter activity assays in *C. roseus* seedlings

To test their ability to repress the above-mentioned *C. roseus* promoters, *CrJAM2* and *CrJAM3* were transiently overexpressed in *C. roseus* seedlings using the efficient *Agrobacterium*-mediated seedling infiltration (EASI) method (Mortensen et al., 2019, 2022). Briefly, 10-day old *C. roseus* seedlings were vacuum-infiltrated with two strains of *A. tumefaciens* GV3101(pMP90). The first strain contained the effector (at OD_600_: 0.34), either *CaMV2×35S::GUS* (pSB161, serving as the control), *CaMV2×35S::CrJAM2* (pSB270), or *CaMV2×35S::CrJAM3* (pSB271). The second strain contained the promoter plus 5’UTR of MIA TF or enzyme driving the *FLUC* reporter (at OD_600_: 0.06) and the constitutive *AtNos-*driven *RLUC* reporter for normalizing transformation efficiency. Cartoons of vectors are given in Figure S 6.

Transformed seedlings were harvested 3 days post-infection and luciferase assays performed using the Luc-Pair^TM^ Duo-Luciferase HT Assay Kit (Genecopeia) as described in Mortensen et al. (2019). For the promoter of the gene of interest, the relative transcriptional promoter activity was calculated by dividing the ratio of FLUC (firefly luciferase) to RLUC (*Renilla* luciferase) luminescence of the effector condition (*JAM2* or *JAM3* overexpression) by the ratio of FLUC to RLUC luminescence of the *GUS* control. The experiment was carried out in three independent assays with 10 biological replicates per assay and each biological replicate or data point consisting of 2 seedlings (Figure 4, Figure S 7).

### Guide design and cloning for CRISPR-Cas9 deletion of *CrJAM2*, *CrJAM3*, and *CrRMT1*

Guide sequences targeting genomic regions of *CrJAM2*, *CrJAM3*, and *CrRMT1* were designed using Benchling.com and on-target scores were predicted (Doench et al., 2016). Guides with high on-target scores (>50) and low chances of off-targets (16/20 or less identity to closest other Blast hit in *C. roseus* genome v2) were chosen (Table S 5). Guides were cloned into the improved pDGE vector system. The pDGE vectors contain improvements in the sgRNA scaffolds and in the *Cas9* sequence and their demonstrated improvements are described in (Grützner et al., 2021).

### Development and maintenance of stable transgenic hairy roots

Transgenic hairy roots were developed and maintained as described previously (Rizvi et al., 2015; Traverse et al., 2022). For the overexpression lines, the *Agrobacterium rhizogenes* R1000 contained one of the following constructs: *CaMV2×35S::CrJAM2* (pSB297), *CaMV2×35S::CrJAM3* (pSB298), or *CaMV2×35S::GUS* (control line, pSB299, Figure S 6). For deletion lines*, A. rhizogenes* R1000 contained one of the following constructs: CRISPR-Cas9 vectors with sgRNAs targeting 1) *CrJAM2* and *CrJAM3* (pSB311), 2) *CrJAM2*, *CrJAM3,* and *CrRMT1* (pSB312), or 3) the control without sgRNAs (pSB310, Figure S 6).

### Confirmation of transgenic hairy root lines

Hairy root tissue was harvested and extracted for genomic DNA (GeneJET Plant Genomic DNA Purification Mini Kit; Thermo Scientific) and genes confirmed by PCR as described previously (Rizvi et al., 2015; Traverse et al., 2022). In addition, the presence of *GUS* (control line, pSB299), *CrJAM2* (pSB297), or *CrJAM3* (pSB298) in the overexpression lines was confirmed using primers shown in Table S 3.

For the deletion lines, the amplification of the sgRNA region or the Cas9 and promoter region in the control lines lacking guides was also confirmed using primers shown in Table S 4; to evaluate the edits, the genomic regions of *CrJAM2*, *CrJAM3*, and *CrRMT1* from the different hairy root lines were amplified from gDNA using primers in Table S 4 and gel-purified PCR products were sent for Sanger sequencing. PCR products with different mutations were cloned into pAGM1311 (Weber et al., 2011) and individual clones were sent for sequencing to distinguish different allelic mutations.

### Illumina RNA-Seq

Total RNA samples were submitted to the MSU Genomics Core for Illumina sequencing. Samples were checked for RNA integrity using an Agilent 2100 Bioanalyzer, and all samples had RIN scores better than 7.0. Illumina TruSeq Stranded mRNA libraries were prepared following the manufacturer’s protocol. Libraries were checked using Qubit dsDNA High Sensitivity and Agilent 4200 TapeStation High Sensitivity DNA1000 assays. The libraries were combined on an equimolar basis into multiple pools, which were quantified using a Kapa Biosystems Illumina Library Quantification qPCR kit. Library pools were each sequenced on one lane of an Illumina HiSeq 4000 flow cell using single end 50 bp chemistry. Demultiplexed FastQ files were prepared for each RNA sample. A minimum of 30 million reads was generated for each sample. FastQ files have been deposited in NCBI Sequence Read Archive (BioProject: PRJNA1001907).

### Analysis of differentially expressed genes in transgenic hairy root lines

Reads were prepared, aligned to the *C. roseus* genome, and differentially expressed genes were identified. All reads were trimmed of adapters and low-quality bases with Trimmomatic version 0.36 (Bolger et al., 2014). The *C. roseus* v2 genome and annotation files were downloaded from Dryad (https://doi.org/10.5061/dryad.08vv50n; Jakob et al 2018), and cleaned RNA-seq reads were aligned to the genome assembly using STAR version 2.5.2b (Dobin et al 2013). Only uniquely mapping reads were retained. The v2 *C. roseus* gene annotation was supplemented with missing ZCT1 and ZCT4 gene models. The featureCounts function from the Rsubread version 3 R package was used to assign read counts to all genes (Liao et al 2013). Read counts were normalized using the TMM function from the edgeR package (Robinson et al 2010), and genes with fewer than 5 counts in five or more samples were discarded. Count data were transformed with the voom function from the limma R package (Law et al 2014). A linear model (genotype + time + genotype:time) was used to fit the data with the lmfit function from limma, and genes with *p*-values less than 0.01 were identified as differentially expressed. Significantly differentially expressed genes were then filtered for those genes with log_2_-fold change greater than 2.

### Statistical analysis of RNA-Seq data

Fold change in gene expression of the two highest *CrJAM2* and two highest *CrJAM3*-overexpressing lines were calculated from their individual RPKM divided by the average RPKM of two *GUS*-overexpressing control lines for each time point and then transformed with log_2_. Fold change in gene expression of two individual *jam2jam3* and two individual *jam2jam3rmt1*-mutant lines were calculated from their individual RPKM divided by the average RPKM of four no-guide control lines for each time point and then transformed with log_2_. Fold changes were unable to be calculated for genes that were not expressed in the control condition. Heatmaps were constructed using GraphPad Prism.

## Results

### *C. roseus* genome encodes for three bHLH IIId JAM-like transcription factors

Shown in Figure S 1, we identified 105 bHLH proteins and three bHLH IIId TFs in the *C. roseus* genome version 3 (Li et al., 2023) as compared to 164 bHLH proteins and four bHLH IIId TFs in *A. thaliana* (Bailey et al., 2003; Heim et al., 2003; Hao et al., 2021; Qian et al., 2021). Together, the phylogenetic tree of all bHLH TFs from *C. roseus* (Figure S 1) and the BlastP results (Table S 1) suggest that the bHLH IIId subgroup consists of just three TFs in *C. roseus,* which were previously identified by Patra et al. (2018) and Schweizer et al. (2018): CrJAM2, CrJAM3, and CrRMT1.

We compared the bHLH domain of CrJAMs and CrMYC2a for their DNA recognition sequences (Figure 2). The characteristic bHLH domain consists of a N-terminal conserved sequence of <u>b</u>asic amino acids, followed by a conserved amphipathic α-<u>H</u>elix, a <u>L</u>oop of variable length (up to 14 amino acids and average of 6 amino acids), and a conserved second amphipathic α-<u>H</u>elix (Heim et al., 2003). The first α-helix domain, specifically the amino acid positions 5-9-13, influences the recognition of target DNA sequences by these TFs (Heim et al., 2003; Toledo-Ortiz et al., 2003); CrJAMs, CrMYCs, and CrBIS all contain the H-E-R amino acids in the 5-9-13 motif within the first α-helix domain (Figure S 1). Therefore, they are expected to bind to the same DNA sequence (G-box: CACGTG; Heim et al., 2003).

**Figure 2.**
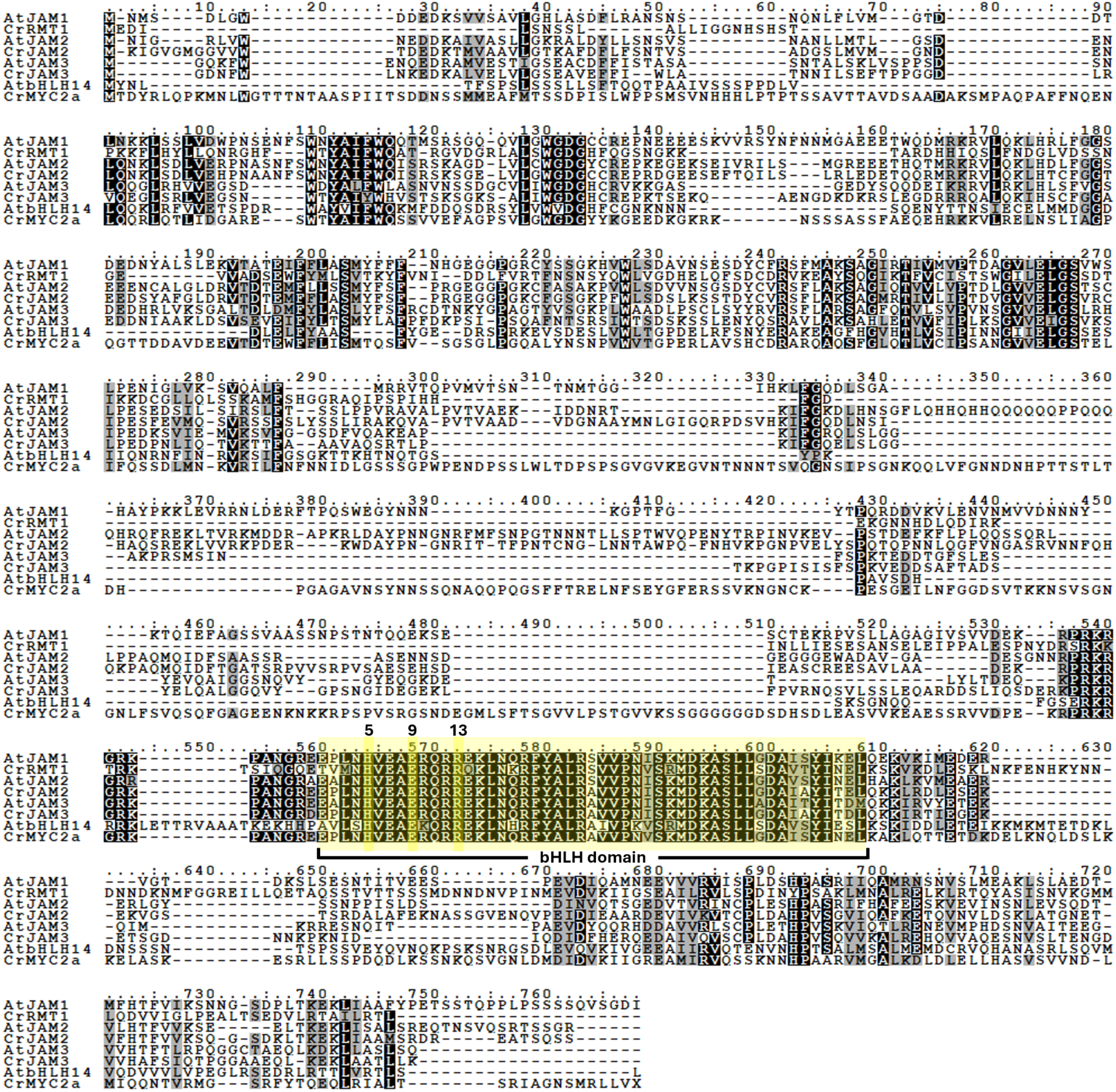
Protein alignment of JAMs from *A. thaliana* and *C. roseus*. Protein sequences of JAMs from *A. thaliana* and *C. roseus* were downloaded from the NCBI database: AtJAM1 (AT2G46510), CrRMT1(KY851107), AtJAM2 (AT1G01260), CrJAM2 (MG676662), AtJAM3 (AT4G16430), CrJAM3 (MG676663), AtbHLH14 (AT4G00870), CrMYC2a (CRO_07G000280). Protein sequences were aligned in CLC Main Workbench (Gap open cost = 10.0, Gap extension cost = 1.0, End gap cost = as any other) and shaded for homologous regions using Boxshade. Black shading indicates residues that are identical to the consensus while grey shading indicates residues that are similar to the consensus. The bHLH region (as predicted by InterPro) is highlighted in yellow and the 5-9-13 motif (amino acids H-E-R) is indicated above the alignment.

The domain responsible for the repressive activity of JAMs has not been clearly identified in the literature. In *Arabidopsis*, the homologs AtJAM2 and AtJAM3 lack an activation domain found in AtMYC2 and therefore are assumed to be passive repressors, competing for the same binding site with AtMYC2 (Fonseca et al., 2014). In contrast, Huang et al (2018) showed that the conserved N-terminal region of AtJAM2 is necessary for repression while the C-terminal region, which includes the bHLH domain, is insufficient for repression. Therefore, functional assays are needed to confirm that the candidate TF is a repressor.

The three bHLH IIId TFs cluster closely with the bHLH IIIe TFs, CrMYC2a and CrMYC2c (Figure S 1). CrRMT1 (Patra et al., 2018) and CrMYC2b (Schweizer et al., 2018) share ∼98% nucleotide (1311/1332 bp) and amino acid (434/443 aa) identity and the genes map to the same locus in the *C. roseus* v3 genome, suggesting that CrRMT1 and CrMYC2b refer to the same gene. The reported differences in the gene sequences may be due to single nucleotide polymorphisms between different *C. roseus* varieties. Of these three bHLH IIId TFs in *C. roseus,* CrRMT1 was previously characterized and shown to repress CrMYC2 targets (Patra et al., 2018). CrJAM2 (1812 bp) and CrJAM3 (1473 bp) were only tested previously for their ability to regulate and repress the *CrBIS1* promoter (Schweizer et al., 2018). Since CrRMT1 has been studied in detail (Patra et al., 2018), we mainly focused on characterizing CrJAM2 and CrJAM3 in this paper.

Previously published transcriptome data supported the JA-responsiveness of *CrJAM2* and *CrJAM3,* particularly in cell suspensions and/or sterile seedlings (Góngora-Castillo et al., 2012) (Figure S 2). Co-expression analysis of *CrJAM2* and *CrJAM3* with known MIA pathway enzymes and TFs showed that *CrJAM3* and *CrMYC2a* shared the most similar expression, while *CrJAM2* and *CrRMT1* shared the most similar expression (Figure S 3). Since a repression domain in JAMs has not been clearly identified, functional assays are necessary to clearly distinguish between the MYC2 and JAM proteins, which act as activators and repressors, respectively. CrMYC2a activity was previously characterized by Zhang et al. (2011) and Schweizer et al. (2018).

### Silencing *CrJAM2* and *CrJAM3* generally increased MIA pathway gene expression and MIA production

We silenced *CrJAM2* and *CrJAM3* in *C. roseus* leaves using virus-induced gene silencing (VIGS) and analyzed its effect on gene expression and MIA production. To ensure sufficient silencing of both *CrJAM2* and *CrJAM3*, we first tested the efficacy of silencing two genes at once and the order of the silencing fragments on overall silencing. Targeting both *CrJAM2* and *CrJAM3* in the same construct was just as efficient as targeting either *CrJAM2* or *CrJAM3* alone and the order of the silencing fragment when targeting both *CrJAM* genes also did not matter; all silencing constructs led to ∼50% silencing of the targeted gene(s) (Figure S 4).

Using the *JAM3/2* construct, significant silencing of *CrJAM2* and *CrJAM3* (by 36 and 62%, respectively) was confirmed in the second leaf pair to emerge after infection (immature leaf, Figure 3A), but not in the first leaf pair (mature leaf, not shown); thus, we only proceeded with analyzing the gene expression and the MIA content of the immature leaves.

**Figure 3.**
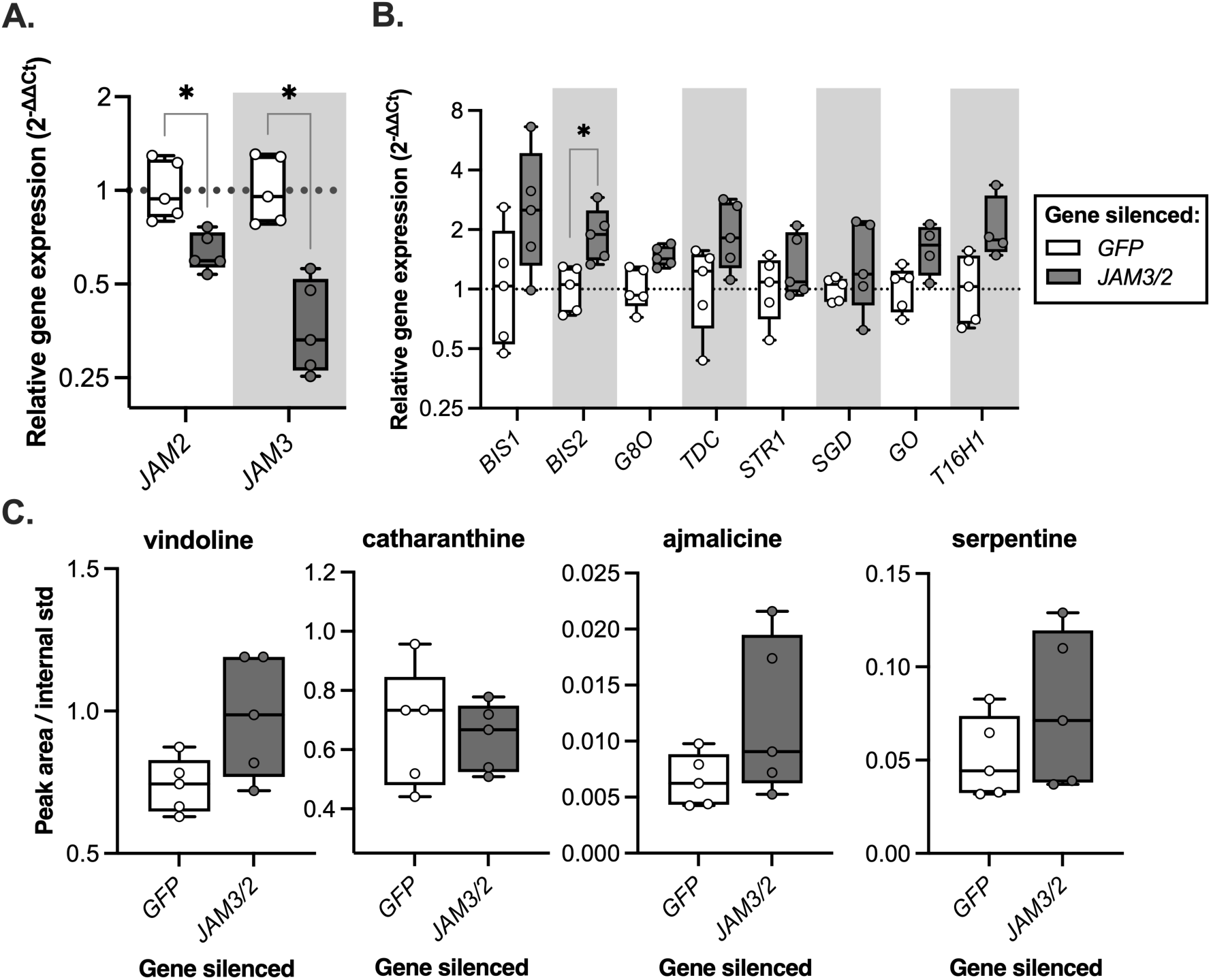
Virus-induced gene silencing of *CrJAM2* and *CrJAM3* in *C. roseus* leaves (indicated as *JAM3/2*) led to increases in MIA gene expression and alkaloid accumulation. **(A)** *CrJAM2* and *CrJAM3* were significantly silenced in *JAM3/2*-silenced plants. **(B)** Silencing *CrJAM2* and *CrJAM3* led to non-significant increases in the gene expression of most MIA TFs and MIA pathway genes. Relative gene expression was measured with qPCR and calculated using the 2^-ΔΔCt^ method (Livak and Schmittgen, 2001) relative to the control condition (*GFP*-silenced plants) and normalized relative to the housekeeping gene, *SAND* (Pollier et al., 2014). * q < 0.05 according to an unpaired two-tailed t-test on ΔΔCt values with a false discovery rate of 5% (two-stage linear step-up procedure of Benjamini, Krieger and Yekutieli, with 17 genes); vindoline pathway genes are shown in Figure S 5. **(C)** Alkaloid accumulation (vindoline, catharanthine, ajmalicine, and serpentine) increased in *JAM3/2*-silenced plants. Metabolite levels were quantified from the peak area of the extracted ion chromatogram using HPLC-MS/MS divided by the peak area of the internal standard, ajmaline. RNA and alkaloids were extracted from the second leaf pair to emerge after infection for each plant (n = 5). No statistical significance in alkaloid levels was found according to an unpaired two-tailed t-test on normalized peak areas with a false discovery rate of 5% (two-stage linear step-up procedure of Benjamini, Krieger and Yekutieli, with 4 metabolites). Boxes represent the 25th and 75th percentile with a line marking the median. Whiskers extend to the minimum and maximum.

We observed ∼2-fold or higher increases in *BIS1, BIS2, TDC,* and *T16H1* when *CrJAM2* and *CrJAM3* were both silenced (Figure 3B), supporting the role of CrJAMs as repressors of the MIA pathway. We could not monitor *ORCA3* as its expression was too low and variable to accurately quantify. We also observed non-significant increases in specific MIAs when both *CrJAM2* and *CrJAM3* were silenced in the immature leaves (33% increase in vindoline, 86% increase in ajmalicine, 51% increase in serpentine, no changes in catharanthine, Figure 3C). Given the observed increase in vindoline, we checked the expression of the other vindoline pathway genes. Several vindoline pathway genes increased by 40 - 60% while two genes decreased with *JAM3/2* silencing (Figure S 5).

The transient nature of VIGS yields partial silencing, leading to non-significant changes in gene expression and metabolite levels. However, our findings are consistent with the potential role of CrJAMs as repressors of MIA biosynthesis.

### CrJAM2 and CrJAM3 repressed the transcriptional promoter activity of several MIA pathway TFs and genes in transiently transformed *C. roseus* seedlings

In validating CrJAM2 and CrJAM3 as repressors of the MIA pathway, we cloned five promoters from MIA pathway regulators (*MYC2a*, *BIS1*, *BIS2*, *ORCA3*, and *ZCT1*) and seven promoters from MIA pathway enzymes (*G8O* or *G10H*, *LAMT*, *STR1*, *GO*, *T16H1*, *T16H2,* and the root-specific *T19H*) to drive the expression of the *FLUC* reporter gene (Table S 2; Table S 6; Figure S 6, Figure S 7). Either *CrJAM2* or *CrJAM3* was transiently co-expressed with these promoter-reporter constructs in *C. roseus* seedlings using the Efficient *Agrobacterium-*mediated Seedling Infiltration (EASI) method (Mortensen et al., 2019, 2022; Cole-Osborn et al., 2024b). If CrJAM repressed these promoters (directly or indirectly), luminescence would decrease relative to that of the overexpressed negative effector control, *GUS*.

The overexpression of either *CrJAM2* or *CrJAM3* significantly reduced the promoter activity of activators (*pBIS1, pBIS2,* and *pORCA3*) and MIA pathway enzymes (*pG8O*, *pGO*, and *pT16H1*) (Figure 4A). These results complement the known regulation of the genes by MYC2a^D126N^, BIS1/2, and ORCA3 (Figure 4b; Schweizer et al., 2018; Traverse et al., 2024). Since CrJAM, CrMYC2a, and CrBIS all bind to the same DNA sequence, promoters activated by CrMYC2a or CrMYC2a^D126N^ (*pBIS1, pBIS2, pORCA3, pZCT1, pG8O, pLAMT, pSTR1, pGO,* and *pT19H*) and by CrBIS (*pBIS2, pORCA3,* and *pG8O*) would also be expected to be repressed by CrJAMs; our results differed in that we did not observe a significant repression of *pZCT1, pLAMT, pSTR1,* and *pT19H*, observed repression of *pT16H1*, and observed unexpected activation of *pT16H2*. These differences might be attributed to the evaluation methods used (promoter driving reporter versus quantitative gene expression) and the tissue contexts used (seedlings versus flowers), resulting in difference in basal expression levels and activator levels that compete with the CrJAM2 and CrJAM3.

**Figure 4.**
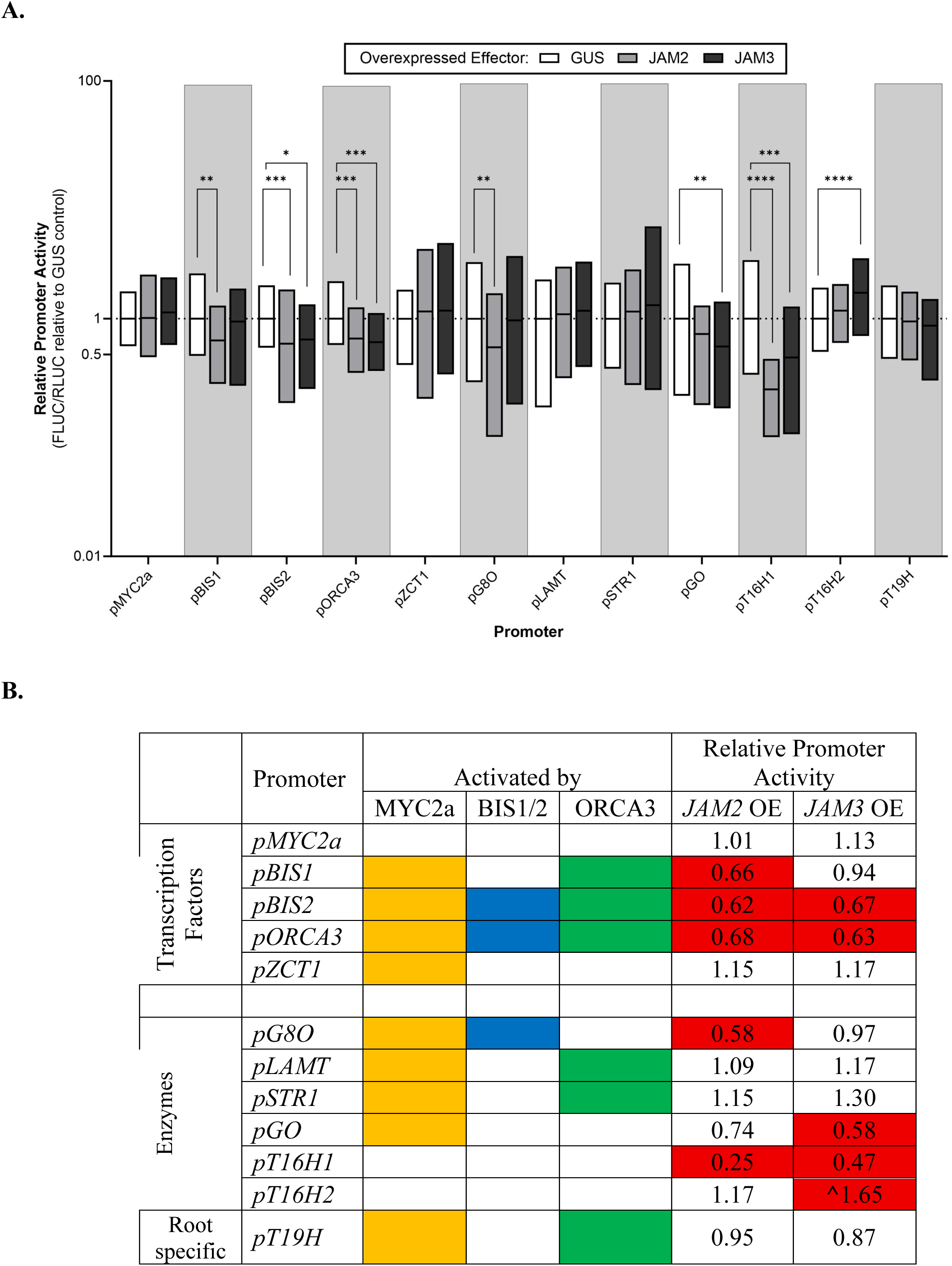
The overexpression of either *CrJAM2* or *CrJAM3* represses *C. roseus* promoters in transiently transformed *C. roseus* seedlings **(A)** *C. roseus* seedlings were vacuum-infiltrated with a combination of two strains of *A. tumefaciens* (total OD_600_ = 0.4): 1) strain containing the promoter and its 5’UTR driving the *FLUC-I* reporter (Figure S 6; OD_600_ = 0.06), and 2) strain containing a *CaMV2×35S* driven effector for transactivation (either *GUS* as the negative effector control – pSB161 (Addgene ID #123197), *CrJAM2* – pSB270, or *CrJAM3* – pSB271; OD_600_ = 0.34). Samples were harvested three days post-infection. The relative promoter activity was the ratio of FLUC to RLUC luminescence for each sample normalized to the average FLUC to RLUC luminescence of the *GUS* overexpression control (set to 1). The experiment was carried out with ten replicates per condition and three independent experiments were performed (n = 30 replicates per condition). The black horizontal line within the boxes shows the median, and the ends of the boxes show the 25^th^ and 75^th^ percentile. The data was log-normal transformed to obtain normally distributed data. Significant differences compared to the *GUS* control were determined using Dunnett’s method on log-normal transformed data. P-values <0.05 are indicated with one star (*), <0.01 are indicated with two stars (**), and <0.001 are indicated with three stars (***), and <0.0001 are indicated with four stars (****). **(B)** Statistical analysis was performed with the Dunnett’s method on log normal transformed data. Significant promoter activity changes (P-values <0.05) are highlighted in red (^ indicates activation rather than repression). Included in the table are genes known to be activated by MYC2a^D126N^, BIS1/2, ORCA3 based on findings reported in Schweizer et al. (2018) and Traverse et al., (2024).

Our transient assay showed that CrJAM2 and CrJAM3 significantly repressed the promoter activity of key activator TFs (*BIS1, BIS2,* and *ORCA3*) and several MIA pathway enzymes (*G8O*, *GO*, and *T16H1*), supporting their potential role as repressors of the MIA pathway.

### Overexpression of *CrJAM2* or *CrJAM3* in hairy root lines repressed MIA pathway gene expression

To evaluate the regulation of the entire MIA biosynthetic pathway, we generated hairy root lines that constitutively overexpressed either *GUS* (negative effector control), *CrJAM2*, or *CrJAM3* under the control of the strong *CaMV 2×35S* promoter and monitored the effect of their overexpression by RNA-Seq. Multiple independent lines were evaluated.

*CrJAM2* and *CrJAM3* overexpression lines resulted in up to a 23-fold and 17-fold increase of *CrJAM2* and *CrJAM3* expression, respectively (Figure S 8). Interestingly, the constitutive overexpression of *CrJAM2* led to an average reduction of *CrJAM3* expression by 27% while the overexpression of *CrJAM3* led to an average reduction of *CrJAM2* expression by 38%. Since MYC2a activates *JAM* expression and since MYC2a and JAMs bind to the same promoter target, the overexpression of one *JAM* would be expected to repress the expression of the other *JAM*. Feedback regulation of MIA TFs (like *CrJAMs*) was also observed with the overexpression of *CrMYC2a* (Patra et al., 2018), *CrBIS* (Schweizer et al., 2018), or *CrORCA* (Schweizer et al., 2018).

Next, the gene expression profile of the two strongest expressing *CrJAM2* (rSB02_297_07_2, rSB02_297_09_1) and the two strongest expressing *CrJAM3* (rSB02_298_32_1, rSB02_298_33_1) hairy root lines from Figure S 8 were assessed and are presented as a heat map in Figure 5. Root lines were elicited with methyl JA (250 μM MJ) and sampled at basal conditions (0 hr), 0.5 hr, and 24 hrs. The 0.5 hr time point after MJ addition monitors expression resulting from pre-existing TFs like MYC2a and JAMs released upon the degradation of the JAZ repressor, while the 24 hr time point after MJ addition monitors the expression from newly transcribed TFs (Goklany et al., 2013; Rizvi et al., 2016).

**Figure 5.**
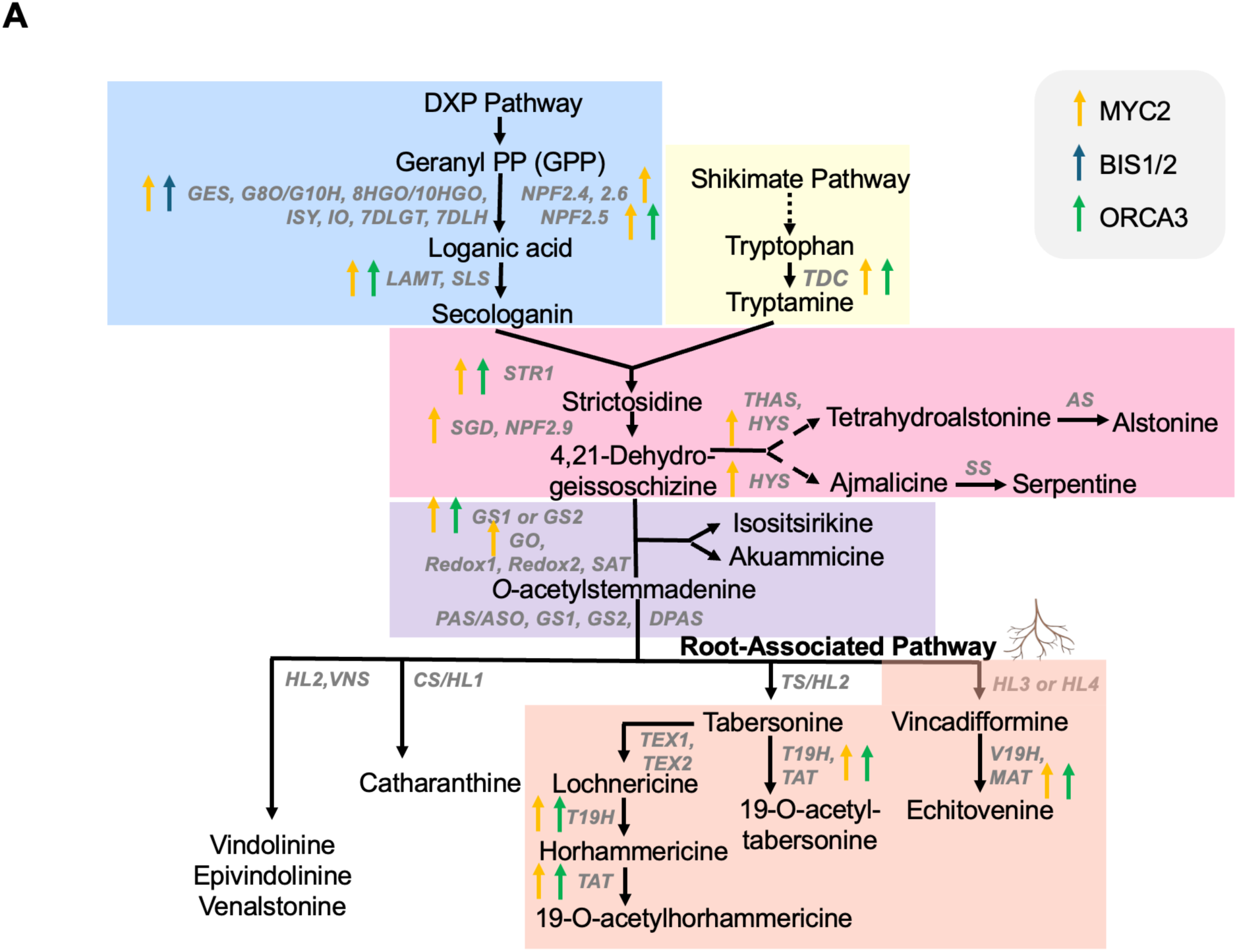
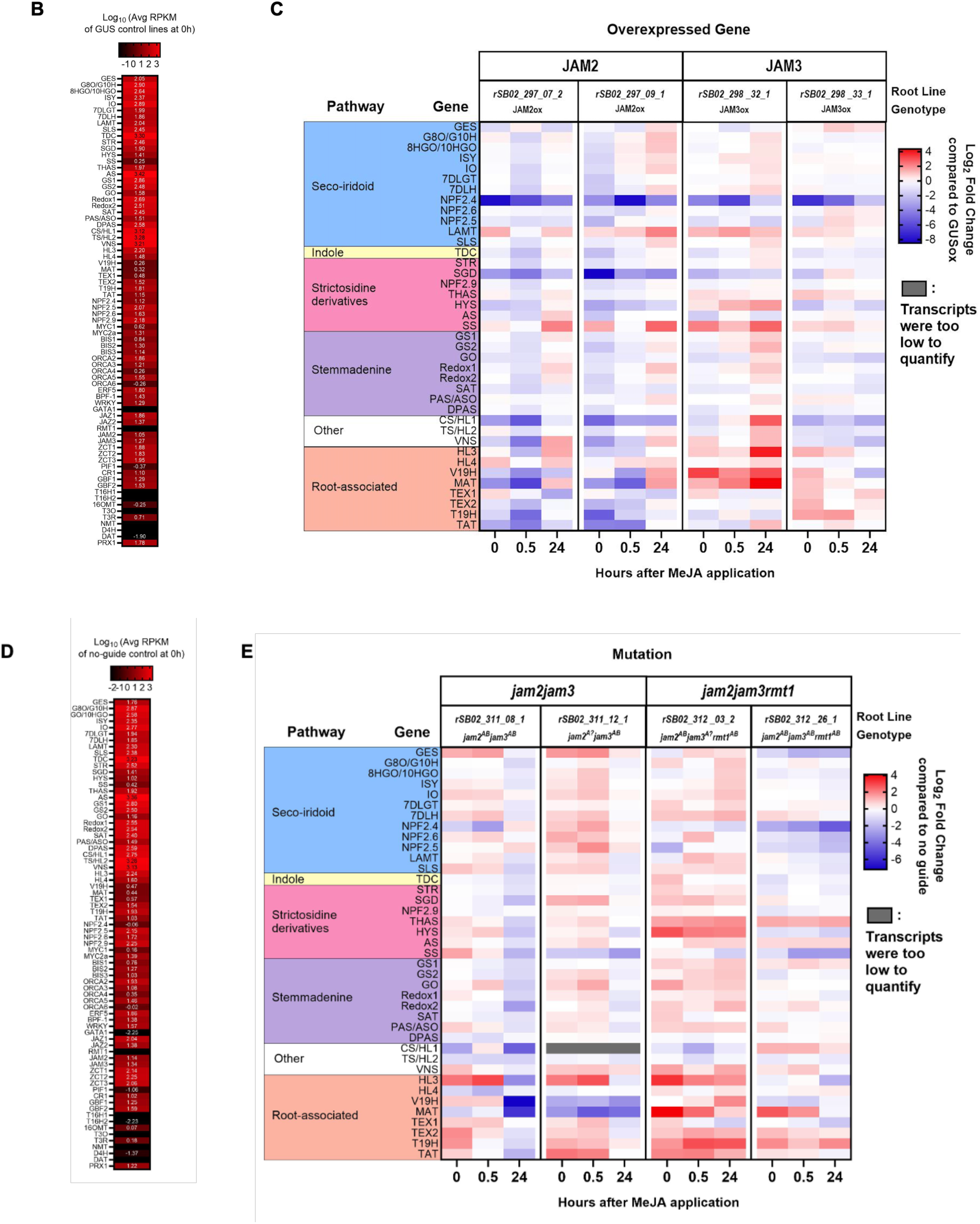
Gene expression profile of *CrJAM* overexpressing and CRISPR-mutated lines. **(A)** The *C. roseus* MIA biosynthetic pathway is color-coded to correspond to the same pathway genes in the heat map shown in part C and E; regulation of genes by MYC2a, BIS1/BIS2, and ORCA3 (Schweizer et al., 2018) is indicated with arrows (yellow, blue, and green, respectively). **(B)** This heat map shows the transcript abundance of MIA pathway in two *GUS* overexpressing hairy root lines as average reads per kilobase of transcript per million (RKPM) mapped reads at 0 hours after methyl jasmonate elicitation. **(C)** This heat map show changes in gene expression at 0, 0.5, and 24 hours after methyl jasmonate elicitation in transgenic hairy root lines overexpressing either *CrJAM2* or *CrJAM3*, normalized to *GUS* overexpression control lines. **(D)** This heat map shows the transcript abundance of MIA pathway in four hairy root lines overexpressing *Cas9* with no guides as average reads per kilobase of transcript per million (RKPM) mapped reads at 0 hours after methyl jasmonate elicitation. **(E)** This heat map show changes in gene expression at 0, 0.5, and 24 hours after methyl jasmonate elicitation in transgenic hairy root lines mutated using CRISPR-Cas9, i.e*. jam2jam3* or *jam2jam3rmt1*, normalized to a no guide control line. The functionality of the CrJAMs in these mutant lines are summarized in Table 1.

**Table 1.**
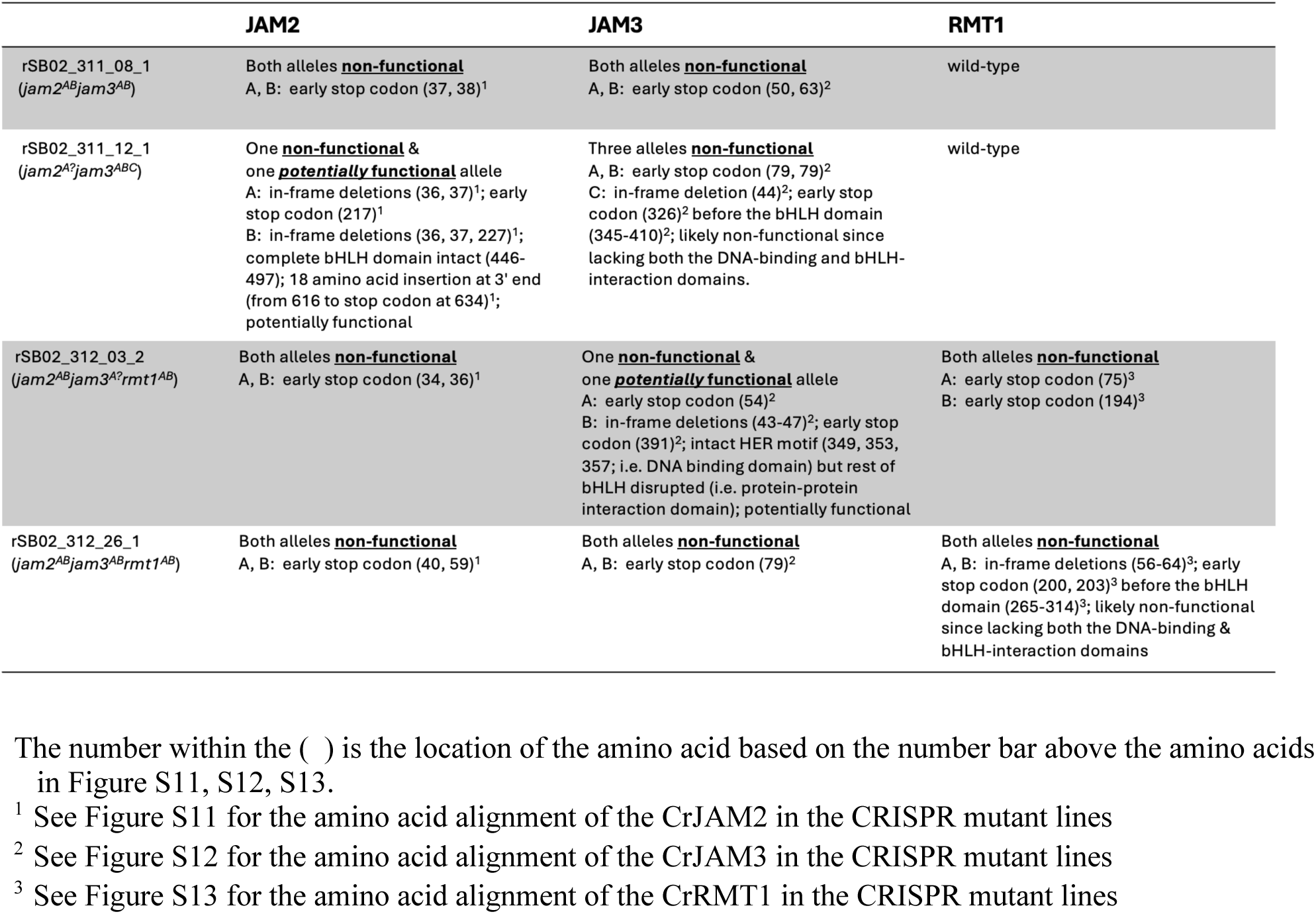
Summary of the functionality of the CrJAMs in the double and triple mutant lines.

In general, the constitutive overexpression of *CrJAM2* repressed the expression of most MIA pathway genes relative to the negative control lines at either 0 or 0.5 hr; specifically, upstream MIA transporter and pathway genes (*NPF2.4*, *TDC, SGD, HYS*), downstream MIA pathway genes (*CS/HL1, VNS*), and the root-associated pathway genes (*V19H, MAT, TEX2, T19H, TAT*) were strongly repressed by >75% in both *CrJAM2* overexpression lines relative to the negative control lines (Figure 5C). The constitutive overexpression of *CrJAM3* only strongly repressed the expression of a few MIA pathway genes at either 0 or 0.5 hr compared with the overexpression of *CrJAM2*; specifically, only *NPF2.4* and *SGD* were repressed by at least 75% in both *CrJAM3* lines. In contrast to the *CrJAM2* lines, the *CrJAM3* lines showed increased rather than decreased expression of genes encoding the root-associated pathway (*HL3, V19H, MAT*), by at least 2-fold relative to the negative control at either 0 or 0.5 hr, suggesting that *CrJAM2* may specifically repress the expression of the root-associated pathway. The transcript levels of most aerial-associated vindoline pathway genes were either too low to measure or very low relative to the other genes (not shown).

However, after 24 hrs, MJ resulted in higher MIA pathway gene expression in the *CrJAM* overexpressing lines (three of the four lines, except for rSB02_298_33_1) than the negative control lines; specifically, MIA pathway genes (*LAMT, SS, VNS*) increased by greater than 2-fold relative to the negative control lines; *CrJAM3* overexpressing line (rSB_02_298_32_1) showed the strongest increases in the expression of genes in serpentine and catharanthine biosynthesis (*SS*, *HL1/CS* by >4-fold) and the root-associated pathways (*HL3, V19H, MAT* by >8-fold) compared to negative controls. Only *NPF2.4* and *SGD* remained repressed (by >50%) 24 hrs after MJ addition in these three *CrJAM* lines.

The constitutive overexpression of *CrJAM2* (and *CrJAM3* to a lesser extent) in hairy root cultures repressed the expression of most MIA pathway genes at either 0 or 0.5 hr after MJ addition, consistent with the results from the promoter transactivation assay in seedlings; however, this repression in hairy root cultures was released 24 hrs after MJ addition, resulting in higher expression of MIA pathway genes compared to the MJ-treated negative control lines. In a subsequent section, we present how the level of activators to repressors correlates with the released repression of the MIA pathway.

### Deletion of *CrJAMs* in hairy root lines increased MIA pathway gene expression

We also generated hairy root lines with multiple *CrJAM* deletions and characterized their effect on MIA gene expression through RNA-Seq. We used an optimized CRISPR-Cas9 construct containing 13 introns and an array of eight guide RNAs to successfully introduce mutations to multiple *CrJAMs* (Grützner et al., 2021). Double and triple mutant lines were generated targeting the following combination of *CrJAMs*: 1) *CrJAM2* and *CrJAM3* (using pSB311, Figure S 6) or 2) *CrJAM2*, *CrJAM3*, and *CrRMT1* (using pSB312, Figure S 6); the negative control lines overexpressed *Cas9* with no guide RNAs.

CRISPR-Cas mediated genome editing is highly specific, nevertheless the occurrence of off-target mutations is possible (Doench et al., 2016). Ordon et al. (2017) did not observe any negative effects from off-targets in transgenic *Nicotiana benthamiana* and *A. thaliana* lines generated with the original pDGE vector system (used to construct pSB311 and pSB312). The presence of 2 or 3 mismatches between guide and target reduced the occurrence of indels in human cells (Slaymaker et al., 2016). Therefore, we minimized off-target mutations by choosing guides with greater than 4 or 5 mismatches (i.e. a maximum identity of 15/20 or 16/20 bp) to the closest off-targets (Table S 5).

Four double mutant lines and four triple mutant lines were characterized in detail for mutations in *CrJAM2*, *CrJAM3*, and *CrRMT1* (Figure S 9 – S 13). CRISPR-Cas9 consistently introduced mutations at each guide RNA location in the double and triple mutant lines (Figure S 9 and S 10) but did not introduce mutations in the target sites of the negative control line expressing *Cas9* with no guide RNAs (*CrJAMs* were sequenced in two control lines). These mutations ranged from small indels to large deletions (>1000 bp) between different sgRNA target sites, as well as the integration of reverse complementary sequences between target sites (Figure S 10). Surprisingly, many lines appeared to have homozygous mutations (7 homozygous out of 20 potentially different alleles). Only one chimeric line with three different mutations was observed (CrJAM3 of rSB02_311_12_01, Figure S 9), suggesting that the mutations might occur early on during hairy root formation.

While the mutations of all four double mutant and four triple mutant lines were sequenced and characterized (Figure S 11 – S 13), only two double mutant and two triple mutant lines were further investigated for gene expression profiling. The functionality of the CrJAMs in these lines is summarized in Table 1. The functionality of a specific CrJAM was predicted based on the presence of the complete bHLH domain. If the bHLH domain was truncated, this CrJAM was considered non-functional since both its DNA-binding and its bHLH-interacting domains were disrupted. If mutations occurred after the H-E-R motif, this CrJAM was considered potentially functional since DNA-binding might still be possible although its bHLH-interacting domain would be missing. Both CrJAM2 and CrJAM3 were non-functional for one double mutant (rSB02_311_08_1 or *jam2^AB^jam3^AB^*) while one allele of CrJAM2 was potentially functional for the other double mutant line (rSB02_311_12_1 or *jam2^A?^jam3^ABC^*). Similarly, all three CrJAMs were non-functional for one triple mutant (rSB02_312_26_1 or *jam2^AB^jam3^AB^rmt1^AB^*) while one allele of CrJAM3 was potentially functional for the other triple mutant line (rSB02_312_03_2 or *jam2^AB^jam3^A?^rmt1^AB^*).

These CRISPR-mutated hairy root lines were elicited with MJ (250 μM) and sampled for RNA-Seq at basal conditions (0 hr), 0.5 hr, and 24 hrs. The expression of most MIA pathway genes was either unaffected or slightly increased in both double mutant lines compared to the control lines at 0 and 0.5 hr (<u>∼</u>2-fold or less; Figure 5E); however, the expression of MIA pathway genes was not consistent between the two triple mutant lines. One line (rSB02_312_03_2 or *jam2^AB^jam3^A?^rmt1^AB^*) displayed increased expression of most MIA pathway genes while the other line (rSB02_312_26_1 or *jam2^AB^jam3^AB^rmt1^AB^*) showed increased expression of only a few genes. Key steps that were strongly repressed in the *CrJAM2* overexpressing lines (like *NPF2.4, TDC, SGD, HYS*, *VNS*) and in the root-associated pathway (*HL3, TEX2, T19H, TAT*) were no longer repressed in the *CrJAM* double mutant lines at 0 or 0.5 hr; the similar expression profile observed in both double mutant lines suggests that both alleles of *CrJAM2* in rSB02_311_12_1 were non-functional (behaved like *jam2^AB^jam3^AB^*). Only the expression of *CS/HL1* remained strongly repressed in both double mutant lines (so strongly repressed in rSB02_311_12_1 that the transcripts could not be quantified). The transcript levels of most aerial-associated vindoline pathway genes were either too low to measure or very low relative to the other genes (not shown).

However, after 24 hrs, MJ resulted in either no change or lower expression of most MIA pathway genes in the double mutant lines than the negative control lines; specifically, downstream MIA pathway genes (*SS, V19H, MAT*) were strongly decreased by greater than 75% relative to the negative control lines. The expression of MIA pathway genes at 24 hrs with the addition of MJ was not consistent between the two triple mutant lines: one triple mutant (rSB02_312_03_2) showed >2-fold increases in the expression of MIA pathway genes (10 out of 39 genes) while the other triple mutant (rSB02_312_26_1) showed >50% decreases in the expression of MIA pathway genes (8 out of 39 genes) relative to the control.

Overall, CRISPR-Cas9 editing was highly effective in *C. roseus* hairy root cultures with mutations at 100% of the guide RNA target sites. Double mutant lines showed slightly increased expression of the MIA pathway genes compared to the control lines while only one triple mutant line exhibited strong activation of MIA gene expression at 0 or 0.5 hr. However, at 24 hrs after MJ addition, the double mutant lines showed decreased expression of MIA pathway genes compared to the MJ-treated negative control lines. Next, we present how the level of activators to repressors correlates with this reversal in MIA pathway expression.

### Overexpression of *CrJAMs* repressed, while loss of *CrJAMs* activated, the basal expression of the transcriptional regulators of MIA biosynthesis

The overexpression or mutation of *CrJAMs* impacted the expression of transcriptional regulators similarly to that of the MIA pathway genes, as shown in Figure 6. As with the expression of MIA pathway genes, the constitutive overexpression of *CrJAM2*, and *CrJAM3* to a lesser extent, repressed the expression of both activators and repressors at either 0 or 0.5 hr (Figure 6); strongly repressed genes in the *CrJAM2* lines (by greater than 75% decrease compared to the *GUS* lines) included the activators *ORCA2*, *ORCA3*, *ORCA5*, and *ORCA6*, and the repressors *JAZ1* and *JAZ2*; *BIS2* and *BIS3* were also repressed (by less than 75%). Since MYC2a activates the expression of *BIS*, *ORCA*, and *JAZ* (see regulatory network in Figure 1), the constitutive overexpression of *JAMs* would be expected to outcompete MYC2a for promoter binding, resulting in repression of *BIS, ORCA, JAZ,* and MIA pathway genes as observed.

**Figure 6.**
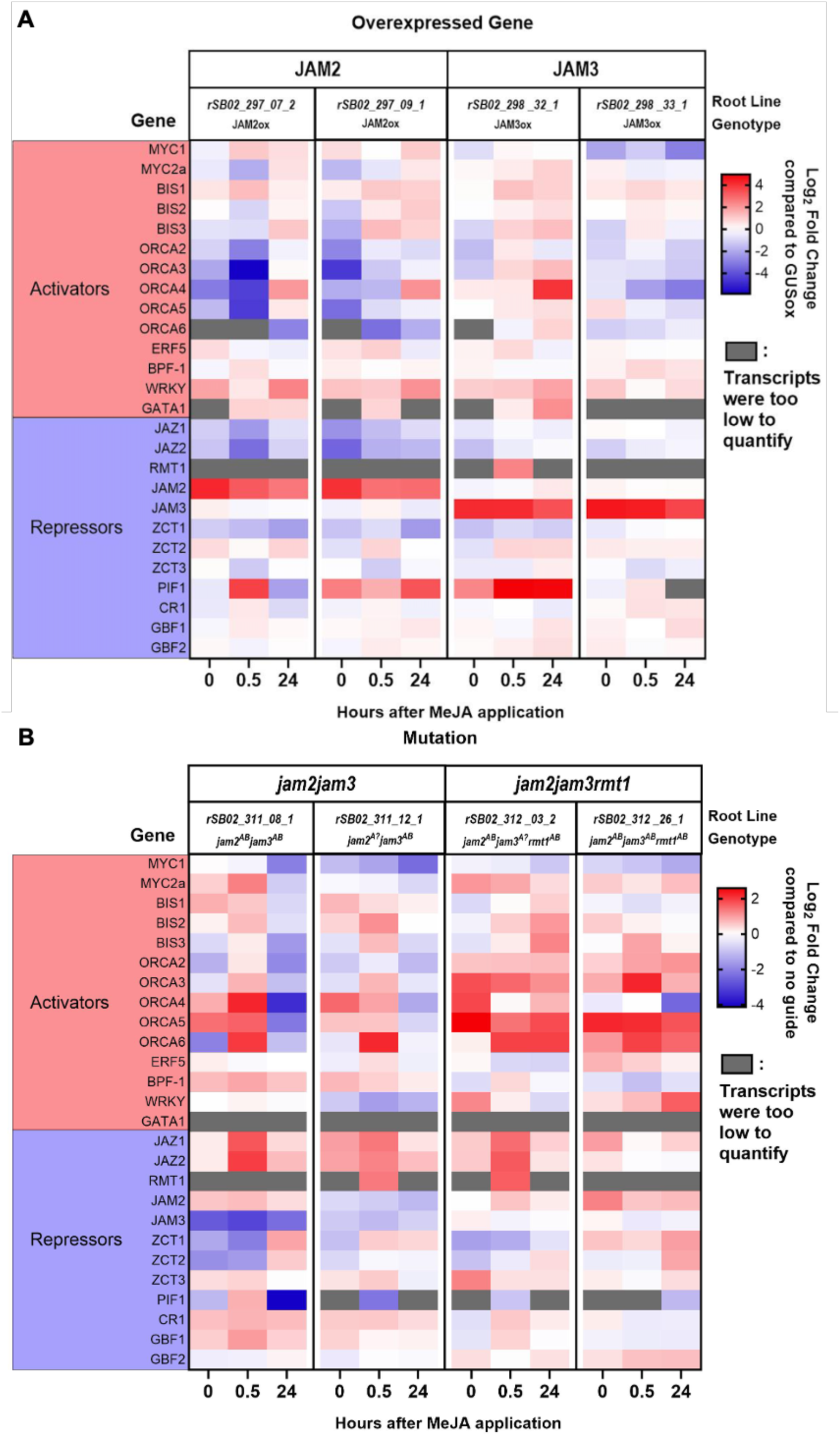
Gene expression profile of activators and repressors in CrJAM overexpressing and CRISPR-mutated lines. Heat maps show RNA-Seq data shifts in transcriptional activators and repressors at 0, 0.5, and 24 hours after methyl jasmonate elicitation in **(A)** transgenic hairy root lines overexpressing either *CrJAM2* or *CrJAM3*, normalized to *GUS* overexpression control lines or **(B)** transgenic hairy root lines where *CrJAMs* were mutated using CRISPR-Cas9, i.e. *jam2jam3* or *jam2jam3rmt1*, normalized to a no guide control line. See Figure 1 for the regulatory model and Figure 5 A for the biosynthetic pathway.

By 24 hr after MJ addition, the repression of most MIA activators in *CrJAM*-overexpressing lines was released; activators such as *MYC2a*, *BIS3*, and *ORCA4* were now induced relative to the negative control, while repressors such as *JAZ* and *ZCT1* remained repressed (Figure 6A). The increased expression of activators relative to repressors could explain the induction of MIA pathway genes at 24 hr in three of the four *CrJAM*-overexpressing lines.

Conversely, the double and triple mutations of *CrJAMs* resulted in higher expression of both activators (*BISs* and *ORCAs*) and repressors (*JAZs*) relative to the negative control at either 0 or 0.5 hr (Figure 6B) as expected. However, at 24 hr after MJ addition, the expression of the activators in the double mutant lines became repressed while the repressors remained induced relative to the negative control lines; the increased expression of repressors relative to activators could explain the inhibition of the MIA pathway genes at 24 hrs (Figure 5E). Unlike the double mutant, the activators and repressors in the triple mutant lines remained slightly induced relative to the negative control 24 hrs after MJ treatment.

## Discussion

In this study, we investigated the hypothesis that CrJAMs repress MIA biosynthesis. To answer this question, we overexpressed or silenced *CrJAM2* and *CrJAM3* using four complementary methods: transient silencing in leaves (VIGS), transient overexpression in seedlings (EASI), stable overexpression in hairy root cultures, and stable CRISPR-mediated knockouts in hairy root cultures. Each method explored the effect of CrJAMs in different contexts (tissue types, overexpression versus silencing, and application of exogenous JA) on the MIA pathway at different layers of regulation (promoter activation, transcript levels, and alkaloid levels). Despite major differences between the four methods, we consistently observed evidence of CrJAMs acting as repressors of the MIA biosynthetic pathway. We integrate this new evidence, the role of JAMs, into an updated model illustrating how the transcription factor triad of JAZ, MYC2a, and JAM regulates MIA biosynthesis in *C. roseus* (Figure 1).

Evidence for regulation by this transcriptional triad comes from both *A. thaliana* and *C. roseus* in the literature or in this study, suggesting a conserved pathway modulating the JA-mediated defense response. The triad consists of multiple feedforward and feedback loops that work together to create a time-constrained pulse of JA-activated gene expression. First, JAZ binds and holds MYC2 and JAMs captive in Arabidopsis (Chini et al., 2007; Sasaki-Sekimoto et al., 2014) and *C. roseus* (Schweizer et al., 2018; only CrRMT1 was studied in Patra et al. 2018). In the presence of JA-Ile, JAZ is degraded and MYC2 and JAMs are released. In *C. roseus,* MYC2a then activates a feedforward cascade (Figure 1), increasing the expression of *ORCA* and *BIS* activators and the expression of MIA pathway enzymes (Zhang et al., 2011; Schweizer et al., 2018). ORCA and BIS further activate expression of MIA pathway genes (Van der Fits and Memelink, 2000; Van Moerkercke et al., 2015; Van Moerkercke et al., 2016; Schweizer et al., 2018).

JAM proteins are simultaneously released from interaction with JAZ repressors (Sasaki-Sekimoto et al., 2013; Song et al., 2013) and compete with MYC2 in binding to the same promoter elements in *A. thaliana* (Fonseca et al., 2014) and *C. roseus*. For instance, in *C. roseus*, Zhang et al. (2011) and Patra et al. (2018) showed that MYC2a and CrRMT1 activate and repress the *ORCA3* promoter, respectively, by binding to the same T/G-boxes. In addition to regulation of *ORCA3*, we consistently observed that CrJAM2 and CrJAM3 repressed the expression of the *BIS* and *ORCA* family (Figure 1, line 2 and line 4), as follows: VIGS of *CrJAM2* and *CrJAM3* together led to >2-fold increases in *BIS1* and *BIS2* (Figure 3B); overexpression of *CrJAM2* or *CrJAM3* in seedlings significantly reduced the promoter activity of *BIS1*, *BIS2*, and *ORCA3* (Figure 4); overexpression of *CrJAM2* or *CrJAM3* in hairy roots decreased the basal transcript levels of *ORCA2*, *ORCA3*, *ORCA5*, and *ORCA6* by over 75%, and of *BIS2, BIS3*, and *ORCA4* more moderately (Figure 6A). Finally, both the *jam* double and triple mutant lines exhibited increased expression of multiple *BIS* and *ORCA* genes at either 0 or 0.5 hrs after MeJA addition. These results are consistent with Schweizer et al. (2018), who observed repression of *pBIS1* with transient overexpression of *CrJAM2* or *CrJAM3*, and Patra et al. (2018), who showed repression of *pORCA3* with transient overexpression of *CrRMT1*. Overall, CrJAM2, CrJAM3, and CrRMT1 have overlapping roles in repressing the expression of multiple *BIS* and *ORCA* activators.

Like its repression of *BIS* and *ORCA* activators, JAMs repress the expression of the MEP-secoiridoid and MIA pathway. We observed that the overexpression of either *CrJAM2* or *CrJAM3* broadly repressed genes in these pathways. For example, transient expression of either *CrJAM2* or *CrJAM3* significantly reduced promoter activity of MEP-secoiridoid (*pG8O*) and MIA pathway (*pGO*), genes previously shown to be regulated by the derepressed CrMYC2a^D126N^ in *C. roseus* flowers (Schweizer et al., 2018). However, we did not observe an effect on *pZCT1*, *pLAMT*, *pSTR1*, and *pT19H* genes that were previously shown to be activated by CrMYC2a^D126N^ (Schweizer et al., 2018; Traverse et al., 2024). In addition, we observed that CrJAMs repressed *pT16H1* but activated *pT16H2*, which were not shown to be regulated by CrMYC2a^D126N^ (Schweizer et al. 2018). These differences between our results, Patra et al. (2018), and Schweizer et al. (2018) might be attributed to the distinct role of the individual CrJAMs and to the evaluation methods and tissue context used.

Furthermore, RNAseq profiling of hairy root lines showed the broad extent of regulation by CrJAMs. To our knowledge, this is the first evidence of the broad regulation of the MIA pathway by CrJAMs. For instance, we discovered that CrJAM2 repress genes in the upstream MIA pathway (Figure 1, line 3: *NPF2.4*, *TDC, SGD, HYS*), downstream MIA pathway genes (line 3: *CS/HL1, VNS*), and the root-associated pathway (*V19H, MAT, TEX2, T19H, TAT*) by greater than 75% compared to the control lines (Figure 5) and the expression of the MEP-secoiridoid pathway genes by less than 75% (Figure 1, line 5).

Conversely, the *jam* double mutant lines (and one triple mutant line: rSB02_312_3_02 or *jam2^AB^jam3^A?^rmt^AB^*) exhibited an overall increase in the expression of the MIA regulatory genes and nearly all MIA pathway genes at either 0 or 0.5 hr (Figure 5, Figure 6); importantly, the increase in the basal gene expression relative to no guide control lines (at 0 hr) occurred in the absence of MJ addition. In these lines, *NPF2.4, TDC, SGD, HYS, VNS,* and most root-associated pathway genes (*HL3, TEX2, T19H,* and *TAT*) were either no longer or less strongly repressed as compared to the *CrJAM2* lines. CrJAM2 appears to specifically repress the root-associated pathway (*HL3, V19H, MAT, TEX2, T19H, TAT*) since the expression of most of these genes increased in the *CrJAM3* overexpression and both the double and triple mutant lines. Only the expression of *CS/HL1* remained strongly repressed in both double mutant lines, suggesting that the wildtype CrRMT1 is repressing *CS/HL1*. Our results indicate that the individual CrJAMs repress overlapping as well as different genes in the MIA pathway.

Consistent with the effect on promoter activity and transcript level, we observed 33 – 86% increases in MIA alkaloids in *CrJAM2* and *CrJAM3* silenced leaves (Figure 3C). Patra et al. (2018) showed that the overexpression of *CrRMT1* in *C. roseus* hairy root cultures decreased ajmalicine, catharanthine, and tabersonine production. Unfortunately, our hairy root lines with either the overexpression or deletion of *CrJAMs* showed poor viability with prolonged subcultures and subsequently died before alkaloid profiling could be completed. The constitutive overexpression of the *CrJAM* repressors or the *Cas9* endonuclease may have caused toxicity during prolonged culturing (Zhang et al., 2019; Gutierrez-Valdes et al., 2020). Future studies could employ an inducible expression system to improve root culture viability.

In parallel with the feedforward induction of activators, MYC2 initiates a negative feedback loop by activating both *JAZ* and *JAM* expression, previously shown in both Arabidopsis and *C. roseus* (Chini et al., 2007; Sasaki-Sekimoto et al., 2013, Patra et al. 2018). In addition, we observed that CrJAMs regulate *CrJAZ* expression in our stable hairy root cultures. When *CrJAM2* was constitutively overexpressed in *C. roseus* hairy roots (and *CrJAM3* a lesser extent), *CrJAZ1* and *CrJAZ2* expression was repressed compared to the control, at all timepoints after MeJA addition. Similarly, in the CRISPR-mutated lines, *CrJAZ1* and *CrJAZ2* expression was elevated compared to the control, at all timepoints after MeJA addition. This finding is consistent with the prior observation that CrMYC2a regulates *CrJAZ* expression (Patra et al., 2018), and since CrJAMs directly compete with CrMYC2a targets, CrJAMs repress *CrJAZ* expression. However, CrJAMs do not likely regulate *CrMYC2a* expression since the *CrMYC2a* promoter lacks G-box and its variants; in Arabidopsis, JAM and MYC2 proteins also do not interact with each other (Nakata et al., 2013). This is consistent with the findings of Schweizer et al. (2018) who found that endogenous expression of *MYC2a* did not change with overexpression of *CrMYC2a^D126N^* in *C. roseus* flowers. This negative regulation of *JAZ* expression by CrJAMs may be responsible for the reversal of gene expression trends we observed 24 hours after JA addition. Repression of *CrJAZ1* and *CrJAZ2* by CrJAM2 and CrJAM3 adds further nuance to the JA-mediated regulatory triad involved in MIA biosynthesis regulation (Figure 1, line 1).

## Conclusion

In summary, we investigated CrJAMs as repressors of MIA biosynthesis. We characterized the recently identified CrJAM2 and CrJAM3 through complementary methods involving distinct tissue types, overexpression versus silencing, and application of the MIA pathway elicitor, JA. Based on this collective evidence, we identified CrJAMs as repressors of the MIA regulatory and MIA biosynthetic pathway. CrMYC2a and CrJAMs appear to modulate the extent of JA-mediated expression of MIA biosynthesis. *CrJAM2* appears to specifically repress the root-associated pathway (*HL3, V19H, MAT, TEX2, T19H, TAT*) while CrRMT1 appears to repress *CS/HL1* expression. Our results suggest that CrJAM2 and CrJAM3, along with the previously characterized CrRMT1, share redundant function, contributing to the robust regulation of JA-inducible MIA biosynthesis in *C. roseus*.

## Author contributions: CRediT

Samuel Breselge: Writing – original draft, review & editing, Investigation, Conceptualization. Krystyna K. F. Traverse: Writing – original draft, review & editing, Investigation. Lauren F. Cole-Osborn: Writing – original draft, review & editing, Investigation. Suphinya Sathitloetsakun: Investigation; Kristina Erwardt: Investigation; Jie Wang: Investigation; Kevin L. Childs: Writing – review & editing, Supervision, Investigation. Erin J. Cram: Writing – review & editing, Supervision, Funding acquisition. Carolyn W.T. Lee-Parsons: Writing – original draft, review & editing, Supervision, Project administration, Funding acquisition, Conceptualization.

## Acknowledgements

The authors would like to thank Dr. Johannes Stuttmann and Dr. Sylvestre Marillonnet for generously providing the pDGE vector.

## Glossary

VIGS: Viral Induced Gene Silencing
EASI: Efficient Agro-mediated Seedling Infiltration
JAZ: Jasmonate ZIM-domain protein
JA: Jasmonic Acid
MJ: Methyl Jasmonate
bHLH: Basic Helix-Loop-helix

## References

1. Aerts, R. J., Gisi, D., Carolis, E. De, Luca, V. De, and Baumann, T. W. (1994). Methyl jasmonate vapor increases the develpmentally controlled synthesis of alkaloids in Catharanthus and Cinchona seedlings. Plant Cell 5, 635–643. doi: 10.1111/j.1365-313X.1994.00635.x

2. Aslam, J., Khan, S. H., Siddiqui, Z. H., Fatima, Z., Maqsood, M., Bhat, M. A., et al. (2010). Catharanthus roseus (L.) G. Don. An Important Drug: It’s Applications and Production. Int. J. Compr. Pharm. 1, 1–16.

3. Bailey, P. C., Martin, C., Toledo-Ortiz, G., Quail, P. H., Huq, E., Heim, M. A., et al. (2003). Update on the Basic Helix-Loop-Helix Transcription Factor Gene Family in Arabidopsis thaliana. Plant Cell Online 15, 2497–2502. doi: 10.1105/tpc.151140

4. Bernonville, T. D., Carqueijeiro, I., Lanoue, A., Lafontaine, F., Sánchez Bel, P., Liesecke, F., et al. (2017). Folivory elicits a strong defense reaction in Catharanthus roseus: Metabolomic and transcriptomic analyses reveal distinct local and systemic responses. Sci. Rep. 7, 1–14. doi: 10.1038/srep40453

5. Bolger, A. M., Lohse, M., and Usadel, B. (2014). Trimmomatic: A flexible trimmer for Illumina sequence data. Bioinformatics 30, 2114–2120. doi: 10.1093/bioinformatics/btu170

6. Chini, A., Fonseca, S., Fernández, G., Adie, B., Chico, J. M., Lorenzo, O., et al. (2007). The JAZ family of repressors is the missing link in jasmonate signalling. Nature 448, 666–671. doi: 10.1038/nature06006

7. Chini, A., Gimenez-Ibanez, S., Goossens, A., and Solano, R. (2016). Redundancy and specificity in jasmonate signalling. Curr. Opin. Plant Biol. 33, 147–156. doi: 10.1016/j.pbi.2016.07.005

8. Cole-Osborn, L. F., McCallan, S. A., Prifti, O., Abu, R., Sjoelund, V., and Lee-Parsons, C. W. T. (2024a). The role of the Golden2-like (GLK) transcription factor in regulating terpenoid indole alkaloid biosynthesis in Catharanthus roseus. Plant Cell Rep. 43, 1–20. doi: 10.1007/s00299-024-03208-9

9. Cole-Osborn, L. F., Meehan, E., and Lee-Parsons, C. W. T. (2024b). Critical parameters for robust Agrobacterium-mediated transient transformation and quantitative promoter assays in Catharanthus roseus seedlings. Plant Direct 8, 1–14. doi: 10.1002/pld3.596

10. Colinas, M., Pollier, J., Vaneechoutte, D., Malat, D. G., Schweizer, F., De Milde, L., et al. (2021). Subfunctionalization of Paralog Transcription Factors Contributes to Regulation of Alkaloid Pathway Branch Choice in Catharanthus roseus. Front. Plant Sci. 12. doi: 10.3389/fpls.2021.687406

11. Dobin, A., Davis, C. A., Schlesinger, F., Drenkow, J., Zaleski, C., Jha, S., et al. (2013). STAR: Ultrafast universal RNA-seq aligner. Bioinformatics 29, 15–21. doi: 10.1093/bioinformatics/bts635

12. Doench, J. G., Fusi, N., Sullender, M., Hegde, M., Vaimberg, E. W., Donovan, K. F., et al. (2016). Optimized sgRNA design to maximize activity and minimize off-target effects of CRISPR-Cas9. Nat. Biotechnol. 34, 184–191. doi: 10.1038/nbt.3437

13. Dombrecht, B., Gang, P. X., Sprague, S. J., Kirkegaard, J. A., Ross, J. J., Reid, J. B., et al. (2007). MYC2 differentially modulates diverse jasmonate-dependent functions in Arabidopsis. Plant Cell 19, 2225–2245. doi: 10.1105/tpc.106.048017

14. Engler, C., Youles, M., Gruetzner, R., Ehnert, T.-M., Werner, S., Jones, J. D. G., et al. (2014). A Golden Gate Modular Cloning Toolbox for Plants. ACS Synth. Biol. 3, 839–843. doi: 10.1021/sb4001504

15. Fernandez-Pozo, N., Rosli, H. G., Martin, G. B., and Mueller, L. A. (2015). The SGN VIGS Tool: User-Friendly Software to Design Virus-Induced Gene Silencing (VIGS) Constructs for Functional Genomics. Mol. Plant 8, 486–488. doi: 10.1016/j.molp.2014.11.024

16. Fonseca, S., Fernández-Calvo, P., Fernández, G. M., Díez-Díaz, M., Gimenez-Ibanez, S., López-Vidriero, I., et al. (2014). bHLH003, bHLH013 and bHLH017 are new targets of JAZ repressors negatively regulating JA responses. PLoS One 9, 1–12. doi: 10.1371/journal.pone.0086182

17. Franke, J., Kim, J., Hamilton, J. P., Zhao, D., Pham, G. M., Wiegert-Rininger, K., et al. (2018). Data from: Gene discovery in Gelsemium highlights conserved gene clusters in monoterpene indole alkaloid biosynthesis. Available at: https://datadryad.org/stash/dataset/doi:10.5061/dryad.08vv50n

18. Franke, J., Kim, J., Hamilton, J. P., Zhao, D., Pham, G. M., Wiegert-Rininger, K., et al. (2019). Gene Discovery in Gelsemium Highlights Conserved Gene Clusters in Monoterpene Indole Alkaloid Biosynthesis. Chembiochem 20, 83–87. doi: 10.1002/cbic.201800592

19. Goklany, S., Rizvi, N. F., Loring, R. H., Cram, E. J., and Lee-Parsons, C. W. T. (2013). Jasmonate-dependent alkaloid biosynthesis in Catharanthus Roseus hairy root cultures is correlated with the relative expression of Orca and Zct transcription factors. Biotechnol. Prog. 29, 1367–1376. doi: 10.1002/btpr.1801

20. Góngora-Castillo, E., Childs, K. L., Fedewa, G., Hamilton, J. P., Liscombe, D. K., Magallanes-Lundback, M., et al. (2012). Development of Transcriptomic Resources for Interrogating the Biosynthesis of Monoterpene Indole Alkaloids in Medicinal Plant Species. PLoS One 7. doi: 10.1371/journal.pone.0052506

21. Grützner, R., Martin, P., Horn, C., Mortensen, S., Cram, E. J., Lee-Parsons, C. W. T., et al. (2021). High-efficiency genome editing in plants mediated by a Cas9 gene containing multiple introns. Plant Commun. 2, 1–15. doi: 10.1016/j.xplc.2020.100135

22. Gutierrez-Valdes, N., Häkkinen, S. T., Lemasson, C., Guillet, M., Oksman-Caldentey, K. M., Ritala, A., et al. (2020). Hairy Root Cultures—A Versatile Tool With Multiple Applications. Front. Plant Sci. 11. doi: 10.3389/fpls.2020.00033

23. Haas, J. H., Moore, L. W., Ream, W., and Manulis, S. (1995). Universal PCR primers for detection of phytopathogenic Agrobacterium strains. Appl. Environ. Microbiol. 61, 2879–2884. doi: 10.1128/aem.61.8.2879-2884.1995

24. Hao, Y., Zong, X., Ren, P., Qian, Y., and Fu, A. (2021). Basic helix-loop-helix (Bhlh) transcription factors regulate a wide range of functions in arabidopsis. Int. J. Mol. Sci. 22. doi: 10.3390/ijms22137152

25. Heim, M. A., Jakoby, M., Werber, M., Martin, C., Weisshaar, B., and Bailey, P. C. (2003). The basic helix-loop-helix transcription factor family in plants: A genome-wide study of protein structure and functional diversity. Mol. Biol. Evol. 20, 735–747. doi: 10.1093/molbev/msg088

26. Huang, H., Gao, H., Liu, B., Fan, M., Wang, J., Wang, C., et al. (2018). bHLH13 Regulates Jasmonate-Mediated Defense Responses and Growth. Evol. Bioinforma. 14, 1–8. doi: 10.1177/1176934318790265

27. Li, C., Wood, J. C., Hamilton, J. P., Rodriguez Lopez, C. E., Payne, R. M. E., Serna Guerrero, D. A., et al. (2023). Data for: Single-cell multi-omics in the medicinal plant Catharanthus roseus [Dataset]. doi: doi.org/10.5061/dryad.d2547d851

28. Li, C. Y., Leopold, A. L., Sander, G. W., Shanks, J. V., Zhao, L., and Gibson, S. I. (2013). The ORCA2 transcription factor plays a key role in regulation of the terpenoid indole alkaloid pathway. BMC Plant Biol. 13, 1–17. doi: 10.1186/1471-2229-13-155

29. Li, Z., Huang, Y., Shen, Z., Wu, M., Huang, M., Hong, S. B., et al. (2024). Advances in functional studies of plant MYC transcription factors. Springer Berlin Heidelberg. doi: 10.1007/s00122-024-04697-8

30. Liao, Y., Smyth, G. K., and Shi, W. (2014). FeatureCounts: An efficient general purpose program for assigning sequence reads to genomic features. Bioinformatics 30, 923–930. doi: 10.1093/bioinformatics/btt656

31. Liscombe, D. K., and O’Connor, S. E. (2011). A virus-induced gene silencing approach to understanding alkaloid metabolism in Catharanthus roseus. Phytochemistry 72, 1969–1977. doi: 10.1016/j.phytochem.2011.07.001

32. Livak, K. J., and Schmittgen, T. D. (2001). Analysis of relative gene expression data using real-time quantitative PCR and the 2-ΔΔCT method. Methods 25, 402–408. doi: 10.1006/meth.2001.1262

33. Lorenz, O., Chico, J. M., Sánchez-Serran, J. J., and Solano, R. (2004). JASMONATE-INSENSITIVE1 Encodes a MYC Transcription Factor Essential to Discriminate between Different Jasmonate-Regulated Defense Responses in Arabidopsis. Plant Cell 16, 1938–1950. doi: 10.1105/tpc.022319.with

34. Mitchell, A. L., Attwood, T. K., Babbitt, P. C., Blum, M., Bork, P., Bridge, A., et al. (2019). InterPro in 2019: Improving coverage, classification and access to protein sequence annotations. Nucleic Acids Res. 47, D351–D360. doi: 10.1093/nar/gky1100

35. Mortensen, S., Bernal-Franco, D., Cole, L. F., Sathitloetsakun, S., Cram, E. J., and Lee-Parsons, C. W. T. (2019). Easi transformation: An efficient transient expression method for analyzing gene function in Catharanthus roseus seedlings. Front. Plant Sci. 10, 1–17. doi: 10.3389/fpls.2019.00755

36. Mortensen, S., Cole, L. F., Bernal-Franco, D., Sathitloetsakun, S., Cram, E. J., and Lee-Parsons, C. W. T. (2022). “EASI Transformation Protocol: An Agrobacterium-Mediated Transient Transformation Protocol for Catharanthus roseus Seedlings,” in Methods in Molecular Biology, eds. V. Courdevault and S. Besseau (Springer Nature), 249–262. Available at: https://link.springer.com/protocol/10.1007/978-1-0716-2349-7_18 (Accessed May 11, 2023).

37. Nakata, M., Mitsuda, N., Herde, M., Koo, A. J. K., Moreno, J. E., Suzuki, K., et al. (2013). A bHLH-type transcription factor, ABA-INDUCIBLE BHLH-TYPE TRANSCRIPTION FACTOR/JA-ASSOCIATED MYC2-LIKE1, acts as a repressor to negatively regulate jasmonate signaling in Arabidopsis. Plant Cell 25, 1641–1656. doi: 10.1105/tpc.113.111112

38. Noble, R. L., Beer, C. T., and Cutts, J. H. (1958). Role of chance observations in chemotherapy: Vinca rosea. Ann. N. Y. Acad. Sci. 76, 882–894. doi: 10.1111/j.1749-6632.1958.tb54906.x

39. Ordon, J., Gantner, J., Kemna, J., Schwalgun, L., Reschke, M., Streubel, J., et al. (2017). Generation of chromosomal deletions in dicotyledonous plants employing a user-friendly genome editing toolkit. Plant J. 89, 155–168. doi: 10.1111/tpj.13319

40. Patra, B., Pattanaik, S., Schluttenhofer, C., and Yuan, L. (2018). A network of jasmonate-responsive bHLH factors modulate monoterpenoid indole alkaloid biosynthesis in Catharanthus roseus. New Phytol. 217, 1566–1581. doi: 10.1111/nph.14910

41. Paul, P., Singh, S. K., Patra, B., Sui, X., Pattanaik, S., and Yuan, L. (2017). A differentially regulated AP2/ERF transcription factor gene cluster acts downstream of a MAP kinase cascade to modulate terpenoid indole alkaloid biosynthesis in Catharanthus roseus. New Phytol. 213, 1107–1123. doi: 10.1111/nph.14252

42. Pauw, B., Hilliou, F. A. O., Martin, V. S., Chatel, G., De Wolf, C. J. F., Champion, A., et al. (2004). Zinc finger proteins act as transcriptional repressors of alkaloid biosynthesis genes in Catharanthus roseus. J. Biol. Chem. 279, 52940–52948. doi: 10.1074/jbc.M404391200

43. Pazour, G. J., Ta, C. N., and Das, A. (1992). Constitutive mutations of Agrobacterium tumefaciens transcriptional activator virG. J. Bacteriol. 174, 4169–4174. doi: 10.1128/jb.174.12.4169-4174.1992

44. Peebles, C. A. M., Hughes, E. H., Shanks, J. V., and San, K. Y. (2009). Transcriptional response of the terpenoid indole alkaloid pathway to the overexpression of ORCA3 along with jasmonic acid elicitation of Catharanthus roseus hairy roots over time. Metab. Eng. 11, 76–86. doi: 10.1016/j.ymben.2008.09.002

45. Pollier, J., Vanden Bossche, R., Rischer, H., and Goossens, A. (2014). Selection and validation of reference genes for transcript normalization in gene expression studies in Catharanthus roseus. Plant Physiol. Biochem. 83, 20–25. doi: 10.1016/j.plaphy.2014.07.004

46. Qi, T., Wang, J., Huang, H., Liu, B., Gao, H., Liu, Y., et al. (2015). Regulation of Jasmonate-Induced Leaf Senescence by Antagonism between bHLH Subgroup IIIe and IIId Factors in Arabidopsis. Plant Cell 27, 1634–1649. doi: 10.1105/tpc.15.00110

47. Qian, Y., Zhang, T., Yu, Y., Gou, L., Yang, J., Xu, J., et al. (2021). Regulatory Mechanisms of bHLH Transcription Factors in Plant Adaptive Responses to Various Abiotic Stresses. Front. Plant Sci. 12. doi: 10.3389/fpls.2021.677611

48. Rizvi, N. F., Cornejo, M., Stein, K., Weaver, J., Cram, E. J., and Lee-Parsons, C. W. T. (2015). An efficient transformation method for estrogen-inducible transgene expression in Catharanthus roseus hairy roots. Plant Cell. Tissue Organ Cult. 120, 475–487. doi: 10.1007/s11240-014-0614-1

49. Rizvi, N. F., Weaver, J. D., Cram, E. J., and Lee-Parsons, C. W. T. (2016). Silencing the Transcriptional Repressor, ZCT1, Illustrates the Tight Regulation of Terpenoid Indole Alkaloid Biosynthesis in Catharanthus roseus Hairy Roots. PLoS One 11, e0159712. doi: 10.1371/journal.pone.0159712

50. Sasaki-sekimoto, Y., Jikumaru, Y., Obayashi, T., Saito, H., Masuda, S., Kamiya, Y., et al. (2013). Basic Helix-Loop-Helix Transcription Factors JAM2 , and JAM3 Are Negative Regulators of Jasmonate. 163, 291–304. doi: 10.1104/pp.113.220129

51. Sasaki-Sekimoto, Y., Jikumaru, Y., Obayashi, T., Saito, H., Masuda, S., Kamiya, Y., et al. (2013). Basic helix-loop-helix transcription factors JASMONATE-ASSOCIATED MYC2-LIKE1 (JAM1), JAM2, and JAM3 are negative regulators of jasmonate responses in Arabidopsis. Plant Physiol. 163, 291–304. doi: 10.1104/pp.113.220129

52. Schweizer, F., Colinas, M., Pollier, J., Van Moerkercke, A., Vanden Bossche, R., de Clercq, R., et al. (2018). An engineered combinatorial module of transcription factors boosts production of monoterpenoid indole alkaloids in Catharanthus roseus. Metab. Eng. 48, 150–162. doi: 10.1016/j.ymben.2018.05.016

53. Sibéril, Y., Benhamron, S., Memelink, J., Giglioli-Guivarc’h, N., Thiersault, M., Boisson, B., et al. (2001). Catharanthus roseus G-box binding factors 1 and 2 act as repressors of strictosidine synthase gene expression in cell cultures. Plant Mol. Biol. 45, 477–488. doi: 10.1023/A:1010650906695

54. Sigrist, C. J. A., De Castro, E., Cerutti, L., Cuche, B. A., Hulo, N., Bridge, A., et al. (2013). New and continuing developments at PROSITE. Nucleic Acids Res. 41, 344–347. doi: 10.1093/nar/gks1067

55. Škubník, J., Pavlíčková, V. S., Ruml, T., and Rimpelová, S. (2021). Vincristine in combination therapy of cancer: Emerging trends in clinics. Biology (Basel). 10. doi: 10.3390/biology10090849

56. Slaymaker, I. M., Gao, L., Zetsche, B., Scott, D. A., Yan, W. X., and Zhang, F. (2016). Rationally engineered Cas9 nucleases with improved specificity. Science (80-.). 351, 84–88. doi: 10.1126/science.aad5227

57. Song, S., Qi, T., Fan, M., Zhang, X., Gao, H., Huang, H., et al. (2013). The bHLH Subgroup IIId Factors Negatively Regulate Jasmonate-Mediated Plant Defense and Development. PLoS Genet. 9, 1–19. doi: 10.1371/journal.pgen.1003653

58. Suttipanta, N., Pattanaik, S., Kulshrestha, M., Patra, B., Singh, S. K., and Yuan, L. (2011). The transcription factor CrWRKY1 positively regulates the terpenoid indole alkaloid biosynthesis in Catharanthus roseus. Plant Physiol. 157, 2081–2093. doi: 10.1104/pp.111.181834

59. Svec, D., Tichopad, A., Novosadova, V., Pfaffl, M. W., and Kubista, M. (2015). How good is a PCR efficiency estimate: Recommendations for precise and robust qPCR efficiency assessments. Biomol. Detect. Quantif. 3, 9–16. doi: 10.1016/j.bdq.2015.01.005

60. Thaler, J. S., Humphrey, P. T., and Whiteman, N. K. (2012). Evolution of jasmonate and salicylate signal crosstalk. Trends Plant Sci. 17, 260–270. doi: 10.1016/j.tplants.2012.02.010

61. Thines, B., Katsir, L., Melotto, M., Niu, Y., Mandaokar, A., Liu, G., et al. (2007). JAZ repressor proteins are targets of the SCFCOI1 complex during jasmonate signalling. Nature 448, 661–665. doi: 10.1038/nature05960

62. Toledo-Ortiz, G., Huq, E., and Quail, P. H. (2003). The Arabidopsis basic/helix-loop-helix transcription factor family. Plant Cell 15, 1749–1770. doi: 10.1105/tpc.013839

63. Traverse, K. K. F., Breselge, S., Trautman, J. G., Dee, A., Wang, J., Childs, K. L., et al. (2024). Characterization of the ZCTs, a subgroup of Cys2-His2 zinc finger transcription factors regulating alkaloid biosynthesis in Catharanthus roseus. Plant Cell Rep. 43, 1–27. doi: 10.1007/s00299-024-03295-8

64. Traverse, K. K. F., Mortensen, S., Trautman, J. G., Danison, H., Rizvi, N. F., and Lee-Parsons, C. W. T. (2022). “Generation of Stable Catharanthus roseus Hairy Root Lines with Agrobacterium rhizogenes,” in Methods in Molecular Biology, ed. A. G. Fett-Neto (Springer), 129–144. doi: 10.1007/978-1-0716-2185-1_11

65. Van Der Fits, L., Memelink, J., Fits, L., and Memelink, J. (2000). ORCA3, a jasmonate-responsive transcriptional regulator of plant primary and secondary metabolism. Science (80-.). 289, 295–297. doi: 10.1126/science.289.5477.295

66. Van der Heijden, R., Jacobs, D., Snoeijer, W., Hallard, D., and Verpoorte, R. (2004). The Catharanthus Alkaloids: Pharamacognosy and Biotechnology. Curr Med Chem 11, 607–628.

67. Van Moerkercke, A., Steensma, P., Gariboldi, I., Espoz, J., Purnama, P. C., Schweizer, F., et al. (2016). The basic helix-loop-helix transcription factor BIS2 is essential for monoterpenoid indole alkaloid production in the medicinal plant Catharanthus roseus. Plant J. 88, 3–12. doi: 10.1111/tpj.13230

68. Van Moerkercke, A., Steensma, P., Schweizer, F., Pollier, J., Gariboldi, I., Payne, R., et al. (2015). The bHLH transcription factor BIS1 controls the iridoid branch of the monoterpenoid indole alkaloid pathway in Catharanthus roseus. Proc. Natl. Acad. Sci. U. S. A. 112, 8130–8135. doi: 10.1073/pnas.1504951112

69. Warnes, G. R., Bolker, B., Bonebakker, L., Gentleman, R., Liaw, W. H. A., Lumley, T., et al. (2009). gplots: Various R Programming Tools for Plotting Data. R Packag. version.

70. Wasternack, C., and Strnad, M. (2018). Jasmonates: News on Occurrence, Biosynthesis, Metabolism and Action of an Ancient Group of Signaling Compounds. Int. J. Mol. Sci. 19, 2539. doi: 10.3390/ijms19092539

71. Weber, E., Engler, C., Gruetzner, R., Werner, S., and Marillonnet, S. (2011). A Modular Cloning System for Standardized Assembly of Multigene Constructs. PLoS One 6, e16765. doi: 10.1371/journal.pone.0016765

72. Zhang, H., Hedhili, S., Montiel, G., Zhang, Y., Chatel, G., Pré, M., et al. (2011). The basic helix-loop-helix transcription factor CrMYC2 controls the jasmonate-responsive expression of the ORCA genes that regulate alkaloid biosynthesis in Catharanthus roseus. Plant J. 67, 61–71. doi: 10.1111/j.1365-313X.2011.04575.x

73. Zhang, Y., Malzahn, A. A., Sretenovic, S., and Qi, Y. (2019). The emerging and uncultivated potential of CRISPR technology in plant science. Nat. Plants 5, 778–794. doi: 10.1038/s41477-019-0461-5

