## Supplements for "CrJAMs are transcriptional repressors of the monoterpenoid indole alkaloid pathway in *Catharanthus roseus*"

SB, KKFT, LFCO are shared first authors
[1] Northeastern University, Department of Biology, Boston, MA, USA 02115
[2] Northeastern University, Department of Chemical Engineering, Boston, MA
[3] Northeastern University, Department of Bioengineering, Boston, MA
[4] Northeastern University, Department of Chemistry and Chemical Biology, Boston, MA
[5] Leibniz University, Department of Life Science, Hanover, Germany
[6] Michigan State University, Department of Plant Biology, East Lansing, MI, USA

### Supplementary Figures

**A.**

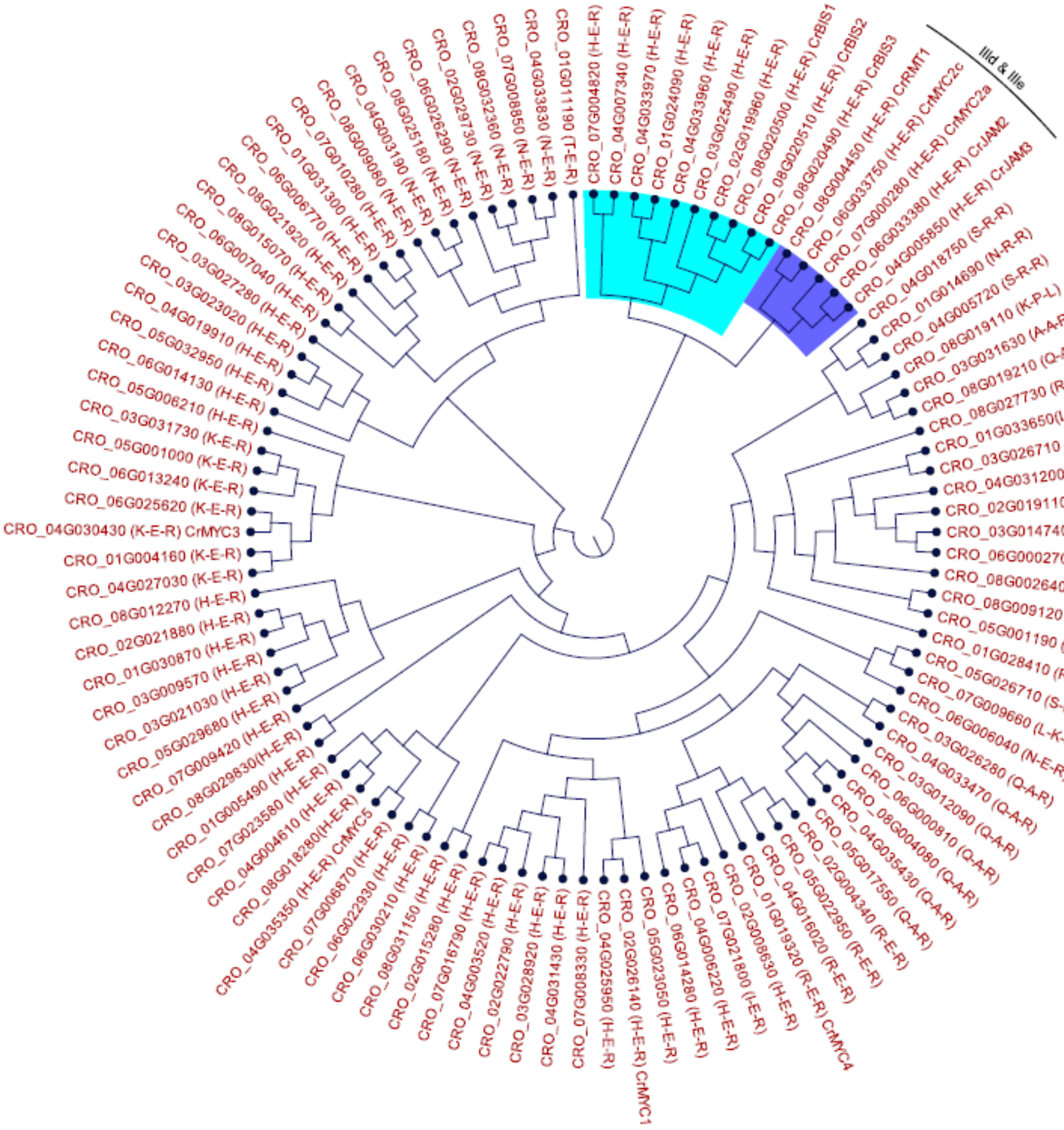

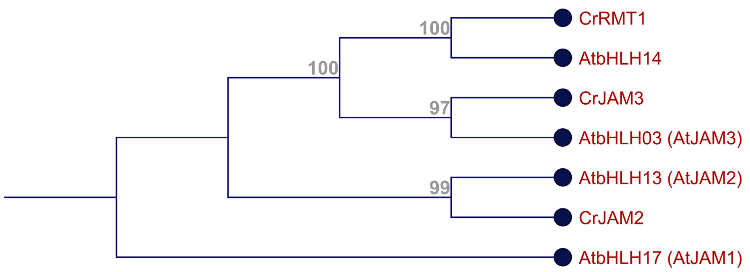


**B.**

**Figure S 1. (A)** Phylogenetic tree of bHLH domains from *C. roseus* is displayed as a circular cladogram. bHLH domains of the bHLH IIId (JAMs) and IIIe (MYCs) subgroup are indicated in dark blue. Domains of the related bHLH IVa (containing BIS1, BIS2, and BIS3) subgroup are indicated in light blue. Given in parenthesis are the amino acid identities of the 5-9-13 motif within the bHLH domain. CrJAMs, CrMYCs, and CrBIS all contain the H-E-R motif within the bHLH domain. **(B)** Phylogenetic tree of *Arabidopsis thaliana* bHLH IIId transcription factors and their homologs in *C. roseus*. *C. roseus* proteins were identified in a BlastP search with *A. thaliana* bHLH IIId TFs as query (E-value < E-30). CrMYC2a was excluded from the tree as the homolog AtMYC2 is part of the bHLH IIIe subgroup. The tree is displayed as cladogram with bootstrapping values as an indication of the confidence in the arrangement of the branches in the phylogenetic tree.

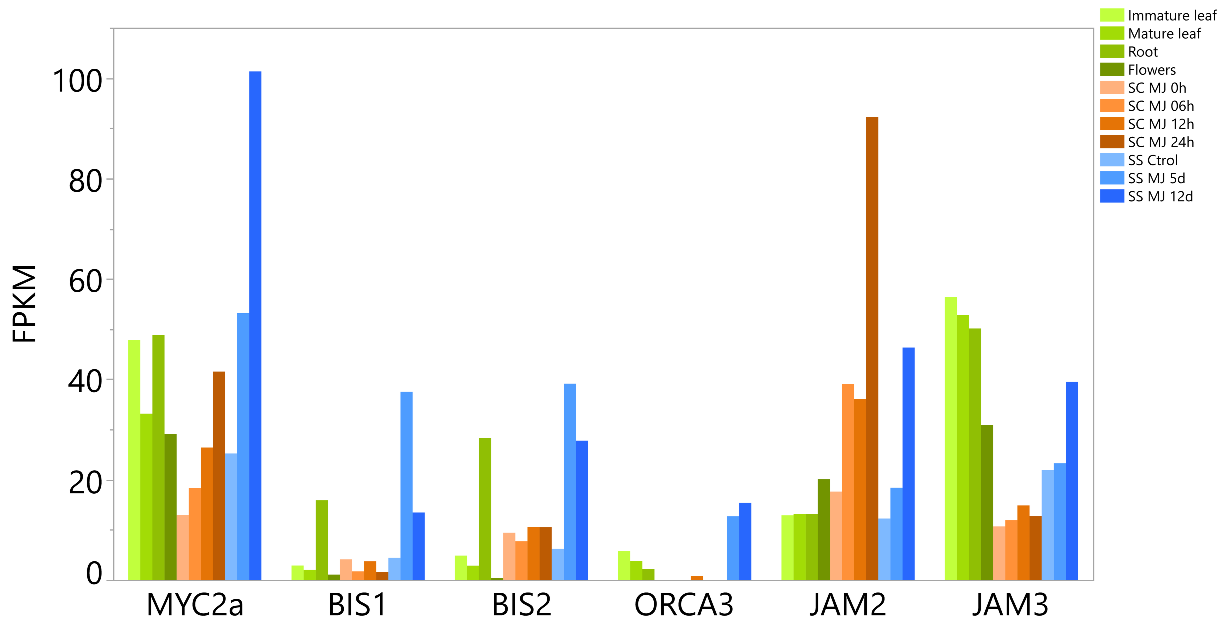


**Figure S 2.** Expression of *C. roseus* transcription factors in different tissues elicited with methyl jasmonate

Transcript levels of *MYC2a*, *BIS1*, *BIS2*, *ORCA3*, *JAM2*, and *JAM3* in terms of FPKM are plotted for different tissues, with or without methyl jasmonate treatment, using the *C. roseus* transcriptome data from Góngora-Castillo et al., 2012. FPKM: fragments per kilobase of exon per million fragments mapped. Different tissues (greens); SC: suspension culture (oranges); SS: sterile seedlings (blues); MJ: methyl jasmonate. *MYC2a*: cra_locus_8576_iso_1_len_2817_ver_3; *BIS1*: cra_locus_12822_iso_1_len_1418_ver_3; *BIS2*: cra_locus_2845_iso_5_len_1555_ver_3; *ORCA3*: cra_locus_20632_iso_2_len_765_ver_3; *JAM2*: cra_locus_4594_iso_2_len_2637_ver_3; *JAM3*: cra_locus_9367_iso_5_len_2326_ver_3.

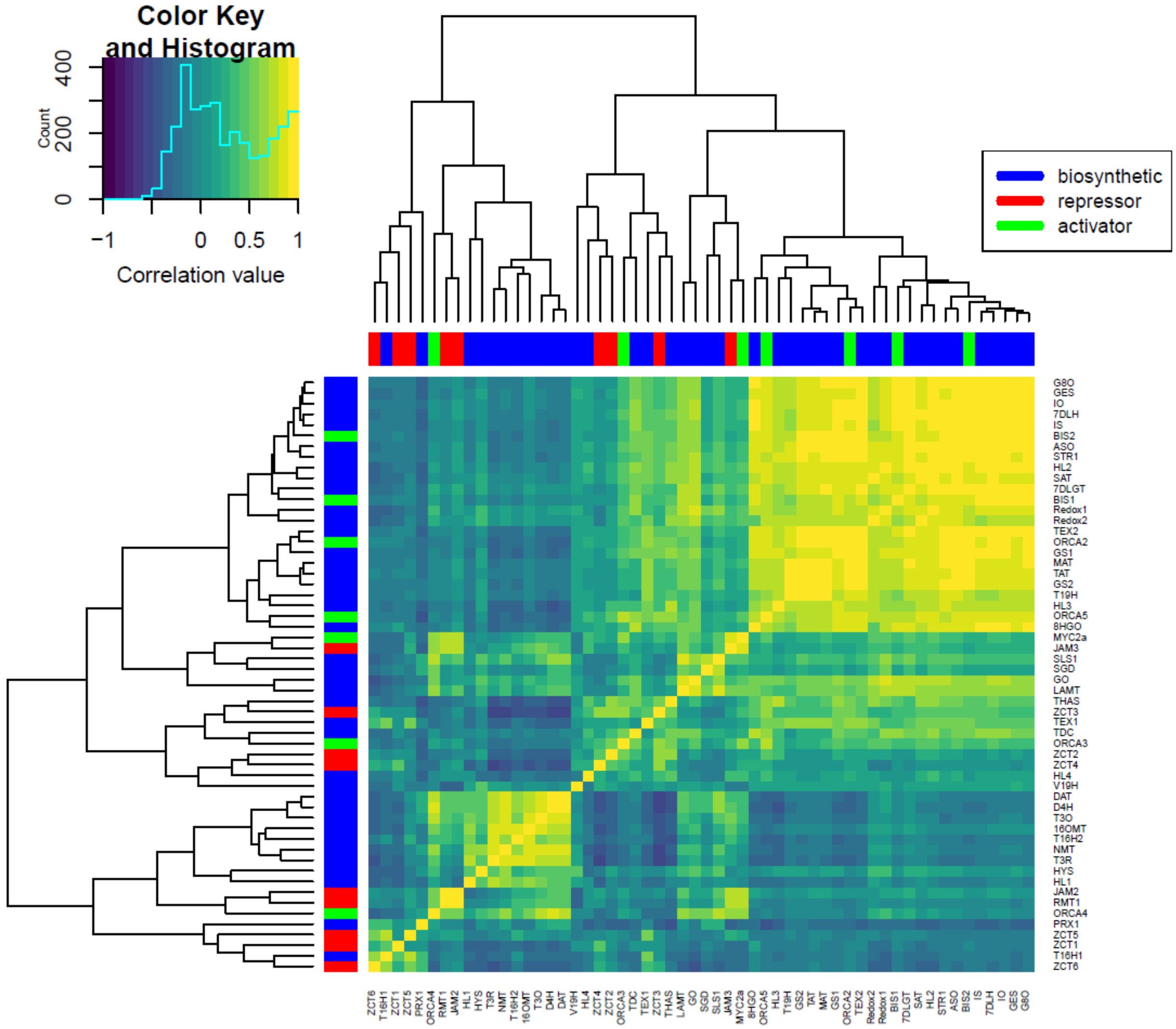


**Figure S 3**. Correlation heat-map and hierarchical clustering of *CrJAM2* and *CrJAM3* with MIA pathway enzymes and transcription factors

The correlation heat-map and hierarchical clustering was drawn using transcriptomic data from Góngora-Castillo et al. (2012). In the heat map, the left and top color bar with tree indicate the expression of MIA pathway enzymes (shown in blue), MIA transcriptional activators (shown in green), and MIA transcriptional repressors (shown in red) that are strongly correlated. The color key and histogram at the top left of the heat map indicate the level of positive correlation (shown as green to yellow) or negative or inverse correlation (shown as green to dark blue). For example,

*JAM3* and *MYC2a* cluster on one subtree and *JAM2* and *RMT1* cluster on one subtree, suggesting a strong coregulation and positive correlation (upregulated together).

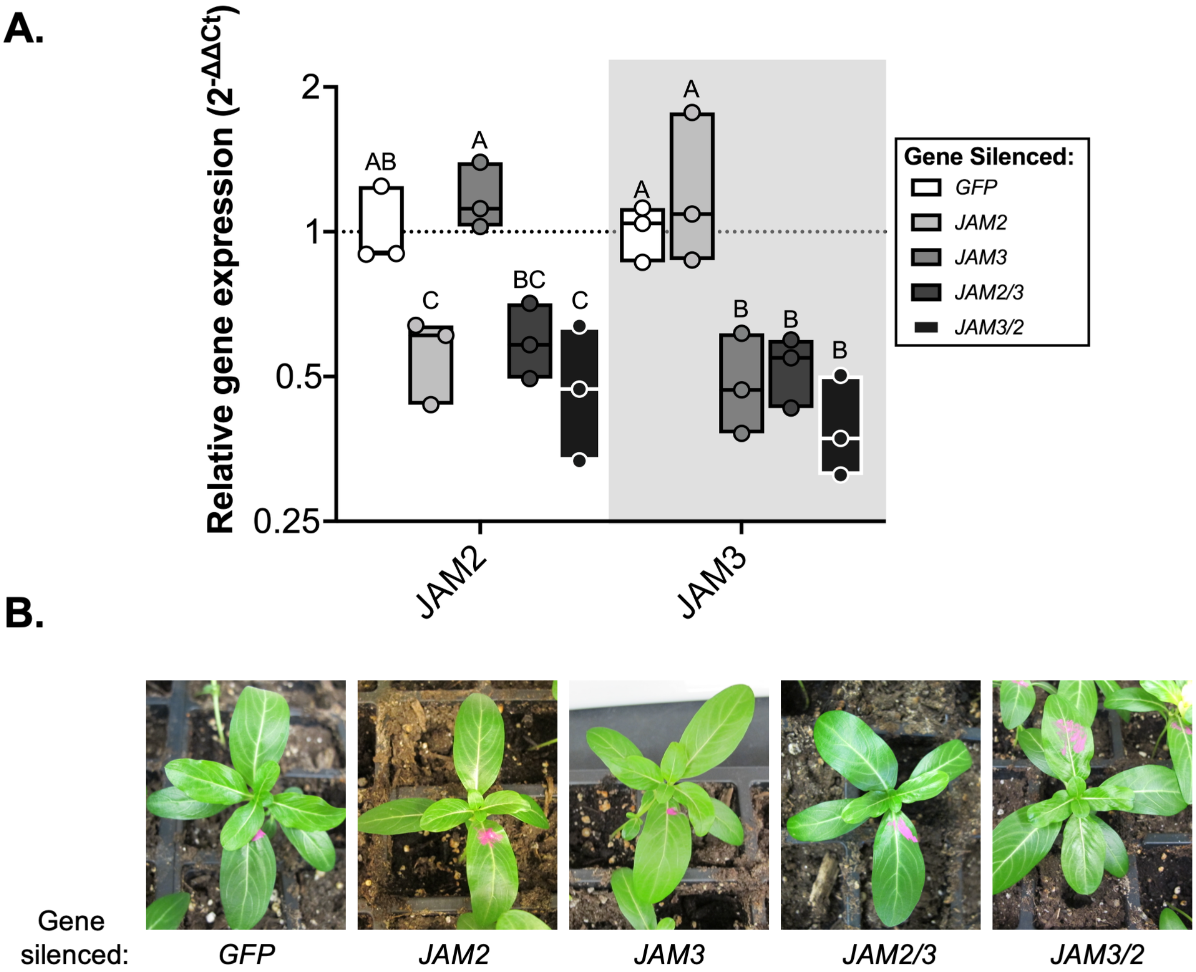


**Figure S 4.** Efficiency of *JAM*-silencing with VIGS constructs in different combinations **(A)** *JAM2* and *JAM3* were silenced either individually or together (*JAM2/3*, *JAM3/2*). The order of the genes in the pTRV2 plasmid was swapped (*JAM2/3* vs. *JAM3/2*). Silencing efficiency was consistently around 50% whether genes were silenced separately, together, or in different orders. Each sample is from an individual plant, grown under low white light (about 15 µmol $m^{-2}$ $s^{-1}$) (n = 3). The fully emerged leaf at the time of infection was marked with pink paint. One leaf from both the first and second leaf pair to emerge after infection was pooled for RNA extraction. Relative gene expression was measured with qPCR and calculated using the $2^{-\Delta\Delta Ct}$ method (Livak and Schmittgen, 2001) relative to the control condition (*GFP*-silenced plants) and normalized relative to the housekeeping gene, *SAND* (Pollier et al., 2014). Different letters indicate values that are significantly different (Tukey-Kramer HSD test on ΔΔCt values, $p<0.05$). The horizontal line within the boxes shows the median, and the ends of the boxes show the lowest and highest data point. **(B)** *JAM*-silenced plants showed either mild or no visible phenotype compared to *GFP*-silenced plants.

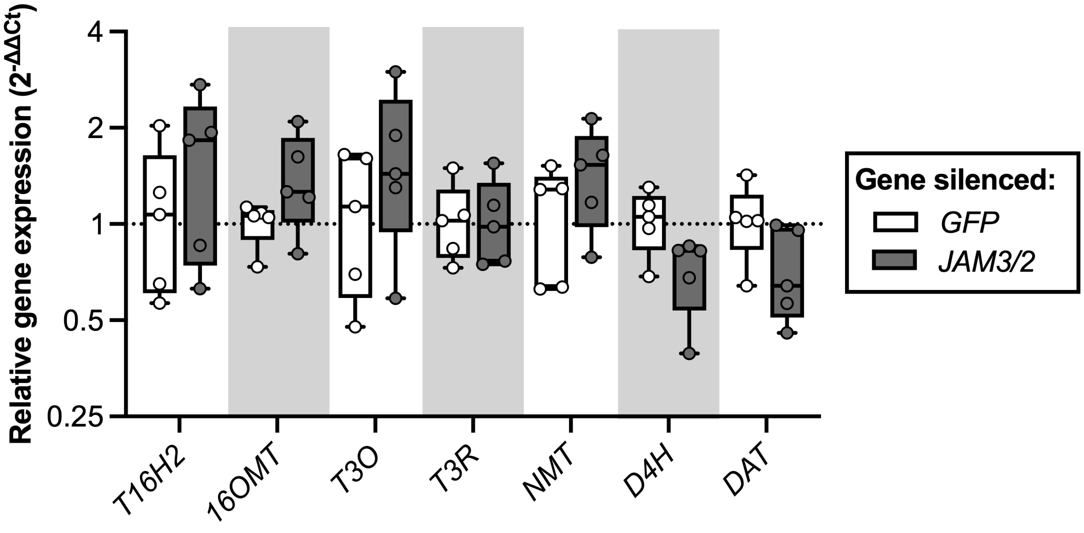


**Figure S 5.** Vindoline pathway gene expression in *JAM*-silenced plants

Expression of the vindoline pathway genes was not significantly affected by *JAM3/2* silencing in immature leaves (n = 5). Relative gene expression was measured with qPCR and calculated using the $2^{-\Delta\Delta Ct}$ method (Livak and Schmittgen, 2001) relative to the control condition (*GFP*-silenced plants) and normalized relative to the housekeeping gene, *SAND* (Pollier et al., 2014). Boxes represent the 25th and 75th percentile with a line marking the median. Whiskers extend to the minimum and maximum.

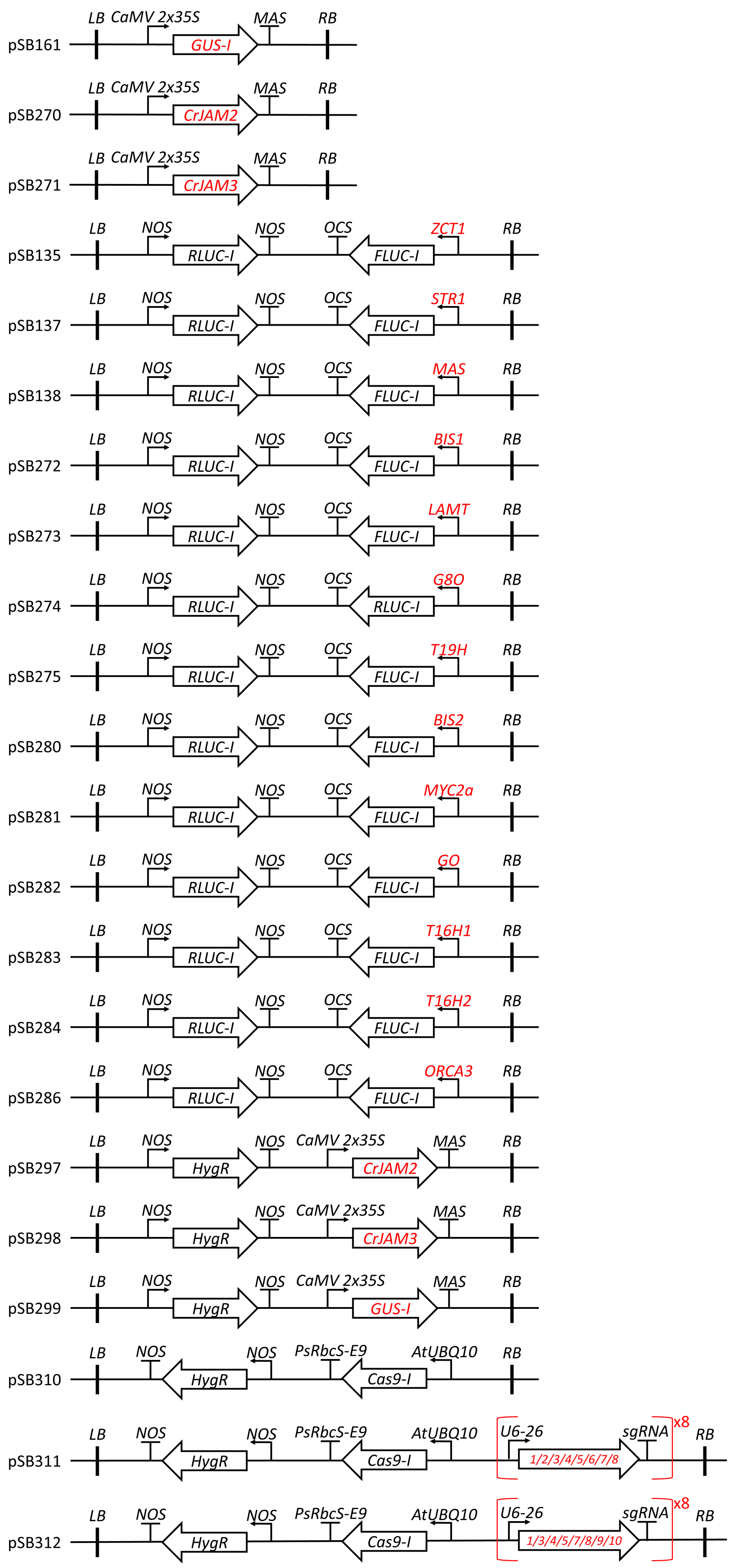
pSB161
LB
CaMV 2x35S
GUS-I
MAS
RB
pSB270
CrJAM2
pSB271
CrJAM3
pSB135
NOS
RLUC-I
OCS
FLUC-I
ZCT1
pSB137
STR1
pSB138
MAS
pSB272
BIS1
pSB273
LAMT
pSB274
G8O
pSB275
T19H
pSB280
BIS2
pSB281
MYC2a
pSB282
GO
pSB283
T16H1
pSB284
T16H2
pSB286
ORCA3
pSB297
HygR
pSB298
pSB299
pSB310
PsRbcS-E9
AtUBQ10
Cas9-I
pSB311
U6-26
sgRNA
x8
1/2/3/4/5/6/7/8
pSB312
1/3/4/5/7/8/9/10

**Figure S 6.** Vector cartoons

Final vectors (except pSB310-312) were assembled in the pSB90 vector backbone (Mortensen et al., 2019) containing *VirGN54D* for increased *Agrobacterium* virulence and *KanR* as the bacterial selection marker. pSB310-312 were assembled in the improved pDGE vector system (Grützner et al., 2021). The *pU6-26::sgRNA_#:scaffold* is repeated eight times in pSB311 and pSB312, with each repeat containing one of the guides. The genes *GUS*, *FLUC*, *RLUC,* and *Cas9* contain at least one intron and are therefore marked with *-I*. Highlighted in red are elements that are different between similar vectors.

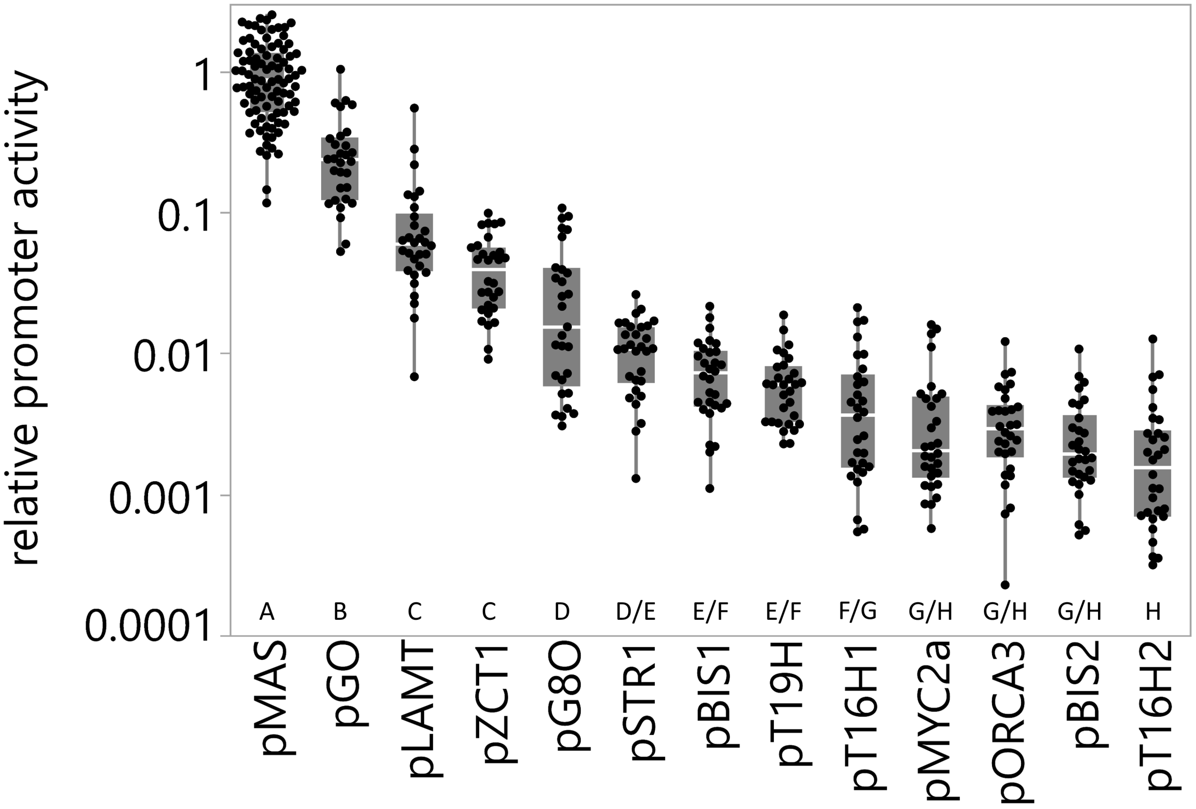


**Figure S 7.** Relative activity of different *C. roseus* promoters in transiently transformed seedlings

*C. roseus* seedlings were vacuum-infiltrated with a combination of two strains of *A. tumefaciens* (total $OD_{600} = 0.4$): 1) strain containing the promoter-driven *FLUC-I* reporter (Figure S 6: $OD_{600} = 0.06$), and 2) strain containing the *CaMV2x35S* driven *GUS* as control for transactivation (pSB161, Addgene ID #123197). Samples were taken three days post-infection. The relative promoter activity was calculated by dividing the FLUC (firefly luciferase) activity of each sample by the average FLUC activity of the *pMAS::FLUC* control (pSB138, set to 1). The experiment was carried out in three independent assays with ten replicates per condition for each independent assay (n = 30 per condition, except *pMAS*). The white horizontal line within the boxes shows the median, the ends of the boxes show the 1st and 3rd quantile, and whiskers show the lowest and highest data point. Different letters indicate values that are significantly different (Tukey-Kramer HSD test on log-transformed data, $p<0.05$).

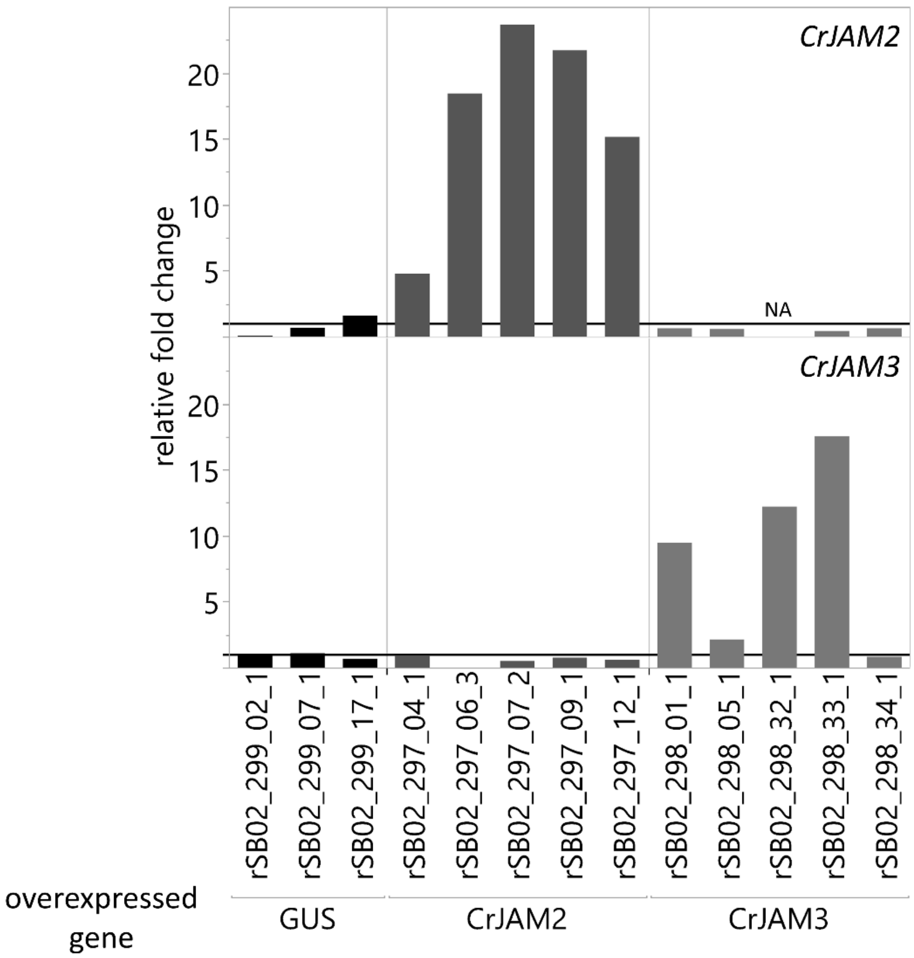


**Figure S 8.** Relative expression of *CrJAM2* and *CrJAM3* in transgenic *GUS*- and *CrJAM*-overexpressing hairy root lines

Relative expression of *CrJAM2* (top) and *CrJAM3* (bottom) in hairy root lines was determined by qPCR. Hairy root lines overexpressed either *GUS* (control line, pSB299), *CrJAM2* (pSB297), or *CrJAM3* (pSB298). Relative gene expression was calculated using the $2^{-\Delta\Delta Ct}$ method (Livak and Schmittgen, 2001) relative to the control condition (*GUS* control lines) and normalized relative to the housekeeping gene, *SAND* (Pollier et al., 2014). Average expression of the *GUS*-lines was set to 1. Displayed are means of two technical replicates. NA: not analyzed.

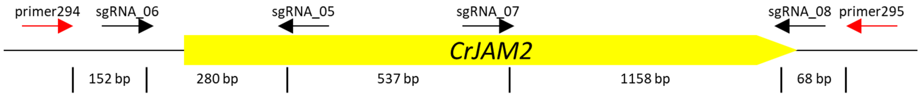


| Line | Allele | sgRNA_06 | sgRNA_05 | sgRNA_07 | sgRNA_08 |
|---|---|---|---|---|---|
| **wt gDNA/rSB02_310 lines** | A (homozygous) | | | | |
| **rSB02_311_07_1** | A (homozygous) | **286 bp deletion** | | **-1** | **-4** |
| **rSB02_311_08_1** | A | -3 | **-8** | **-8** | **+1** |
| | B | -2 | **-33+1** | **-1** | **-4** |
| **rSB02_311_12_1** | A | -2 | -6 | **-1** | **-1** |
| | B | -2 | -6 | -3 | **-1** |
| **rSB02_311_21_2** | A (homozygous) | -3 | **-5** | **+1** | **-1** |

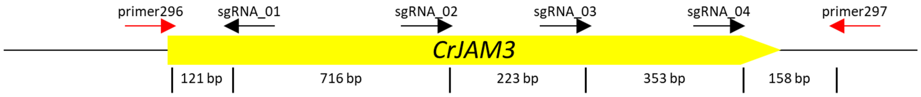


| Line | Allele | sgRNA_01 | sgRNA_02 | sgRNA_03 | sgRNA_04 |
|---|---|---|---|---|---|
| **wt gDNA/rSB02_310 lines** | A (homozygous) | | | | |
| **rSB02_311_07_1** | A (homozygous) | **941 bp deletion** | | | **-1** |
| **rSB02_311_08_1** | A | **1290 bp deletion** | | | |
| | B | **1291 bp deletion** | | | |
| **rSB02_311_12_1** | A | **-4** | **-7** | **-5** | **-5** |
| | B | **-4** | **-4** | **-1** | **-1** |
| | C | -3 | **-7** | **-5** | **-1** |
| **rSB02_311_21_2** | A (homozygous) | **717 bp deletion** | | **+1** | **-4** |

**Figure S 9.** Characterization of CRISPR-Cas9 deletions in *CrJAM2* and *CrJAM3*

Overview of mutations detected in *C. roseus* hairy root lines with guides targeting *CrJAM2* and *CrJAM3*. Primers shown in red were used for cloning and sequencing of the genes. Additional internal primers were used for sequencing as needed. Major deletions and frame-shift mutations are bolded: rSB02_311 denotes lines transformed with pSB311 (Figure S 6), containing eight guides targeting *CrJAM2* and *CrJAM3*; rSB02_310 denotes lines transformed with pSB310 (Figure S 6), containing the *Cas9* and *HygR* (*hpt*) sequence but no guides; wt: denotes gDNA from wild type seedlings. Lines rSB02_311_08_1 and rSB02_311_12_1 were pursued for RNA sequencing (Figure 5, Figure 6). This figure is modified from our previous publication (Grützner et al., 2021).

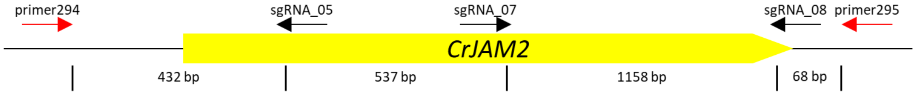


| Line | Allele | sgRNA_05 | sgRNA_07 | sgRNA_08 |
|---|---|---|---|---|
| wt gDNA/rSB02_310 lines | A (homozygous) | | | |
| rSB02_312_03_2 | A | **550 bp deletion** | | **-4-4-1-2** |
| | B | **1711 bp deletion** | | |
| rSB02_312_18_1 | A (homozygous) | -3 | **-2** | **-1** |
| rSB02_312_23_5 | A | **-5** | **+1** | **+1** |
| | B | **558 bp deletion** | | **-16 & T to G** |
| rSB02_312_26_1 | A | **+1** | **-14** | **-19** |
| | B | **reverse complementary & additional ~350 bp deletion** | | **-34** |

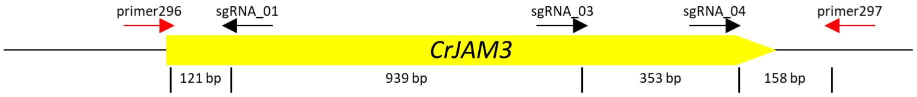


| Line | Allele | sgRNA_01 | sgRNA_03 | sgRNA_04 |
|---|---|---|---|---|
| wt gDNA/rSB02_310 lines | A (homozygous) | | | |
| rSB02_312_03_2 | A | **reverse complementary** | | -3 |
| | B | -15 | **-5** | -63 |
| rSB02_312_18_1 | A (homozygous) | **reverse complementary (sgRNA_01-03) & 344 bp deletion (sgRNA_03-04)** | | |
| rSB02_312_23_5 | A (homozygous) | **1298 bp deletion** | | |
| rSB02_312_26_1 | A | **-64** | **+1** | **-28** |
| | B | **-1** | **+1** | **-28** |

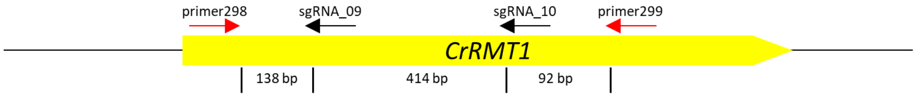


| Line | Allele | sgRNA_09 | sgRNA_10 |
|---|---|---|---|
| wt gDNA/rSB02_310 lines | A (homozygous) | | |
| rSB02_312_03_2 | A | **1kb insertion from 3' of RMT1** | -3 |
| | B | **+1-8** | -3 |
| rSB02_312_18_1 | A | **-10** | **-25** |
| | B | -27 | -72+9 |
| rSB02_312_23_5 | A | -27 | **-1** |
| | B | -27 | **-13** |
| rSB02_312_26_1 | A | -27 | **+1** |
| | B | -27 | **-8** |

**Figure S 10.** Characterization of CRISPR deletions in *CrJAM2*, *CrJAM3*, and *CrRMT1*

Overview of mutations detected in *C. roseus* hairy root lines with guides targeting *CrJAM2*, *CrJAM3,* and *CrRMT1*. Primers shown in red were used for cloning and sequencing of the genes. Additional internal primers were used for sequencing as needed. Major deletions and frame-shift mutations are bolded: rSB02_312 denotes lines transformed with pSB312 (Figure S 6), containing eight guides targeting *CrJAM2*, *CrJAM3,* and *CrRMT1*; rSB02_310 denotes lines transformed with pSB310 (Figure S 6), containing the *Cas9* and *HygR* (*hpt*) sequence but no guides; wt denotes gDNA from wild type seedlings. Lines rSB02_312_03_2 and rSB02_312_26_1 were pursued for RNA sequencing (Figure 5, Figure 6).

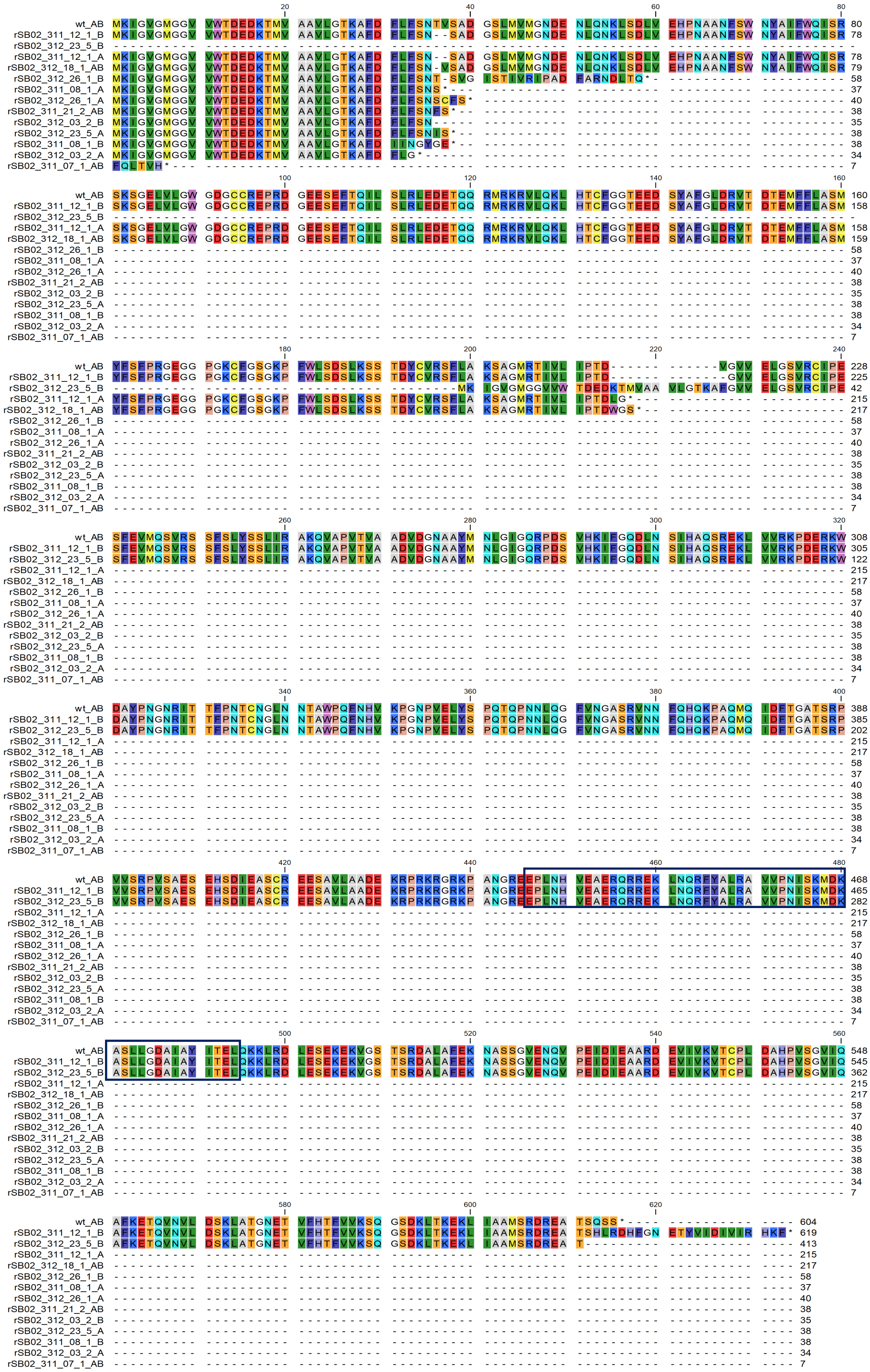

wt_AB MKIGVGMGGV VWTDEDKTMV AAVLGTKAFD FLFSNTVSAD GSLMVMGNDE NLQNKLSDLV EHPNAANFSW NYAIFWQISR 80
rSB02_311_12_1_B MKIGVGMGGV VWTDEDKTMV AAVLGTKAFD FLFSN--SAD GSLMVMGNDE NLQNKLSDLV EHPNAANFSW NYAIFWQISR 78
rSB02_312_23_5_B ----------
rSB02_311_12_1_A MKIGVGMGGV VWTDEDKTMV AAVLGTKAFD FLFSN--SAD GSLMVMGNDE NLQNKLSDLV EHPNAANFSW NYAIFWQISR 78
rSB02_312_18_1_AB MKIGVGMGGV VWTDEDKTMV AAVLGTKAFD FLFSN-VSAD GSLMVMGNDE NLQNKLSDLV EHPNAANFSW NYAIFWQISR 79
rSB02_312_26_1_B MKIGVGMGGV VWTDEDKTMV AAVLGTKAFD FLFSNT-SVG ISTIVRIPAD FARNDLTQ*- 58
rSB02_311_08_1_A MKIGVGMGGV VWTDEDKTMV AAVLGTKAFD FLFSNS*--- 37
rSB02_312_26_1_A MKIGVGMGGV VWTDEDKTMV AAVLGTKAFD FLFSNSCFS* 40
rSB02_311_21_2_AB MKIGVGMGGV VWTDEDKTMV AAVLGTKAFD FLFSNFS*-- 38
rSB02_312_03_2_B MKIGVGMGGV VWTDEDKTMV AAVLGTKAFD FLFSN----- 35
rSB02_312_23_5_A MKIGVGMGGV VWTDEDKTMV AAVLGTKAFD FLFSNITS*- 38
rSB02_311_08_1_B MKIGVGMGGV VWTDEDKTMV AAVLGTKAFD IINGYGE*-- 38
rSB02_312_03_2_A MKIGVGMGGV VWTDEDKTMV AAVLGTKAFD FLG*------ 34
rSB02_311_07_1_AB FQLTVH*--- 7
wt_AB SKSGELVLGW GDGCCREPRD GEESEFTQIL SLRLEDETQQ RMRKRVLQKL HTCFGGTEED SYAFGLDRVT DTEMFFLASM 160
rSB02_311_12_1_B SKSGELVLGW GDGCCREPRD GEESEFTQIL SLRLEDETQQ RMRKRVLQKL HTCFGGTEED SYAFGLDRVT DTEMFFLASM 158
rSB02_312_23_5_B ----------
rSB02_311_12_1_A SKSGELVLGW GDGCCREPRD GEESEFTQIL SLRLEDETQQ RMRKRVLQKL HTCFGGTEED SYAFGLDRVT DTEMFFLASM 158
rSB02_312_18_1_AB SKSGELVLGW GDGCCREPRD GEESEFTQIL SLRLEDETQQ RMRKRVLQKL HTCFGGTEED SYAFGLDRVT DTEMFFLASM 159
rSB02_312_26_1_B ---------- 58
rSB02_311_08_1_A ---------- 37
rSB02_312_26_1_A ---------- 40
rSB02_311_21_2_AB ---------- 38
rSB02_312_03_2_B ---------- 35
rSB02_312_23_5_A ---------- 38
rSB02_311_08_1_B ---------- 38
rSB02_312_03_2_A ---------- 34
rSB02_311_07_1_AB ---------- 7
wt_AB YFSFPRGEGG PGKCFGSGKP FWLSDSLKSS TDYCVRSFLA KSAGMRTIVL IPTD------ ------VGVV ELGSVRCIPE 228
rSB02_311_12_1_B YFSFPRGEGG PGKCFGSGKP FWLSDSLKSS TDYCVRSFLA KSAGMRTIVL IPTD------ -------GVV ELGSVRCIPE 225
rSB02_312_23_5_B ---------- ---------- ---------- ---------MK IGVGMGGVVW TDEDKTMVAA VLGTKAFGVV ELGSVRCIPE 42
rSB02_311_12_1_A YFSFPRGEGG PGKCFGSGKP FWLSDSLKSS TDYCVRSFLA KSAGMRTIVL IPTDLG*--- 215
rSB02_312_18_1_AB YFSFPRGEGG PGKCFGSGKP FWLSDSLKSS TDYCVRSFLA KSAGMRTIVL IPTDWGS*-- 217
rSB02_312_26_1_B ---------- 58
rSB02_311_08_1_A ---------- 37
rSB02_312_26_1_A ---------- 40
rSB02_311_21_2_AB ---------- 38
rSB02_312_03_2_B ---------- 35
rSB02_312_23_5_A ---------- 38
rSB02_311_08_1_B ---------- 38
rSB02_312_03_2_A ---------- 34
rSB02_311_07_1_AB ---------- 7
wt_AB SFEVMQSVRS SFSLYSSLIR AKQVAPVTVA ADVDGNAAYM NLGIGQRPDS VHKIFGQDLN SIHAQSREKL VVRKPDERKW 308
rSB02_311_12_1_B SFEVMQSVRS SFSLYSSLIR AKQVAPVTVA ADVDGNAAYM NLGIGQRPDS VHKIFGQDLN SIHAQSREKL VVRKPDERKW 305
rSB02_312_23_5_B SFEVMQSVRS SFSLYSSLIR AKQVAPVTVA ADVDGNAAYM NLGIGQRPDS VHKIFGQDLN SIHAQSREKL VVRKPDERKW 122
rSB02_311_12_1_A ---------- 215
rSB02_312_18_1_AB ---------- 217
rSB02_312_26_1_B ---------- 58
rSB02_311_08_1_A ---------- 37
rSB02_312_26_1_A ---------- 40
rSB02_311_21_2_AB ---------- 38
rSB02_312_03_2_B ---------- 35
rSB02_312_23_5_A ---------- 38
rSB02_311_08_1_B ---------- 38
rSB02_312_03_2_A ---------- 34
rSB02_311_07_1_AB ---------- 7
wt_AB DAYPNGNRIT TFPNTCNGLN NTAWPQFNHV KPGNPVELYS PQTQPNNLQG FVNGASRVNN FQHQKPAQMQ IDFTGATSRP 388
rSB02_311_12_1_B DAYPNGNRIT TFPNTCNGLN NTAWPQFNHV KPGNPVELYS PQTQPNNLQG FVNGASRVNN FQHQKPAQMQ IDFTGATSRP 385
rSB02_312_23_5_B DAYPNGNRIT TFPNTCNGLN NTAWPQFNHV KPGNPVELYS PQTQPNNLQG FVNGASRVNN FQHQKPAQMQ IDFTGATSRP 202
rSB02_311_12_1_A ---------- 215
rSB02_312_18_1_AB ---------- 217
rSB02_312_26_1_B ---------- 58
rSB02_311_08_1_A ---------- 37
rSB02_312_26_1_A ---------- 40
rSB02_311_21_2_AB ---------- 38
rSB02_312_03_2_B ---------- 35
rSB02_312_23_5_A ---------- 38
rSB02_311_08_1_B ---------- 38
rSB02_312_03_2_A ---------- 34
rSB02_311_07_1_AB ---------- 7
wt_AB VVSRPVSAES EHSDIEASCR EESAVLAADE KRPRKRGRKP ANGREEPLNH VEAERQRREK LNQRFVALRA VVPNISKMDK 468
rSB02_311_12_1_B VVSRPVSAES EHSDIEASCR EESAVLAADE KRPRKRGRKP ANGREEPLNH VEAERQRREK LNQRFVALRA VVPNISKMDK 465
rSB02_312_23_5_B VVSRPVSAES EHSDIEASCR EESAVLAADE KRPRKRGRKP ANGREEPLNH VEAERQRREK LNQRFVALRA VVPNISKMDK 282
rSB02_311_12_1_A ---------- 215
rSB02_312_18_1_AB ---------- 217
rSB02_312_26_1_B ---------- 58
rSB02_311_08_1_A ---------- 37
rSB02_312_26_1_A ---------- 40
rSB02_311_21_2_AB ---------- 38
rSB02_312_03_2_B ---------- 35
rSB02_312_23_5_A ---------- 38
rSB02_311_08_1_B ---------- 38
rSB02_312_03_2_A ---------- 34
rSB02_311_07_1_AB ---------- 7
wt_AB ASLLGDAIAY ITELQKKLRD LESEKEKVGS TSRDALAFEK NASSGVENQV PEIDIEAARD EVIVKVTCPL DAHPVSGVIQ 548
rSB02_311_12_1_B ASLLGDAIAY ITELQKKLRD LESEKEKVGS TSRDALAFEK NASSGVENQV PEIDIEAARD EVIVKVTCPL DAHPVSGVIQ 545
rSB02_312_23_5_B ASLLGDAIAY ITELQKKLRD LESEKEKVGS TSRDALAFEK NASSGVENQV PEIDIEAARD EVIVKVTCPL DAHPVSGVIQ 362
rSB02_311_12_1_A ---------- 215
rSB02_312_18_1_AB ---------- 217
rSB02_312_26_1_B ---------- 58
rSB02_311_08_1_A ---------- 37
rSB02_312_26_1_A ---------- 40
rSB02_311_21_2_AB ---------- 38
rSB02_312_03_2_B ---------- 35
rSB02_312_23_5_A ---------- 38
rSB02_311_08_1_B ---------- 38
rSB02_312_03_2_A ---------- 34
rSB02_311_07_1_AB ---------- 7
wt_AB AFKETQVNVL DSKLATGNET VFHTFVVKSQ GSDKLTKEKL IAAMSRDREA TSQSS*---- ---------- ---- 604
rSB02_311_12_1_B AFKETQVNVL DSKLATGNET VFHTFVVKSQ GSDKLTKEKL IAAMSRDREA TSHLRDHFGN ETYVIDIVIR HKF* 619
rSB02_312_23_5_B AFKETQVNVL DSKLATGNET VFHTFVVKSQ GSDKLTKEKL IAAMSRDREA T--------- ---------- ---- 413
rSB02_311_12_1_A ---------- 215
rSB02_312_18_1_AB ---------- 217
rSB02_312_26_1_B ---------- 58
rSB02_311_08_1_A ---------- 37
rSB02_312_26_1_A ---------- 40
rSB02_311_21_2_AB ---------- 38
rSB02_312_03_2_B ---------- 35
rSB02_312_23_5_A ---------- 38
rSB02_311_08_1_B ---------- 38
rSB02_312_03_2_A ---------- 34
rSB02_311_07_1_AB ---------- 7

**Figure S 11.** Predicted CrJAM2 amino acid sequences in the CRISPR-Cas-deletion lines. The predicted amino acid sequences of CrJAM2 for all the double mutant and triple mutant lines from Figures S 9 and S 10 are aligned and its bHLH domain outlined in black.

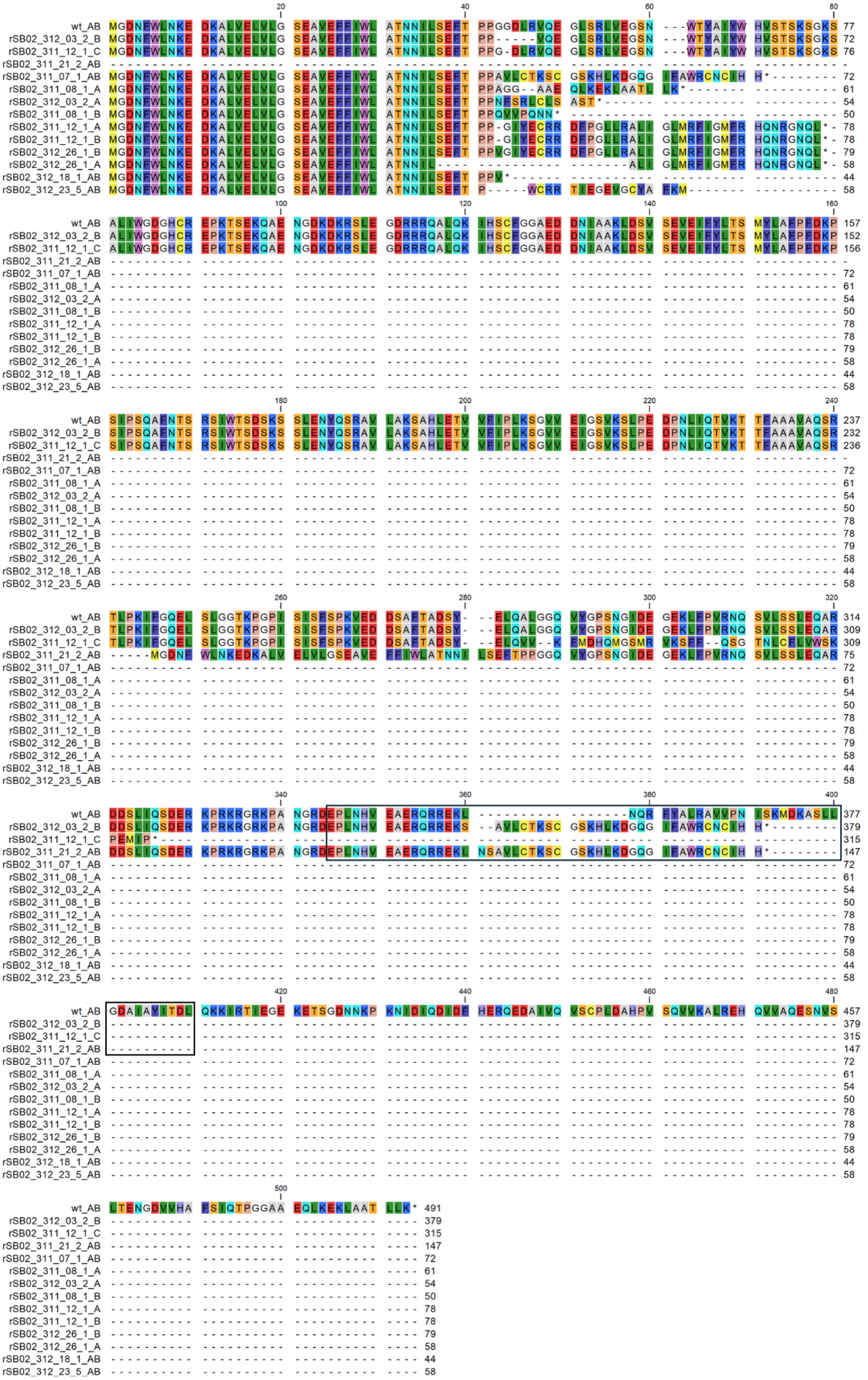

**Figure S 12.** Predicted CrJAM3 amino acid sequences in the CRISPR-Cas-deletion lines. The predicted amino acid sequences of CrJAM3 for all the double mutant and triple mutant lines from Figures S 9 and S 10 are aligned and its bHLH domain outlined in black.

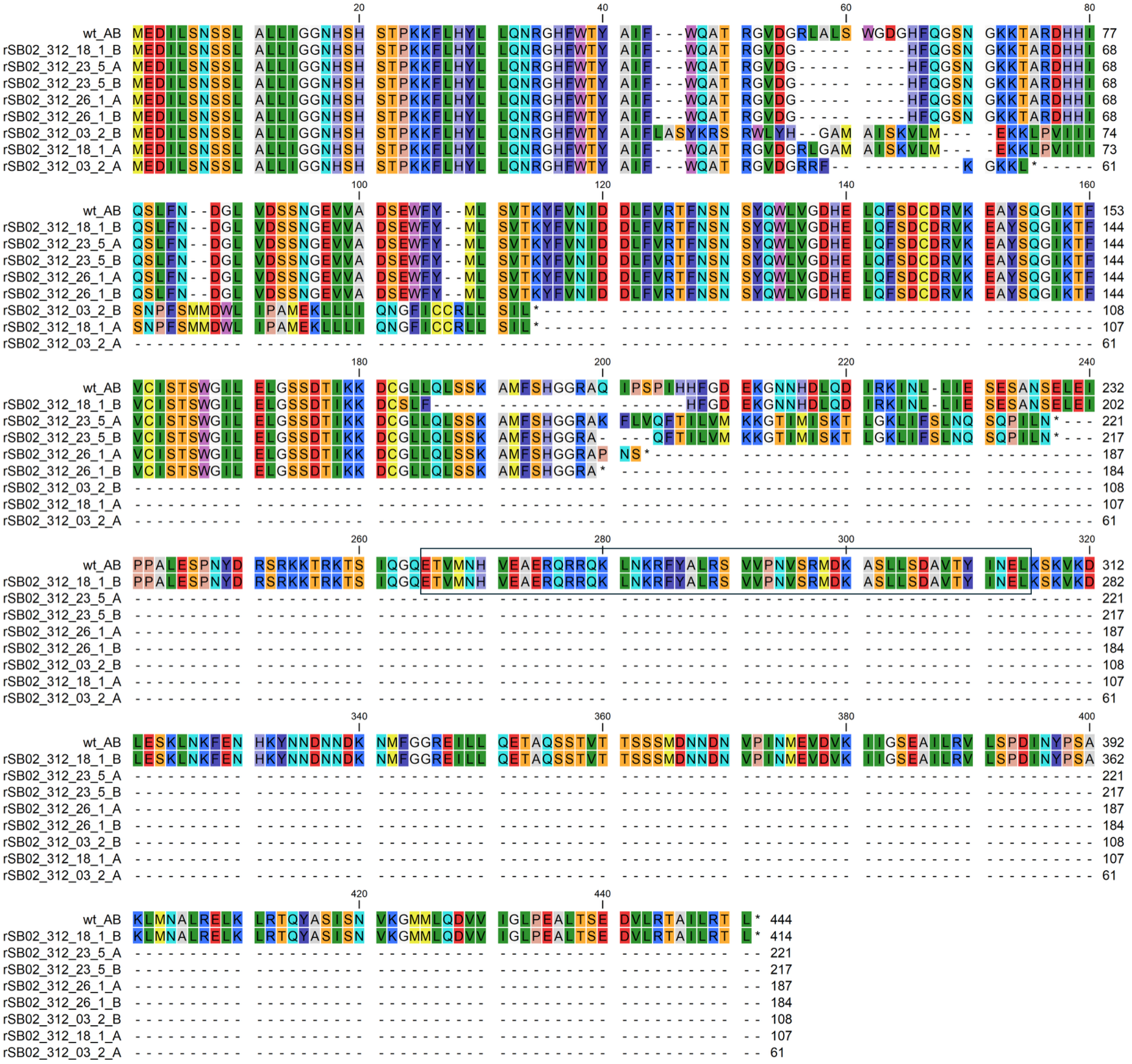

**Figure S 13.** Predicted CrRMT1 amino acid sequences in the CRISPR-Cas-deletion lines. The predicted amino acid sequences of CrRMT1 for all the double mutant and triple mutant lines from Figures S 9 and S 10 are aligned and its bHLH domain outlined in black.

### Supplementary Tables

**Table S 1.** Identification of potential bHLH IIId JAM-like transcription factors in *C. roseus*

| BlastP input sequence | *C. roseus* version 3 BlastP hit | Protein | E-value | % Identity |
|---|---|---|---:|---:|
| AtbHLH17 (AtJAM1) | Hit 1: CRO_06G033380 | CrJAM2 | 0.00E+00 | 49% |
| NC_003071 | Hit 2: CRO_04G005850 | CrJAM3 | 1.31E-103 | 37% |
| | Hit 3: CRO_07G000280 | CrMYC2a | 4.84E-44 | 47% |
| | Hit 4: CRO_06G033750 | CrMYC2c | 1.42E-30 | 40% |
| AtbHLH13 (AtJAM2) | Hit 1: CRO_06G033380 | CrJAM2 | 0.00E+00 | 52% |
| NM_001197954 | Hit 2: CRO_04G005850 | CrJAM3 | 2.02E-45 | 50% |
| | Hit 3: CRO_07G000280 | CrMYC2a | 4.68E-31 | 42% |
| | Hit 4: CRO_08G004450 | CrRMT1 | 7.52E-22 | 34% |
| AtbHLH03 (AtJAM3) | Hit 1: CRO_04G005850 | CrJAM3 | 4.66E-159 | 48% |
| NC_003075 | Hit 2: CRO_06G033380 | CrJAM2 | 6.98E-55 | 55% |
| | Hit 3: CRO_07G000280 | CrMYC2a | 4.15E-35 | 40% |
| | Hit 4: CRO_08G004450 | CrRMT1 | 2.52E-29 | 28% |
| AtbHLH14 | Hit 1: CRO_08G004450 | CrRMT1 | 3.79E-46 | 33% |
| NC_003075 | Hit 2: CRO_07G000280 | CrMYC2a | 1.56E-31 | 47% |
| | Hit 3: CRO_06G033750 | CrMYC2c | 6.84E-28 | 45% |
| | Hit 4: CRO_06G033380 | CrJAM2 | 2.38E-23 | 43% |

**Table S 2.** *C. roseus* promoter constructs used for testing repression by CrJAM2 and CrJAM3

| Promoter and 5'UTR | L0 vector (Pro+5U denotes promoter plus the 5'UTR) | Size in bp (upstream of ATG) |
|---|---|---|
| pBIS1 | pSB219 - pL0_pBIS1 (Pro+5U) | 2016 |
| pBIS2 | pSB225 - pL0_pBIS2 (Pro+5U) | 1960 |
| pG8O | pSB94 - pL0_pG8O (Pro + 5U) | 1000 |
| pGO | pSB245 - pL0_pGO (Pro+5U) | 1643 |
| pLAMT | pSB235 - pL0_pLAMT (Pro+5U) | 1998 |
| pMYC2a | pSB290 - pL0_pMYC2a (Pro+5U) | 1291 |
| pORCA3 | pSB190 - pL0_pORCA3 (Pro + 5U) | 817 |
| pSTR1 | pSB62 - pL0_pSTR (Pro + 5U) | 586 |
| pT16H1 | pSB248 - pL0_pT16H1 (Pro+5U) | 1675 |
| pT16H2 | pSB278 - pL0_pT16H2 (Pro + 5U) | 1378 |
| pT19H | pSB124 - pL0_pT19H (Pro + 5U) | 1007 |
| pZCT1 | pSB84 - pL0_pZCT1 (Pro+5U) | 1000 |

**Table S 3.** Oligonucleotides used in cloning promoters and coding sequences, PCR, or qPCR

| Name | Sequence (5'-3') | Note |
|---|---|---|
| **Cloning of promoters** | | |
| pBIS1_F_1 | aaGGTCTCAACATGGAGTCATACTGCTATGGAGTAAATGG | cloning of *pBIS1* |
| pBIS1_R_1 | ttGGTCTCTACAACACTTTTATCAAATGAGTCCTCAG | |
| pBIS1_F_2 | AAGGTCTCaACATAGTGAAAAGTGATTTGAAATCTAATG | |
| pBIS1_R_2 | ttGGTCTCtACAAGTGAGTCCTGATTTCTTGTACG | |
| pBIS1_F_3 | aaGGTCTCaACATTCACATAGGAGAGATTGAAATTG | |
| pBIS1_R_3 | ttGGTCTCtACAACATTTTTAGATGAGCTCCAACTAAAC | |
| pBIS2_F | ttgaagacaaGGAGCATAGTGGTACAGATAGTATACAGATTATCC | cloning of *pBIS2* |
| pBIS2_R | ttgaagacaaCATTTCCAAAGGCAAATGAAGAAG | |
| G10H-1000_F | ttgaagacaaGGAGATCAAACCCTTGCTCTTCCTTATATATAGAG | cloning of *pG8O* (also called *pG10H*) |
| G10H-1_R | ttgaagacaaCATTGGAATGGAAGTGTACAATTTGTTCTC | |
| pGO_F1_Frag1 | aaGGTCTCaACATGGAGCGTAATGAATTTAGTGACGGATCG | cloning of *pGO* |
| pGO_R1_Frag1 | ttGGTCTCtACAACAGTGGATTGCAAGCTGAGC | |
| pGO_F2_Frag2 | aaGGTCTCaACATACTGTTATGAGCAGAGAAACTGTGGTAC | |
| pGO_R2_Frag2 | ttGGTCTCtACAACATTGTGTTTTGTTCAAGAGAGGAG | |
| pLAMT_F_1 | aaGGTCTCaACATGGAGGCAAAAGAAAGAGAGAAAG | cloning of *pLAMT* |
| pLAMT_F_1 | ttGGTCTCtACAATGACTTCACTTCATGAATTAGTAGTACG | |
| pLAMT_F_2 | aaGGTCTCaACATGTCAAAAAGATTAATAAAAATGTGAACG | |
| pLAMT_R_2 | ttGGTCTCtACAATCTCGAAATCGAGCAACAG | |
| pLAMT_F_3 | aaGGTCTCaACATGAGACTTCAGTTACAATGGTCAAATATG | |
| pLAMT_R_3 | ttGGTCTCtACAAGCTTAATATATTTGGAAGTCTTTTGTATTAAG | |

| | | |
|---|---|---|
| pLAMT_F_4 | aaGGTCTCaACATAAGCCCGTAATTTATATATAACTCTTTCTTTATTAG | |
| pLAMT_R_4 | ttGGTCTCtACAACATTTTTCTTTCTTCTCTGTATGAGTTTTTTG | |
| 213_pMYC2a_frag_1 | aaGGTCTCaACATGGAGATAGCATATAATTACTCTCTAGTTGTCTCGAC | cloning of *pMYC2a* |
| pMYC2a_R_1 | ttGGTCTCtACAAAGTCTGTTTGAGTGAGTGAAAAATTG | |
| pMYC2a_F_2 | aaGGTCTCaACATGACTTCTTCTTCTCGAATTCAACAATC | |
| 235_pMYC2a_frag2.1R | ttGGTCTCtACAAAAGAATCAAAAAGGGTTCTTGTTTCTG | |
| 236_pMYC2a_frag_2.2F | ACATTCTTCAATTTCTTTTTTTTTTTTTGTTTGATGGAATG | |
| 237_pMYC2a_frag_2.2R | ACAACATTCCATCAAACAAAAAAAAAAAAAGAAATTGAAGA | |
| Porca3_R | ttgaagacaaCATTTTTCTTCTTTTTTAGAAACTGATTTTTG | cloning of *pORCA3* |
| Porca3_F3 | ttgaagacaaGGAGTGTCCCAAGTAAATAAAAACGGAGAG | |
| STR_pro_F | ttgaagacaaGGAGGATCTTTATTGTTTGTATTTTCATAATTAATTAATCCC | cloning of *pSTR1* |
| STR_pro_R | ttgaagacaaCATTGTTGACTGTAAATGATGGGAGTG | |
| pT16H1_F1_frag1 | aaGGTCTCaACATGGAGCCGGGTAGGCAGTTTGATTG | cloning of *pT16H1* |
| pT16H1_R1_frag1 | ttGGTCTCtACAACGATTTTTTCCTCACGTGGG | |
| pT16H1_F2_frag2 | aaGGTCTCaACATATCGCAAAACACAATCAAAGC | |
| pT16H1_R1_frag2 | ttGGTCTCtACAACATTAGTGAAAATTATGCTTGTTATTGATTTGC | |
| pT16H2_F1_65 | aaGGTCTCaACATGGAGAGTAATCTCATTGATGTGATGGGC | cloning of *pT16H2* |
| pT16H2_R2_64 | ttGGTCTCtACAACATTAGGTAGAGAAAGTTGACCAAAATC | |
| T19H_F | ttgaagacaaGGAGCCATGAGGTAGAAATTATGAAAGAAATTAAGG | cloning *pT19H* |
| T19H_R | ttgaagacaaCATTATTTTTCTTGCTAGAGAAATCTAGTTGTTTTTGC | |
| Pzct1-1000_F | ttgaagacaaGGAGTGTCTCATTTTCGGAGTTTAGCTTGC | cloning *pZCT1* |

| | | |
|---|---|---|
| Pzct1-1_R | ttgaagacaaCATTGTTTTTTGTT ATTGAATTGAAACTTGCGAC | |
| **Cloning of coding sequences** | | |
| CrJAM2_F_1 | ttgaagacaaAATGAAAATAGGAG TCGGTATGGG | cloning *CrJAM2* CDS |
| CrJAM2_R_1 | ttgaagacaaAAGATCTAACGGAC TGCATGACC | |
| CrJAM2_F_2 | ttgaagacaaTCTTCCTTCTCATT GTATTCGTCAC | |
| CrJAM2_R_2 | ttgaagacaaAAGCTCACGAAGAT TGTGAGG | |
| JAM3_F_1 | ACATaATGGGGGATAACTTTTGGT TAAATAAAGAAGATAAG | cloning *CrJAM3* CDS |
| JAM3_R_1 | ACAACTTATCTTCTTTATTTAACC AAAAGTTATCCCCCATt | |
| JAM3_F_2 | aaGGTCTCaACATTAAGGCTTTGG TTGAGCTTGTATTG | |
| JAM3_R_2 | ttGGTCTCtACAATGCGCCGATCT CCTTCCAACG | |
| JAM3_F_3 | aaGGTCTCaACATCGCAGACAGGC ACTTCAGAAGATTC | |
| JAM3_R_3 | ttGGTCTCtACAAGATCTTCAGGG AGCGACTTTACTG | |
| JAM3_F_4 | aaGGTCTCaACATGATCCTAATTT GATTCAGACAGTTAAAACTAC | |
| JAM3_R_4 | ttGGTCTCtACAAAAGCTCATTTT AAAAGCGTAGCAGC | |
| **Confirmation of hairy roots** | | |
| VirD2F1 | ATGCCCGATCGAGCTCAAGT | *virD2*; elimination of *Agrobacterium* (Haas et al., 1995) |
| VirD2R224 | TCGTCTGGCTGACTTTCGTCATAA | |
| rolC_F | CAACCTGTTTCCTACTTTGTTAAC | integration of *rolC*; confirmation of hairy roots (Suttipanta et al., 2011) |
| rolC_R | AAACAAGTGACACACTCAGCTTC | |
| HygR_F | tgacctattgcatctcccgccg | integration of *HygR* / *hpt* (T-DNA LB) |
| HygR_R | tccatacaagccaaccacggcc | |
| GUS(328)F | GGCTATACGCCATTTGAAGC | integration of *GUS* (in rSB02_299 lines) |
| GUS(1059)R | AGAGGTTAAAGCCGACAGCA | |
| P2x35S_F | GAGGACACGCTCGAGTATAAGAGC | integration of *CaMV 2x35S::CrJAM2* (in rSB02_297 lines) |
| sgRNA_05_F_SM | attgACCGTCAGCTGAAACAGTAT | |
| P2x35S_F | GAGGACACGCTCGAGTATAAGAGC | integration of *CaMV 2x35S::CrJAM3* (in rSB02_298 lines) |
| sgRNA_01_F_SM | attgACTCGTAAATCCCCACCAGG | |

| | | |
|---|---|---|
| **RT-qPCR** | | |
| JAM2q_set3F | GTTGCAGAGAACCTAGAGATGG | *CrJAM2* expression |
| JAM2q_set3R | TCTGTTCCACCAAAGCATGTA | |
| JAM3q_set3F | CAATCTAGAACGCTCCCAAAGA | *CrJAM3* expression |
| JAM3q_set3R | CAATTCATACGAGTCCGCAGTA | |

**Table S 4.** Oligonucleotides used for the creation and confirmation of CRISPR-Cas HR lines

| Oligonucleotide name | oligonucleotide (5' - 3') | Note |
|---|---|---|
| sgRNA_01_F | attgACTCGTAAATCCCCACCAGG | guide for *CrJAM3* CDS |
| sgRNA_01_R | aaacCCTGGTGGGGATTTACGAGT | |
| sgRNA_02_F | attgTATGAATTGCAAGCCTTAGG | guide for *CrJAM3* CDS |
| sgRNA_02_R | aaacCCTAAGGCTTGCAATTCATA | |
| sgRNA_03_F | attGGCGTGAAAAGTTAAACCAG | guide for *CrJAM3* CDS |
| sgRNA_03_R | aaacCTGGTTTAACTTTTCACGC | |
| sgRNA_04_F | attgTTCTCCATCCAAACTCCAGG | guide for *CrJAM3* CDS |
| sgRNA_04_R | aaacCCTGGAGTTTGGATGGAGAA | |
| sgRNA_05_F | attgACCGTCAGCTGAAACAGTAT | guide for *CrJAM2* CDS |
| sgRNA_05_R | aaacATACTGTTTCAGCTGACGGT | |
| sgRNA_06_F | attGAGGCAGTAAGAGTCGAGAT | guide for *CrJAM2* 5' |
| sgRNA_06_R | aaacATCTCGACTCTTACTGCCT | |
| sgRNA_07_F | attGCTGATTCCTACTGATGTCG | guide for *CrJAM2* CDS |
| sgRNA_07_R | aaacCGACATCAGTAGGAATCAG | |
| sgRNA_08_F | attgAAATGATCACGAAGATTGTG | guide for *CrJAM2* CDS |
| sgRNA_08_R | aaacCACAATCTTCGTGATCATTT | |
| sgRNA_09_F | attgCGCCCCATGATAAAGCGAGA | guide for *CrRMT1* CDS |
| sgRNA_09_R | aaacTCTCGCTTTATCATGGGGCG | |
| sgRNA_10_F | attgTTGGACTAGGAATTTGGGCA | guide for *CrRMT1* CDS |
| sgRNA_10_R | aaacTGCCCAAATTCCTAGTCCAA | |
| primer294 | aaGGTCTCaACATTTCTTCTGGTATGTGTATATGTTCTTCC | *CrJAM2* cloning/sequencing |
| primer295 | ttGGTCTCtACAAGTATACCAAAATCTCTGCAGAAGC | |
| primer296 | aaGGTCTCaACATGTTGAATAGTTCAAGGATGGG | *CrJAM3* cloning/sequencing |
| primer297 | ttGGTCTCtACAACCATCTATTTTTCCTCGGC | |
| primer298 | aaGGTCTCaACATGGAGGACATTCTTTCCAATTC | *CrRMT1* cloning/sequencing |
| primer299 | ttGGTCTCtACAATTCAGAATTGGCTGACTCTG | |
| VirD2F1 | ATGCCCGATCGAGCTCAAGT | *virD2*; elimination of *Agrobacterium* (Haas et al., 1995) |
| VirD2R224 | TCGTCTGGCTGACTTTCGTCATAA | |
| rolC_F | CAACCTGTTTCCTACTTTGTTAAC | integration of *rolC*; confirmation of hairy roots (Suttipanta et al., 2011) |
| rolC_R | AAACAAGTGACACACTCAGCTTC | |
| HygR_F | tgacctattgcatctcccgccg | integration of *HygR* / *hpt* (T-DNA LB) |
| HygR_R | tccatacaagccaaccacggcc | |

| | | |
|---|---|---|
| primer41 | ttgaagacaaGGAGaggcgggaaacgacaatctg | integration of sgRNA region (T-DNA RB) |
| seqTU5 (JS838) | GCCAGCTTCTATGAGTACTGA | |
| primer41 | ttgaagacaaGGAGaggcgggaaacgacaatctg | integration of promoter - *Cas9* region (T-DNA RB) |
| primer283 | GTATTCATCGGTGATAACGGC | |

**Table S 5.** Evaluation of guide sequences

| protospacer | 5'-target-3' | PAM | on-target score** | bp identity to closest off-target |
|---|---|---|---|---|
| sgRNA_01 | ACTCGTAAATCCCCACCAGG | CGG | 76 | 15/20 |
| sgRNA_02 | TATGAATTGCAAGCCTTAGG | TGG | 64 | 15/20 |
| sgRNA_03 | GGCGTGAAAAGTTAAACCAG | CGG | 72 | 16/20 |
| sgRNA_04 | TTCTCCATCCAAACTCCAGG | TGG | 62 | 16/20 |
| sgRNA_05 | ACCGTCAGCTGAAACAGTAT | TGG | 53 | 16/20 |
| sgRNA_06 | GAGGCAGTAAGAGTCGAGAT | TGG | 55 | 15/20 |
| sgRNA_07 | GCTGATTCCTACTGATGTCG* | GGG | 64 | 15/20 |
| sgRNA_08 | AAATGATCACGAAGATTGTG | AGG | 65 | 16/20 |
| sgRNA_09 | CGCCCCATGATAAAGCGAGA | CGG | 64 | no off-target blast hits |
| sgRNA_10 | TTGGACTAGGAATTTGGGCA | CGG | 60 | 16/20 |

*red: 1 bp mismatch between guide and target sequence (genomic target is T instead of C). The mismatch is due to a difference in the sequence between the cultivar used in our experiments and the sequenced genome.

**On-target score prediction in Benchling (Doench et al., 2016)

**Table S 6.** Coding sequences of *CrJAMs* and promoter plus 5'UTR sequences of MIA pathway genes and TF genes

Highlighted in red are former BpiI/Eco31I/Esp3I recognition sites that were removed by replacing with a synonymous codon (in CDS) or 1 bp A to T or T to A mutations (in promoters and 5'UTRs) for compatibility with MoClo cloning. Highlighted in the promoters and 5'UTRs: blue are G-boxes (CACGTG), green are G-like boxes (CATGTG), grey are T/G-boxes (AACGTG), pink are G/A-boxes (CACGAG), dark green are G/C-boxes (CACGCG).

>*CrJAM2* CDS
ATGAAAATAGGAGTCGGTATGGGAGGTGTTGTTTGGACTGATGAAGATAAAACAA
TGGTGGCGGCGGTTTTGGGGACTAAGGCTTTTGATTTCTTGTTTTCCAATACTGTTT
CAGCTGACGGTTCATTAATGGTTATGGGGAATGATGAAAATTTGCAGAATAAGCT
TTCAGATCTCGTTGAACATCCAAACGCTGCCAATTTTAGTTGGAATTATGCAATTT
TCTGGCAAATTTCGCGGTCCAAGTCCGGCGAATTGGTATTAGGATGGGGTGATGG
CTGTTGCAGAGAACCTAGAGATGGGGAGGAATCCGAATTTACCCAAATTTTAAGT
CTCCGGCTTGAGGATGAAACTCAACAGAGGATGAGGAAAAGAGTTCTTCAGAAAC
TGCATACATGCTTTGGTGGAACAGAAGAGGATAGTTATGCTTTTGGATTGGATAG
GGTTACTGATACTGAGATGTTCTTCCTGGCATCCATGTACTTTTCATTTCCTCGAGG
AGAAGGAGGTCCGGGGAAGTGTTTTGGATCTGGGAAGCCTTTTTGGTTATCTGATT
CATTGAAGTCATCTACAGATTATTGTGTTAGGTCATTTCTAGCGAAGTCGGCTGGT
ATGCGAACTATTGTGCTGATTCCTACTGATGTTGGGGTAGTTGAATTAGGTTCAGT
TAGATGCATTCCGGAAAGTTTTGAGGTCATGCAGTCCGTTAGATCTTCCTTCTCAT
TGTATTCGTCACTAATTAGGGCCAAACAAGTTGCACCTGTGACAGTGGCAGCCGA
CGTGGATGGCAATGCAGCTTACATGAATTTGGGGATAGGGCAACGGCCAGACAGC
GTTCACAAGATTTTTGGTCAAGATTTGAATTCAATTCATGCTCAAAGTCGGGAAAA
GCTTGTCGTCAGAAAGCCAGACGAAAGAAAGTGGGATGCTTATCCAAATGGAAAC
AGAATTACTACGTTTCCAAACACGTGCAATGGTCTGAATAACACCGCGTGGCCAC
AGTTCAATCATGTCAAGCCAGGCAATCCCGTGGAACTTTATAGTCCCCAGACACA
ACCAAACAATTTACAGGGGTTTGTTAATGGAGCAAGTCGTGTAAACAATTTTCAG
CATCAAAAGCCAGCTCAAATGCAAATTGACTTTACAGGAGCGACATCAAGACCTG
TAGTTTCAAGACCTGTAAGTGCTGAGTCTGAGCATTCAGATATTGAGGCTTCATGC
AGAGAAGAGAGTGCAGTTCTAGCTGCTGATGAAAAGAGGCCTCGGAAGCGTGGT
CGGAAACCTGCCAATGGAAGAGAAGAGCCCCTCAATCATGTCGAGGCAGAGAGG
CAGCGAAGAGAGAAGCTGAATCAGCGGTTTTACGCACTGAGAGCTGTTGTACCCA
ATATTTCTAAAATGGATAAAGCATCCCTTTTGGGAGATGCCATTGCTTACATAACC
GAGCTGCAGAAGAAGCTTAGGGACCAGGAATCAGAAAAGGAGAAAGTGGGAAGC
ACCTCAAGAGATGCATTAGCTTTTGAGAAGAACGCGAGTTCTGGCGTTGAAAATC
AAGTGCCTGAAATCGATATTGAAGCCGCACGTGATGAAGTTATTGTGAAGGTGAC
CTGTCCATTGGATGCTCATCCTGTATCTGGAGTCATTCAAGCATTCAAAGAAACAC
AGGTGAACGTCTTGGATTCAAAACTGGCCACAGGTAATGAAACAGTGTTTCACAC

ATTTGTTGTCAAGTCTCAAGGATCAGACAAGTTAACAAAGGAAAAGTTGATAGCA
GCAATGTCCCGAGATCGTGAGGCAACCTCACAATCTTCGTGA
>*CrJAM3* CDS
ATGGGGGATAACTTTTGGTTAAATAAAGAAGATAAGGCTTTGGTTGAGCTTGTATT
GGGCAGTGAAGCTGTTGAATTTTTCATTTGGTTGGCTACAAATAACATACTGTCCG
AATTTACACCGCCTGGTGGGGATTTACGAGTGCAGGAGGGACTTTCCAGGCTTGTT
GAGGGCTCTAATTGGACTTATGCGATTTATTGGCATGTTTCGACATCAAAATCGGG
GAAATCAGCTTTAATATGGGGTGATGGGCATTGTAGGGAACCCAAGACAAGTGAG
AAGCAGGCTGAAAATGGTGATAAAGATAAAAGATCGTTGGAAGGAGATCGGCGC
AGACAGGCACTTCAGAAGATTCATTCTTGTTTTGGGGGAGCAGAGGATGATAACA
TTGCTGCGAAGCTCGATTCTGTTTCAGAAGTGGAGATTTTTTATCTAACTTCTATGT
ATTTAGCATTTCCATTTGACAAGCCTTCTATTCCTTCACAGGCTTTTAATACTAGTA
GATCGATTTGGACTTCTGATAGCAAGAGTAGTTTAGAAAATTACCAATCCAGAGC
CGTTCTAGCAAAGTCGGCTCATCTTGAGACAGTAGTGTTTATACCGCTGAAATCAG
GAGTAGTGGAGATTGGTTCAGTAAAGTCGCTCCCTGAAGATCCTAATTTGATTCAG
ACAGTTAAAACTACTTTTGCAGCTGCTGTTGCCCAATCTAGAACGCTCCCAAAGAT
TTTCGGCCAAGAACTTAGCCTTGGAGGTACCAAGCCAGGTCCAATTAGTATTAGTT
TTTCTCCAAAGGTAGAAGATGATTCAGCTTTTACTGCGGACTCGTATGAATTGCAA
GCCTTAGGTGGTCAAGTTTATGGACCATCAAATGGGATCGATGAGGGTGAAAAGC
TTTTTCCAGTCAGGAACCAATCTGTGCTTTCTAGTTTGGAGCAAGCCAGAGATGAT
TCCTTGATCCAGTCTGATGAGAGGAAACCTAGAAAGAGAGGTAGAAAGCCAGCC
AATGGCCGGGATGAACCTCTGAATCATGTGGAAGCAGAGAGGCAGAGGCGTGAA
AAGTTAAACCAGCGGTTTTATGCACTAAGAGCTGTGGTTCCAAACATCTCAAAGA
TGGACAAGGCATCTTTGCTTGGCGATGCAATTGCATACATCACTGATCTTCAGAAG
AAGATAAGGACTATTGAAGGTGAGAAGGAGACTAGTGGGGATAATAATAAGCCA
AAGAATATTGATATCCAAGATATTGATTTTCATGAAAGGCAAGAAGATGCAATTG
TACAAGTTAGCTGTCCGTTAGATGCTCATCCAGTTTCTCAAGTTGTAAAGGCACTG
AGGGAGCATCAAGTTGTAGCACAAGAATCTAATGTTTCTCTGACCGAAAATGGGG
ATGTTGTTCACGCCTTCTCCATCCAAACTCCAGGTGGTGCCGCAGAACAATTGAAG
GAGAAGTTGGCTGCTACGCTTTTAAAATGA
>*BIS1* promoter and 5’UTR
TCATACTGCTATGGAGTAAATGGAAAAAAAAAAACTTAAAATTCTTATTTTCTCAT
AAAACAAAAATGATCAAACGAGAAACAAGTATTTAAAAGAAAAGAAGTGATCAT
TCTTTTAAAAGGAGGGGTGGTTATTTTATATGTACATTTTTTATTTTAAAGAAAAA
AGAATTATATATTATTAAAATTCAATAAAAATAGCTTGAGATATATCAACGGTAA
GTTTCTAGTCCAAACCTAACAGGACTATCAGAATTTGAGGCGAGTTTTACTAGAA
GATGGTTCGAACTATTACCAAATCTATACACATATTTTTATACGTACATCTTTTGTA
CATGATCTTTTCTATATTTATTTAAAATTTTGTTTTTTTATTAATTTTCTGAGGACTC
ATTTGATAAAAGTGAAAAGTGATTTGAAATCTAATGTTTGAACTCTAAATTTGAAA
ATTAGAATCTGAATTAATTAATTAGTGTTAATTATATATATACATAAAATATTTAA
GAGTTCATAGTTAAATTACGATTTTTTACTTAAAATTTATAATTATAATTGCACATC
TTAAACAATTAGGTGATTTGTAAAAATTATTATCAAACACATTTAAAATATTAAAT
ATTTAAAAATTATTAATTATTAAACCTATTTAATTTAATAAATTCTTAAGTAATTTA
TTTTAAAACCAGGAAAGCTCTGATACTAAACATCAACGAGGGTCCGGCCCTTTAT
CCCTCTCCTATTTTCAATTATATTCAAGTATTTAATTTGACTTCCTCTTAGCTGTTAT
GCTCAGGCCTTGGCCTTCCAGGAACCATCCTAAAGTATATCTTCCTTCCGATTTGA

TTATACATGATATAAACTTTGAACATGTTTGTATGTGTGGGCGGCCGGAGATATAG
AAACTGACTTCCCAAAGGCACTCATCCGAGTTATGGGACTGGATGGAAGTGAACA
TTACTTAAAGGGCTTTCACGACTTCACTAGATTGCTTAATTTCTATCCAACTGTAC
AACATTTAAGTAGAATATATAAATTATGCTGATTCAATATTTCTGCAGATCGGAGG
CCTAAAATTTGGTGTGCATAAGGAATCAACGGTTATGCAAATTAAAGATCGAAGA
TCTAATGCGTACAAGAAATCAGGACTCACATAGGAGAGATTGAAATTGCTATATT
TTTCCTCTATCAAATCTTGAAAACTTTATATATATATATATATATATATGAATTAGG
ATATTTGTTCATTGTTAATTTGGCTAAACAAAAAACTGATGACGTTGAATATAGAG
GGAAGGAAGGGTTAAGAGTTTATATTATCCACTACTTATCACAGACATCATCATA
GACTAGTGTGATACTACTATAAGTAAAAGAATTTTCAAAATAATGGTACTGTAAC
GTCAAAGAATGATTTCTAATTTTTAATCCCAATTAATTTCTTTTGGGTGAATAACA
ACAAAAGGAAAATGAAAGCAAAAGAAAAAAGGATTGTTTAAATAAATGGTGCAT
TATTAATTCCACAACAAGTAAAGTAACGACGAATTTGAAATATTTGGAATGTAGT
GATAGTGATGATGATGAGAGGAGAGTTTGTGTGTGTGTGTAAGGCACATGGCATG
CGCTCAGCCTGAAGATTTAAACCACAACACAACAGCAACCGCAACCGCAGAGTAT
ATATTTGGAGAGAGAAGTGTTTCCGCTCCCTTCTTCGCCTTCTGCGCTCCCTCTAA
GTTGGATGCCTACATTTCTACTACGGAGCGCTGTAGGCTTCTTTTGTAGCGAATTT
ATCACATTTTCCAATTATTGATTTCCCTATTTAAACATCACCAAAACTTTCTTACTT
TCCCCCACCATCCACTTCCCGGAGCTCTTCTTCAGACAAGTTCACGCTCCATCCCA
GCTTTTCCTCTTCCATTCTCTTTCATCTCCTTTTTCGTTATTCTCTCCTCGTTCTTCAT
TTTAATTACCTTTTTCTCCTCCGATTTCACTGAGAGGTTTAGTTGGAGCTCATCTAA
AAATG
>*BIS2* promoter and 5’UTR
CATAGTGGTACAGATAGTATACAGATTATCCGAACCCATACCAGAGAATCTTTGG
GATTCATACAATGATTTTTATTTTTGGCAAGATATAATTATTTATTCTTTCTTTAAT
TTCCTTTATCAATTAATCTTAGAAAAAAATTATTTCCTTTTTCTTTTAATTCACTTTT
TCCACTAGCACCATTGGAACATTTTATCTAACATCTCCTATAATATTTACTTTTTAT
AATATATTTTCTTCTTGTGGTTTATTTTTTCCTTATAAGATTTATATTTTATTTGATA
AAATTTATTAAGAGAAATTAATTATAAATACCTAAAACATTTTTCGATTTTCTCTTT
TTACCATAAATATTTTATTTTAACTCTTTATGCCCTCAAACATTTTTAAATATGCTT
ATTATTATAGTTATGGTGAAGGATAGCAGAATGCATGGTGGTAAGCATGTCTCAC
TCATGCTCACACTAAAAAAAGCCAAATTAATGTTTTGTTTTAATTTTTTATGCCCTC
GAACATTTTTTAATTGCATATTTTTATAATTATAATGAAGGATAACAGAATCTGAT
GGCGATGAACATGGCAAAATGCAAACTAAAAAAAAAAGTCAAAAATTGACATTT
TTTAGTGTGCAGGTGAGTGAGGCGTGCTCATTGCTACCGAATTTCGATGTCCTTCA
CCATAACTATAATAATATGCAAATCAAAAAATGTTTTGAATGTATAAAGAGCTAA
ATATTTTATGTTTGAGGGTATAAAGAGTTAAAATAAAATATTTAAATAATAAAGA
GAAAAGTATAAAATGTTTGAAAGTATTTATAGCTAATTTCTATTCTATTAATAAAA
TATATCAAATTAAAAATATTATAGAGAGTATTACCAGAGATGTTACAGAAGATTT
GAACCTTCTTCCTTCTCTCCCCACCAAACTACTCGATACTCATGTGGAGGATTTAA
ATTATTGGGACTTAACGTTCCTGCAGACTAAAGGCCTAAGACTTGATGCGCACAA
AGAAGCACCCCACATTGGAGAGGTTGAAATTCTCACACTATCACGGTTAGGATTC
GAATCTTGACTCTTAGTACCAACTCTACAGTTCCAAAACCAGTTGAGCCACGTCCA
TCAGAACCACAATTAATAATTTTTTAAAAAAATAATAAAAGAAGTAACAACATAT
AACTAGCTTTTTTTTCTTGACAACCAAATATAACTAGCTTTGAAAATAAACATTAG
TTCTTTTACTTATCAATATATAAAAAAAAATTTTAAAAAATGCTCTTTTTAATTGA

GAAAACAGTGTAAAACTAAAATATTTCTTATTTAGCCAGAATTTAACCCAAGCTT
AATTAAGAATACAACATAAAATCCTGTCACATGTCATAAGCAATGGGGACGAAAT
TATACAGCGAGTATATGGAATGTGATGATATCATATAATAATAATAATAAATAAT
AATAATAATAATGAAAATATTTGGAGGGCCACACCCACATCCCCGCATGACATGC
ATGTCATATATAGAAACCAAACCGTAGAGTATAAATTTGGAGAGAGAAGTCCTTC
CGCTCCTTTCTTCGCCTTCTGCGCTCTCACAAAGGTGGCTACAGCCCAACAATGGA
GCGCAGTAGGCTTCTTTTGTAGCGAATGTATCAAACCTCGGCCCCCTCCCTCCCCA
CCCCAAATAATGGCAATCATTCACTACTGATGTATCTATATATTAGATGCCAGCCA
AACATACTTGGCTACCCCCCACCAGCAGCAGCCTCCTTCATCCATCACTCACTGCC
GGAGCTCAGAAAGAGGTGAGAGTGAGCCTCTCACATCATTATTTTTAATGGATCTT
CTCTACTAGATAGAACAGAGATTGATCTCTTGTTTATAAATTAGTAGAGCTAGCTA
GGCAGCAGGCTGATGATTTATTCCCTTAATTTGTGCAGAAACTTCTTCATTTGCCTT
TGGAAATG
>G8O promoter and 5’UTR
ATCAAACCCTTGCTCTTCCTTATATATAGAGTTTTTTTTTTTTTACAAATTTATGGA
CAATGAATTTTTACAAGGATGAATTTCTATTAATCACTTTTAGATTTATAATTTTAT
AAGTTTTTTTCCACCATATCATTTTTTACTTCCTTTATCATAATTAATTGGGTTCTTT
TTCTAATTAATAGCATCATGACTGTCATATATATATATATATATATATATAGAGAA
AGTTAAGCTCTAAACCCTAGCAAAAAGAAAATAAAATATATTGAGAAGGCAGAA
TTGTGCAAGGAAGCAAAGGAGAAGAAAAAAATAGCAAACTAGATACATGTGCGA
AAGATTAATTAAAGGAATTTGCATTGTATCAATTGTTTGATGAATATTTTTTCTTA
AAATTTGCTATTTAACATTTTCCTTGTACTATTTGATAAAAAAAAATTTTTATTTAG
TAATTGTATTAAGTTTTCTCGTTATAGAATTTCTCGCTTCAAGGATCAAGAGGAAA
GTTTTAATATTTAATTTATCTATCCAAATAAAAATTCAAATAATTTTTTGTTTTTTT
ATTTTGGCTTTTTATTATTTTATGGTTGCAATTTTTTTTATTAAATTAATTGATTTTT
GTTAATTTTCTTTTACATTGTTGTCAAATTATAATTATCTATTCGTATGTAATTTATT
GGATGATGTTATAATTCTAAAAACTTTTAATTATGAAAATATTTCATTAAATTTTA
ATATTTAATTTAAATAATAATATTCATCTATTTAATGACCTCACGTGGGTTGATTG
GTTAGTAATATAAAAAGTAAATAAATATTTTTAAAAAAAATTAGACAAGTAACAT
AAAATATTTTAAGTAACTCGAGCTTCTCGAGTTCTGTCCGACTTGGCCAGTAGCTA
TCAATCTCCATTCTCCAGTAGTAGTATATTAAAGAACACTGCCTCAAATCTCCATA
GCCACAGAGCAAGCAAGGAGAACAAATTGTACACTTCCATTCCAATG
>GO promoter and 5’UTR
CGTAATGAATTTAGTGACGGATCGAAATTGAGGAGTCACTAAATTTTCTGTTTTTA
GTGATTTGGTTGGAGATAGCCTAATAAGCAATCCAAATAGCAAATTATCGTTTCCT
GTCTTTTTTGACTAAATCATGTTTTCCTCTGTCCACTCTATCTATAGGCTATAATAA
ATGCGTTAGAATGAAATTAAATCCCTTTCCATCTAAAAAAAATACAAAAAATCAA
TGTCACTGAGAATGAAATAAGAATCTTCATTTCAATGTTTTCATTTTGTTTGGTAG
GGAAAATTTGAATACGATGGTTATTCTTTCTTTTTTTTTTTTTAGTCCCATTAATAG
TAAACAGATTTTAAAATTCTTTCGAAAGTCTTTGTTCAACAAAAAAATCTTAACTG
AGAAAAAAAATACTTGACTGAACAAACTAGCTCAGCTTGCAATCCACTGTTATGA
GCAGAGAAACTGTGGTACATGGAAAAAACTTTTTTATTCCCCATAGAAATGACCA
ACTAGCTTTTAGATTGAGAAATCAGTAATATTAGTCTTTAAATTATTTTTTTAATTT
TAATTTAATTCTTAATTTTTATTTTAATTTTTTAGTCTTTAAATAATTATTTCTAAAA
ATTTTAATCATTATCATTAATTTCATAATAAAATTTAACAATAAAAATTTGTGAGT
CTCACAAAACTAATGAACAACGTTAAAAAATACATTTTACTTTATTCAACTCATCA

TTACTCTGTTAAACACAATAATGATGGTCTATTTGAATATTTGAAATATTTTTGAG
GTTTATGTGAAATTTTATTTAAATACATAAAAATTTTCTCCAAGAAAATTTCCCTCT
AAATTTTGAGAAATTTCTCTAATTAGATATAGGAAATATTTTTAGGGTTTAAACTT
GTTTTCGGAATTTAGAGAAAAAATCACATAAACTCTAAGAATATTTCAGGTACTC
AATTAGACATCATTGCTTTATTTAAAAGAGTAATGAGTAAGTGAAATAATTTTTTA
ATCATATTCATGATTCAAGTGAGATTCACATGACTTTTACTACTGATTTTTTTATA
AAATTAATGGCGAACATTAAAACTTTTTAAAAATAAATGTGCAGAACTAAAAAAT
GAAAACAAAATTTAAATATTAAGTTAAGATAAAAAATAATTTACACACTAATATT
ATATTATATTGCTTCTTAGAAAATTTTGAACGCATATCTTTTTGGTTTGGGGGTAA
ATTATGATTTCAGGGCAGCCATTTCAATATTAATAAGATAAAATAATGTAATGTAT
TATGTGCATTACACGTGTACTTATTAGTAGAATCATTTAACGTTTTTTCCTCCAAAA
GAGAAAAAGGAATCATTACACGTAGCAGCAGCCCTTATATAAATATATGATTATT
CTTTCGTCATTTACTCGCAACTAGTCAAAATAAGTGGACAATTATAATTTACAAAA
AAGTTAACCTGAAATAGCAGCCAAATCTGAACTAAAACTAAAACAGCAATAAAAT
ATTATGTATTCTCAGCCACCTTTGAGTCGCCCTATTATATATATATAATGAGTGGG
ATGCTAAGCTAAAGAAATCATTCCCCATTTTCCTTCCTTCCTTCCTTCTTCAGTCTC
CTCTCTTGAACAAAACACAATG
>LAMT promoter and 5’UTR
GGAGGCAAAAGAAAGAGAGAAAGAAGAAAAAAGAAATAAGGAGGAAGAAGAA
AGAGAGAAAAAAATGGGCGTGCGACTGAGGCAAAAGAAAGAGAGAAAGAAGAA
AGAGAAAGAAAAAAGAGGCGGCGCGGCTAGGGTTTATAACGTAATAACGCATAG
TCATTGTGTGTTATCACTCATAATAATGGACAATGGACGTGCGTTATCATGAATGA
TAACGCACAATGGCTGTGCGTTATTGCCACATCAGTGCCAGATCATCGATTGATAA
CGCACAATGGCTGTGCATTATCGTCACATCAGCGCCACATCACCGCCACGTTAAA
AATGCTATTTAACGCGGCTTAACGTCTCTTAAGCTGGTGATTAGCGATAACGCACA
ATCATTGTGCGTTATCTGAGGTGATAACGCACAATGACCGTGCGTTATAGCCACTT
TCAAATTTTAAAAAAAATATTGACCTATTTAGTACCAATTTCTTGAAATTAGGCTA
AAGAAAGAAAAAACCTGTACAATACCTAAACGTGCTTCCGTCACCGGAGGTAGTA
AAATTTCAGTTGTAAAAACATAAAAAAATAAAATAAAATGCATAAAAACAGAGA
TAATTAAAAAACAATACATATAGGATGTAGTGAATGATGGAAATTCTTTCTTTGAC
CAAACAAAGCTCATCATATTTGAACGCAAGAAAGTTCTCACTCGGAAGTCCAAGT
GTGAAAATTCTTATATATGTTATAGGCAAATTATTTGCTGTTTTTTTCTCCAAATA
CGTCACAAAAAAAAATTTAATTAATCCTTACTTATGTCTCCAAATATACTTACCTG
ATTTAATATTCCTTCCTTGAAAATATTAAAAATGATTTTAATTTTGACTATTTACAT
TAATGAATAAACATGCATTTATAGTCTATTTCGTACTACTAATTCATGAAGTGAAG
TCAAAAAGATTAATAAAAATGTGAACGTTTAATTTGAAACACATGTTAGATGATA
TGTGTGGCCGCCGCCTGTTGCTCGATTTCGAGACTTCAGTTACAATGGTCAAATAT
GAAACCTGTTAATAATTCATATGAATTATTATTAAAAAATTATTATAAAGAATTA
TTCAATTCATATATGTATCGATGTATGATGCGAATATATTTTTTTTTTACCTCGCA
TTTTTCTGTTTTATTTGCACTTTTATCTGTTCTATTATTATTAAATTGTCACTTGTTT
TATTATTAAATAATTGTCATTTGATTTATTCTTTTTAAGTAGTATTTCTCCTTTTTAT
TATTTTTTAAAAATAAATTAACAAATGAAAATTTGATGGTAATGGAAGTAGATATT
AATATGATTATGTTTTTAATTTAAAATTTTAATTCTTTTATAAAGTAATAAATTTTA
TAAGATTTATAAAATTTATGTTGATATATAAAAAAAATTAATCATACACACTCATT
AACTATTTTGTCCTTTTTATTCTATTAAACTATTTTTTCAAAAGTTTATAATTATAA
ATTATTTACAATTTTACATATATATCAAATTATACTTAATACAAAAGACTTCCAAA

TATATTAAGCCCGTAATTTATATATAACTCTTTCTTTATTAGGTTATAACTACACAA
TTAAAATTAAAATAATACATTTAAAAAATACATATCTCTATTATCGTTCTATACAC
ATCTCCTAAAAATAATATATAGAACCCAATTATTCCTTGGTAGGTGATCATATAAT
AAAGGAAGGAGATTGGAAGCCGTGGAGGTTCATTCATGCAGTAAAAAAAAAAAA
ACCTATCTAACCATGTGCTGATTCATGTGGCTGCTAAGTAATAAAAATAAAAGAA
AATTCTTAGAAGAAAAAAAAAAGAAAATTAGAACATTTATTGCCTATATAACCAC
TTCAGTTTGCACTCACTTCTTCATCTCTCTCTTCACAACAAAAACAGGAGAGAGAT
TTCTTATTCAACAAAAGCTCAAAAAACTCATACAGAGAAGAAAGAAAAATG
>MYC2a promoter and 5’UTR
ATAGCATATAATTACTCTCTAGTTGTCTCGACTTATATTATTAAACAATGATATTTC
CATATGACATCATTTATTCTTTTATTATCTCGACTAATAATTTTAGACAATGATATT
TCTATCGCTATAGAATAAGTATTCTCTAGTTAAATTCGAATCACATTTGATCCATG
GCCTATAATATTTTTTTTATGTAACCAAATCACCCTTATAAAGAAAGTTGAAAAAA
AAAATCTTTCTAACGGGTTTAAATAAGTTGTTACTAATGTTTTAATAAGGATAATA
TGAATAAATACTTTTTTTTATATTGATATTTCTAAAATTGTTTAGATAAAATCATTA
TCTAATCTTGAACATTAATAATCATATTACATTTATCAACTTTCCATTCTCGCCTAT
CTATTATATTTACCAGACCAATGAGATCAAGTCATAACAATAATCTTAGTTTTATG
GCCAAGAATTCATCAAATTCTTTAATCAAACAACAAATATTATCTATAACCAAGCT
CCAACAATTAAGATCTAAAAACAAATAATTTAGAAGATTCATTTTCCAATTTAATT
GATAAAACATAGTAATATCTAAAAACTTAAACAAAGATTATTTAGACATCAATGG
AGATGGACCCAAACATTAATAATCAAGATTCATTTGATAGTTCAAATATTCACTTA
TCGATTGGACTTATATTCACTCATAATTTTAAGAGAATCAATAATAAAAATCTCAT
AACTCATGAAACTAATAAAGATTTTTTTTTAAAGAAGTTTAACTAATACAATAACA
TAAAAGATCTAAATACAATAAGATAAAAACGTATTTATAATTTATACATAATAGA
AAAGAATAGGGGTGTAAAATTAGAATTTATACAAAGTTCAAGGACCTAAATTATT
AATAGGGTGAAAAGAGCGCCGGCTTTAAGTTAAGAGAAGAGAAGAGAAGAGTAG
ACGCCAAGACGATTAAATCGGCGGGGGGGTTCGTGGAACCCGCGGTTCTTTGTCC
ACTCTCCTCAAATTCCATAAACTCTAAATTCAGCTCAAATTCATCTCAAATAAATC
CCAAACCTAAAATCAAAAATGCATCCAACAATTTCTCACAAATCCCTCGCCCTATC
ACCACTTTTCTCTCCCCACCACTAGCCGCCCTCCCATTTCTCTCTCTCTCTTCCTTCA
ATTTTTCACTCACTCAAACAGACTTCTTCTTCTCGAATTCAACAATCACTGCCATAC
CCAGAAACAAGAACCCTTTTTGATTCTTCAATTTCTTTTTTTTTTTTTGTTTGATGG
AATG
>ORCA3 promoter and 5’UTR
GGAGTGTCCCAAGTAAATAAAAACGGAGAGTAAAAACTATACTNNTGCTTAGGA
GCAGAAAAAAATAAATGTTGGATAAGATGAACGAATCAATTATTATGTAAATGTG
GTGGCTAGTTTCAAAAAAATACTTTATCATCATCACGGTATGGACTTCGTCAGTAT
TCATCAATGCCAACTTTTTCTTTCCGTCCAAAATGTATAATTGTTTTAAAATACACT
ATCTAAACATTTAAGTCATATTTTTTTACGTGAGTAAGGATATTTCTGTCTCACGTC
ATGCGTAATTTCATCATTGTCTGTTGAATACTTTTATTCGTGTTAGAGGGTGGCTCA
AATGTGTAGATGACTGTATTTATATTAATCTTTGATATTTCTTTTATATTTTTATCTC
AAAATTGATTGAATCGTGTATATAATCTTCAAAATTATTTTTTTTTAACTTCAAAA
AATTTTCAAATTTTAAAAAATTACCACATTTTTTGGAAAGAAAGTAAAAACAACA
TGTATAGAGACAGAAATTTTTTTCTGATATAAATTTTATATTCTTTTACGTAATTTA
TCGGCCCAAAATAATTTACTTAAGAAATACGTGTTCCATAATTATTTAATTCTGTG
TAAACGTGACTAATTGCACCTCCCAAGCGCCATCGCTGGACCTTCAAGGAACACT

CTACAAATTATTTTTCAATAAAATATTTATCCATTTTTCAAACGTGCCTTTTTACCT
CTTAGTATATAAATTCCACTCCTCACTCTATTCATCACTCCAAATCCAAAAAACTT
CATAAAAAATCAGTTTCTAAAAAAGAAGAAAAATG
>STR1 promoter and 5’UTR
GATCTTTATTGTTTGTATTTTCATAATTAATTAATCCCTACAACTGATTTCAAAAAA
AAATCTATTTGAAAAAAATAATTCTGCTACTAAAAATTTTAAAATCTTCCTATAAT
ATCATTTTACGAGTTTAATTTATTTATTATCATATTAAAACAATAATATTATTCATT
ACAGAAATTGGACAAACAAAAAGATCTAGCAAAAGTATTATGATTTTAATATATA
GTCGAAAATACGAATTAGACAAGCGAAACATTTTGTAATTAAATCGCTATTTATC
ATATAATTATTTTACAATAATTAGTATTAGGAATTATAATTATTAATTGATGTATA
AACCGATAAAAATATATTTATGATATATACAAAATAAAGGAATAAAAGAATGCAT
AAGTATATTGGATAAGCCTATAAATATTACTCCACGTGGTACATCACTCTTAGACC
GCCTTCTTTGAAAGTGATTTCCCTTGGACCTTGTTTGGTGAGTTTGTGGGCTTATTA
TATTCCTTCACTGTCCCTCTAAGTATCAAACCATACCCCATAGCTGCCCCTCTCTGC
TTCACTCCCATCATTTACAGTCAACAATG
>T16H1 promoter and 5’UTR
CCGGGTAGGCAGTTTGATTGAAACATCAGTCTGAGAAATCTCCTACAAAAAAGAG
TTTTGTTTTCAATATTTTCAAGAAATGTTCAATTACTCTCCCTCACCTTTTTTATTAT
TATTATTTTTTTTTTTATTTGTCTAACACATGGATAAAACATACATACTAAATATGG
ATTTTAGATGTTCTGTATCACTTTGTGTACCATTCTTTTTATCTGTATCACATTTTAA
TTTCATATATTTTATTAATAAATAAAAGATAAATTCTAAGAAAAAAATAAACTCTA
AAAAAAATATGTTATAAAAAGAATGTGACATATAGAGTAATATATAAAGTGGTAC
ATAAAATCTTGTGCCTTAAATATGAATTAAATATGAATAGAACATAGTCTATATAA
TATTTTAAGTATAATAAAGTTTTTAAAATTTAATAATTAATGTAAATTATTGAATTT
TAAGAACATTTATACTATATTAAATTATAAAAGTAATCCACATTATGTATTCCCTT
CGATATACGGTTAGACAAACTTTAACATACAATTATTAAGAAAATAATTAATTTAT
AGACTAAATTGTTATTATTTAAAAATATGAGTTTATATGAGTCATGTTTAAATATT
CTATAAGATCGTCTATTTAAATAATAAAAAGTAATAACATGTGTGAAAAAATTTTC
ATTTATTAATCTTTTAATTTTTAAAGTAACAAGTATTTTAAAATTGAAAAATATTTT
AAAGTGAAAAAAATATATTCCAACCATAAATAAATTGAATGGAGGAAATATAATT
AAAAATATGTTAACAGACAATAAAAGCACGGTTCCCACGTGAGGAAAAAATCGC
AAAACACAATCAAAGCACGGCTAGAAGCAGAATTTGCGTCTTTCCATTGATTTCC
ATTAAAGATTTTTTCATCGAGTCCAAAAACATGATAATTAATTGTTGAAAAGTTCG
TTTGTCTTTCCCAAGAGAGTGAAAGGCAATCAAAATAAATTTAATTTTTTTAAAAT
AAAAAATGAGAAACATATTAAAATTATAAATTTTTTAATCATTTTTTGCTAAATAT
TAAAAAGTTTATATTTTCTTACTTCTTTTTTATTAAAAATTCAACAGATATTTAAAG
AAATTATTTTTGATGTAGTGTTAATGATGCTTATATATAAGTTTCTAAATATGATTT
GATCAATAACATTTCATTCATTATCGGTAAAAATAATCAAAAATTACTTCTAAATC
TTTTAAAAAAAATAATTATTATGTCATAAGCATTAAATGAACTTTGTAAATCATTCA
TATAATTTTTATTTTTTTAAAAAATACATAACCAAAACATAAATTAAATACTCAAA
TATTAAAATATGAAATATAGTATATAATTTACAATAATTTAAGCTTTAGAGATAAT
ATTTTTTATACTAAAAATATTTAATTACTTTAATTAAAACTTAATTATCACATTAAA
AAATTCATATACCACCCAAACTACCAACCACTAGAGAAGAAACACATCCTCCATA
AGTCAAAAAATAATTGATGTTAGAAAAAAATTAAAAATTGCCAAAAAAGCAACA
TATGGACCAAGACAAGTGAAGGAATTATAGAGAATCCCCTATAAAAGCCATGCAT
CGTAAGGGCTTTTATCCTAAAGCAAATCAATAACAAGCATAATTTTCACTAATG

>T16H2 promoter and 5’UTR
AGTAATCTCATTGATGTGATGGGCGGTGGTATGGTAATGCTATTTTTAATTTGATA
TATTTTATTAATAAAATTTAAGGTTAATATAAAAAAAGTACCTTGAACTTTGTCCA
TTATGTCAATTAAGCACCTAATCTTTGAAAAAGAAAATTAAGGAATAATTTTGCTT
TTTCGATAAAATAAATTGAACGAAAACTGATCAGAAATGAGTAATGTATGTTATG
CCCTAATAAAATTTGGAAAACTTAAATTTATCCTTTTATAGGAGGAAAGCTTGTCA
TCAAAATGGAGGGAAGAAATATAATTAAAAAAAAATTCTCTTCCAAGACCATCAA
AATACCCAATTACTACCAAATCCTATCAGCAGAAACAAATGGCATGAGTTTCTTTC
TCTTCCTTCCCCAGACCCTTAAAAACCATATACCAAAGCTTTCACTTCCTTCTCCAA
ATCCTCAAAAAGCATATAAAAGTTTCATAACACCATTATTGTGACTTAAAAAACC
ATTGTTTTGCTTGCAATCAACTTTAATGGGTAAGAAATATTATAGAGATTAAAAAG
TTGTAACTTCTAAAATATATGAAATTATCAAAAAGATATTCTGGGGAACGATTAAT
ACAAAAAAATTACCATGAACTTTGTCCATTGTCAGATTATTCTTTTTGACAATATC
ACAATGTTTTTTATTATATTTTTGATGACCAGCTCCCCTCCTATGCAAGGATAAAA
GTTGGTTTTCCAAATTTTATTAGGGTAGAACAGAAACTACTCATTTCTGATCAGTT
TCCGTTCAATGCAGTTTACTAGAAAAGAAAAATTATTCTCTTGTTATTTTGAAAGA
TTGAATACTTAATTGATATAATGACAAAATTCAAGATACTGCTTTTGTATTAACAG
TAAAATTTATCAAAAGAAAAAAATTCTTAAGGGAAAAGATAAAAATTAAAAAAA
TATATATTATAAAAGAAATAGTATAAAGAATAGTATAGAAAAATGGTAGAGATGA
TCCAGAGTGGTGGGAGCCTAGAAATACAAGTAATTCTATGGGGATCAATATCAAT
CAAACTAATTGTGTGTTAGAATAGGATAAAAATGTTTATTTATGGAAAAACTACTT
ATATCCATAGAAGGAAAAAACTATAATTCTCTCTCTCTCTCTTTTTTTTCCCCCTAA
TTTTACTCTTTTTGAGAAATCTATGATTCTCGTGTGGAAAATAAAGGTGTTTATTTC
TCAATAAAAGCCATTCCATGTTTAAGCAAATAAACAACAGCCACACAGCTGAAAG
TATCTTTCTTCAAATCTATAAAAGCCATTCCATCTTACTAGTGTTAAGCAAACTAC
ACAGCAAAAATCTTGATTTTGGTCAACTTTCTCTACCTAATG
>T19H promoter and 5’UTR
CCATGAGGTAGAAATTATGAAAGAAATTAAGGTAATAAATTGATTTTTCTTTTATA
CCTTTTCCTAATATTGCATGAATGGGTACCGGCTCTAAAAAATGGATGTAGTTTGT
ACATATTAATCTTAGCATATATTATATTTAATTATAGAGTATCGTCGGAAGGCGCT
TCCAAACTGTAGGATCCGTAGGAGGAAGCCGTAGATGATGACGGACATATGGTCG
ATGCGTAAGATGAAGCGGCAGCATCGGAACGTGCCAGTAACTGGTGGCTGTGTTG
GTGGATCTTGATCTGTACGTATCTATGGATCCTGCTATACTTGAGTCAATGAATTT
TACTATGTCGTAGGCTGTGACTTCTACTGTGTCGAGGGCGTTAGATCCTTTCTCTTA
CGTCCTCGAGCCATCAAACCTTTAATATAAGTAAAAGGATAGTATTTGTTACGCCT
CTAATTCACATACCCACACAAACTCAACTGCGTCTAATAGTCATTGGTAATTTATT
TACCGACGGCGTTCATCATCGGTAAAATGTCATCACTTTCTGGCACACCCTTTCCT
TCTTTTTTTAGTTGGAAATGTGAAAATCCATTGACATACAAGTTTTCATCGGTACC
CCATCAGTGTTTAGTAACGATAAACTGTAATGTTAGTAAAGTCCGTCGGTAACTGT
TTATTTATATATATATATATATATAATATGCATAATTGATCACGCGGCTTTATATG
AATGGACGATCCGTTTTCTCCGCTATGTGTTCGTTGGGCAATCTTCTTCTTAGCATG
CCTTTCATTGACTCATTGGTGTTTCTTTAGCTATTAAATTAGTATTAAATTAAACTA
TTCCAATGCTTTAAAATGAAATGACATAAAAAAATCAGTATAATAATACATCAAT
GATCATGTAAATCTGGACTTTTCTGAACAAATCCATCTATAAATGATGAGAATATC
TAAAGAAGAGAGCCATTCATAGCAAAAACAACTAGATTTCTCTAGCAAGAAAAAT
AATG

>ZCT1 promoter and 5’UTR
TGTCTCATTTTCGGAGTTTAGCTTGCAAATTGAGCATTCATGACCCCTATGCTCCA
ACTTGAGGGGAAAGAGGGGGAGGGGTATTGGCGAGAATATCTCCATGTTTCCTTA
CCATTCTTAGGGTGGTTCATTGTTGTCAATTGGCACATATTGGCACATATTATGCG
TAGGATTACTGCGATTAATTAGGATATTACCTAATGTAATTTTAGCTTAATTTCCT
AATTTATCTACGTGTAAGATGTATATAACTTTGAAAGGATGATCAATGAAAGATA
GAGAAAGCATGTTAGTTTTTCCGCTGAAAACTTTCAATAAAAATAATCCATGAGT
ATAAATAAATAAACTTTAATATTTTAAATGTAATGTAATTTAGAAAAATAGTTAAA
TTACATTTTAAAAAAATTTGACAACGAGATAGTAGTATGATGATAAGACTCTTCTC
TTTCAAATATCTCTGCAAGATAAATGAGAAATTTTAAGTTGTAATTATTTTAGACG
AGATTTTTATTAAATATTTAGCCTTATAATATAAATGCAAGTCTTGTACACCAAAT
TCTTTGTTAGAGTAGTCAAAATCTATGGGATAAAAAAGTTAAAAAAAAGCAAATA
GATAGTAGGTCGGTTTGTCTTGTGGCGCGTAGCTTCACATAATCTGTTGATTAAAA
GATGAGAAGGTTCCTAGTTGAGGAAGAGTCAAACAAGGTTCACGCCTTTCTTATG
AATCCTTGTCTATCTTCCCTCCCTTACTCATCTTGTTTTTCCACCGTGAGCAGTGAC
GATTATCTACACAAACCCACACGTCAATATCACCTCTCTCTCTGCTTTCTCTTTCTT
AGTCCACAAGCAAAATCTTACTCGTACTTTCCCACTCCCACCTGTATATATACCCT
CAATCTCTCTACTAATTCCTCACTCACAGCTTTCACAATTCCACCTAATTAAAATTA
ATTATCTAATCAATTCCCAAGTCGCAAGTTTCAATTCAATAACAAAAAACAATG
213
